# Nuclear N-Glycosylation Redefines the Glycoscape and Directs Cell Identity

**DOI:** 10.64898/2025.12.17.694357

**Authors:** Fenjie Li, Xingui Wu, Yixuan Xie, Jiafeng Yang, Jiaojiao Xi, Yuetong Sun, Jiazhang Chen, Xiao Wu, Shenwei Huang, Chi Zhang, Yi Zheng, Wenwen Li, Xiaoyu Zuo, Yang Li, Kaining Chen, Xingqiang Lai, Xingyu Liu, Wei Zhong, Jixiao Zeng, Qiang Wu, Benjamin A. Garcia, Haojie Lu, Huimin Xia, Wenhao Zhou, Yan Zhang

## Abstract

Glycans modify nucleic acids, proteins and lipids in fundamental biological processes, but have long been considered confined to the secretory pathway and cell surface. Here, we challenge this view with evidence that N-glycosylation, a ubiquitous glycan modification, occurs within cell nuclei across tissues and species, exhibiting cell type-dependent abundance and localization patterns. Using a multi-faceted approach, we show that numerous classical membrane and secreted glycoproteins localize to the nucleus with N-glycan modifications, exemplified by the cell adhesion molecule L1CAM. Mechanistically, we show that the N-glycosylated L1CAM is transported to the nucleus by KPNB1, where it transcriptionally modulates neuronal differentiation. Moreover, this nuclear N-glycosylation event contributes to neurodevelopmental disorders and is pharmacologically reversible. These findings revise the cellular geography of N-glycosylation and expand the known routes of protein trafficking, and highlight nuclear N-glycosylation pathways as potential therapeutic entry points for developmental disorders.

## INTRODUCTION

Glycans are ubiquitous and structurally complex biomolecules that modulate nucleic acids, proteins, and lipids across all domains of life^[1–3]^. Unlike information-encoding nucleic acids or proteins, glycans are often highly branched and heterogeneous, which makes them exceptionally difficult to structurally analyze, synthesize, enrich, or detect^[1,2,4–6]^. As a result, glycan remains one of the least understood molecules, leaving major dimensions of the cellular glycosylation landscape unexplored.

N-glycosylation, a widespread glycan modification, plays vital roles in diverse biological processes including development, oncogenesis, and immune responses^[5,7–10]^. The canonical view in cell biology has long been that N-glycans are strictly limited to proteins (and glycosylated lipids) that transit through the secretory pathway or reside at the cell surface^[11–16]^. Indeed, the enzymatic machinery for N-glycan assembly is sequestered in the endoplasmic reticulum (ER) and Golgi lumen, any presence of N-glycosylation in the nucleus has long been deemed implausible^[17]^. Even the recent discovery that small noncoding RNAs can carry N-linked glycans on the cell surface (so-called glycoRNAs) broadened the paradigm of what types of molecules can be glycosylated, yet still did not challenge the dogma that N-glycans reside exclusively outside the nucleus^[18–23]^.

Despite the prevailing paradigm, a handful of intriguing observations over the years have hinted that nuclear proteins might carry N-glycans^[24–28]^. As far back as the late 1970s and 1980s, certain nuclear and chromatin-associated proteins were reported to bind glycan-specific lectins and incorporate radiolabeled sugars in a manner suggestive of N-glycosylations^[24]^. However, because definitive structural characterization of the glycans was lacking and such results were difficult to reproduce in intact cells, these claims were largely dismissed as experimental artifacts or contamination from the secretory pathway^[28]^. Thus, whether N-glycosylated protein occurs in the cell nucleus remains an open question. Technical barriers to detecting and validating glycosylation in specific subcellular compartments have so far precluded a definitive answer.

Motivated by this knowledge gap, we set out to directly test for the existence of nuclear N-glycosylation using a multi-pronged strategy that circumvents previous technical obstacles. Recent advances in glycan detection and labeling technologies enabled us to probe this question with unprecedented sensitivity^[18,29,30]^. By combining lectin-based fluorescence imaging and metabolic glycan labeling with azide-functionalized sugars, we visualized N-glycosylation patterns in intact cells. To confirm the glycan nature of these signals, we enzymatically removed N-glycans with PNGase F (PNGF) and blocked new N-glycan addition by either pharmacological inhibition of the oligosaccharyltransferase (OST) complex (with the inhibitor NGI-1) or targeted OST subunit knockdowns. In parallel, we enriched nuclear glycoproteins and performed glycoproteomic analyses to identify N-glycosylated proteins in the nucleus. Using this multifaceted approach, we discovered unequivocal evidence that N-glycans are present within cell nuclei.

Our findings reveal the nucleus as a previously unrecognized locale for N-glycosylation. We observed robust nuclear N-glycosylation signals across multiple species, tissues, and cell lineages, definitively overturning the long-held assumption that N-glycans are restricted to secretory compartments. Moreover, we discovered that N-glycoproteins can enter the nucleus via the importin KPNB1 and directly influence transcriptional programs governing cell-fate decisions, identifying a mechanistic link between nuclear N-glycosylation and gene regulation. This insight suggests that nuclear glycoproteins could serve as potential therapeutic targets in certain developmental disorders. In summary, this unexpected discovery expands the known geography of protein glycosylation and marks a conceptual shift in our understanding of the cellular “glycoscape”.

### Systematic Validation of N-glycosylation in Nucleus

We systematically investigated the presence of N-glycoproteins within the nuclear compartment by employing complementary methodologies. Specifically, we employed lectin-based detection and click chemistry-mediated metabolic labeling in hiPSCs (human induced pluripotent stem cells) and HEK293T (293T) cells (**Figure 1A**). For lectin-based detection, we used a panel of lectins: Concanavalin A (ConA) for general N-glycosylation, Pisum sativum agglutinin (PSA), Aleuria aurantia lectin (AAL), and Ulex europaeus agglutinin I (UEA I) for fucosylation, and Maackia amurensis lectin II (MAL II) and Sambucus nigra agglutinin (SNA) for sialylation. In parallel, for metabolic labeling, we utilized azide-tagged probes including UDP-GlcNAz and Ac_4_GlcNAz for general glycan incorporation, GDP-FucAz for fucosylation, and ManNAz for labeling the sialylation precursor ManNAc. To validate that any nuclear signal indeed represented N-glycans, we enzymatically removed N-glycans with PNGF, blocked N-glycan addition onto nascent proteins by knocking down key OST complex components (STT3A, STT3B, DDOST), or by pharmacological OST inhibition with NGI-1 (**Figure 1A**).

**Figure 1.**
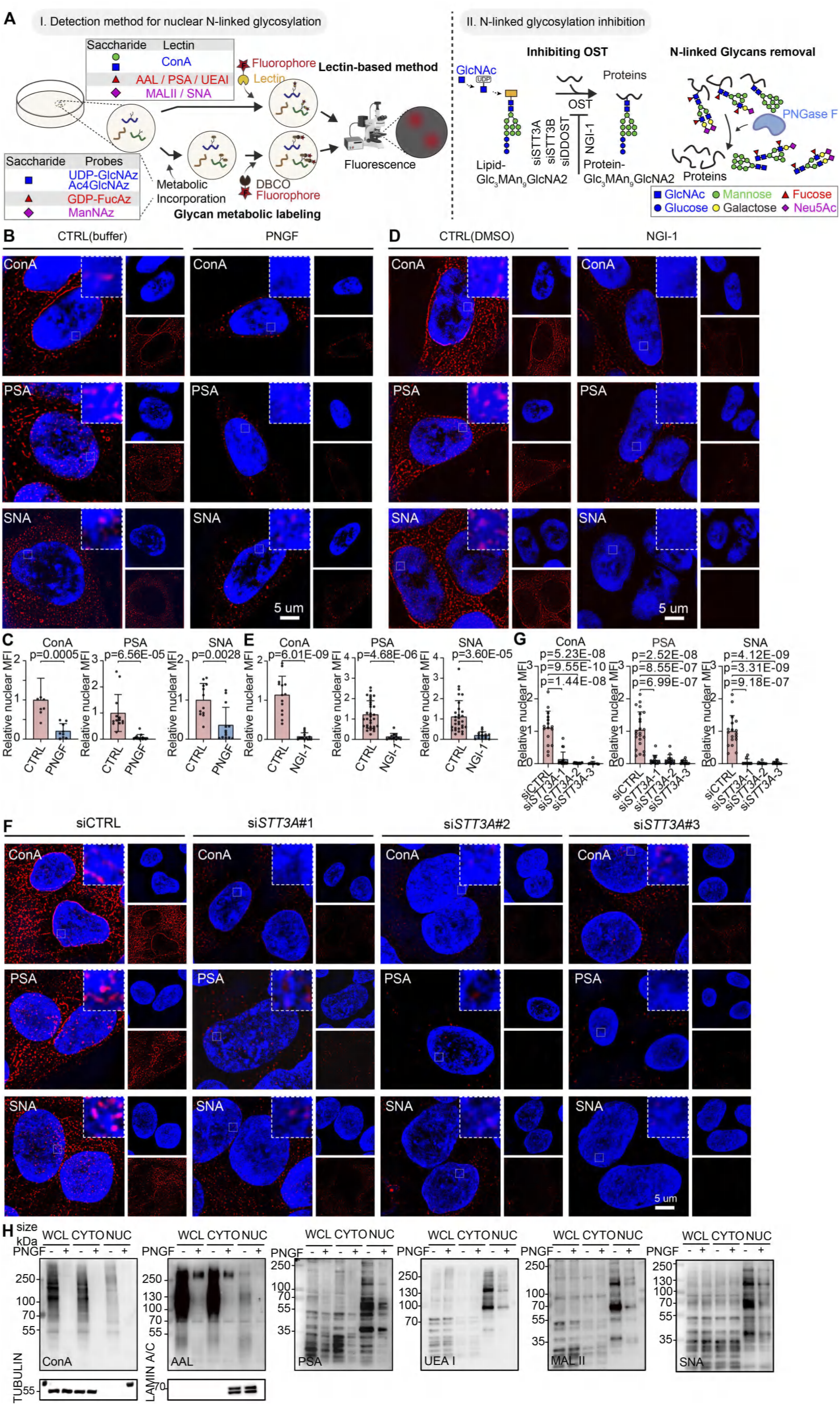
Systematic validation of nuclear N-linked glycosylation. **(A)** Schematic diagram illustrating the experimental workflow for precise detection of nuclear N-linked glycosylation using complementary approaches, including lectin-based recognition and metabolic labeling via click chemistry. Specific lectins employed are ConA for N-linked glycosylation; PSA, AAL, and UEA I for fucosylation; and SNA and MAL II for sialylation. Metabolic labeling utilized azide-tagged substrates (UDP-GlcNAz and Ac_4_GlcNAz for N-acetylglucosamine incorporation; GDP-FucAz for fucosylation; ManNAz for labeling the sialylation precursor ManNAc). To verify glycosylation specificity, cells were treated with PNGase F (PNGF) to enzymatically remove N-linked glycans, siRNA-mediated knockdown of the OST complex subunit *STT3A,STT3B, DDOST*(si*STT3A,*si*STT3B,*si*DDOST*), or pharmacological inhibition of OST activity with NGI-1. Images made using biorender.com. (**B-F**) Super-resolution fluorescence imaging of hiPSCs stained with indicated lectins (ConA, PSA, SNA, shown in red) recognizing nuclear N-linked glycosylation. Nuclei stained with DAPI (blue). Cells were subjected to PNGF (**B**), NGI-1 (**D**), or si*STT3A*(**F**), treatment as indicated. CTRL, control. Scale bars, 5 μm. Statistical analysis was assessed by t-test (two tailed) for PNGF (**C**), NGI-1(**E**), or si*STT3A*(**G**), treatment. MFI, mean fluoresence intensity. **(H)** Lectin blot analysis of whole-cell lysates (WCL), cytoplasmic (CYTO), and nuclear protein (NUC) fractions from hiPSCs treated with PNGF. TUBULIN and LAMIN A/C served as cytoplasmic and nuclear markers, respectively. Additional details are shown in **fig. S1-9.**

Super-resolution fluorescence microscopy, combined with lectin staining and the metabolic labels, revealed clear nuclear signals corresponding to N-glycans in both hiPSCs and 293T cells. Critically, these nuclear N-glycosylation signals were significantly diminished following PNGF treatment, NGI-1 inhibition, or OST knockdown in both cell types (**Figure 1B-G and Figure S1-9**). This loss of nuclear staining upon removing or blocking N-glycans confirms that the observed nuclear signal indeed represents N-glycosylation.

Among the lectins tested, PSA specifically recognizes core-fucosylation, a hallmark of N-glycan modification that is catalyzed exclusively by the fucosyltransferase 8 (FUT8). We further verified that PSA is indeed detecting core-fucosylated N-glycosylation in the nucleus. siRNA-mediated knockdown of FUT8, as well as pharmacological FUT8 inhibition (using the inhibitors FDW or SGN) both caused a marked loss of nuclear PSA staining (**Figure S10**). This result validates that the nuclear PSA signal specifically reflects core-fucosylated glycans.

We additionally corroborated the presence of nuclear N-glycoproteins via lectin blotting and immunoblotting analysis on isolated nuclear fractions. Untreated nuclear extracts from hiPSCs showed strong N-glycosylation signals that were nearly abolished after PNGF-mediated deglycosylation (**Figure 1H and Figure S2D**). Collectively, these results provide compelling evidence that N-glycoproteins are widely present in the nuclear compartment.

### Broad Distribution of Nuclear N-glycosylation Across Tissues and Species

Having established nuclear N-glycosylation in cultured cells, we next examined its presence across diverse tissues and species. Using the panel of lectins described above, we systematically profiled nuclear glycosylation in multiple human tissues. Among these, PSA lectin, which specifically recognizes core-fucosylated glycans, revealed robust nuclear staining signals across various human tissues, including brain, colon, liver, and kidney (**Figure 2A**). Likewise, PSA-detectable nuclear N-glycosylation was observed in multiple vertebrate species such as mouse, rat, pig, chicken, and monkey, indicating that nuclear core-fucosylation is broadly distributed across different organisms (**Figure 2B-F**).

**Figure 2.**
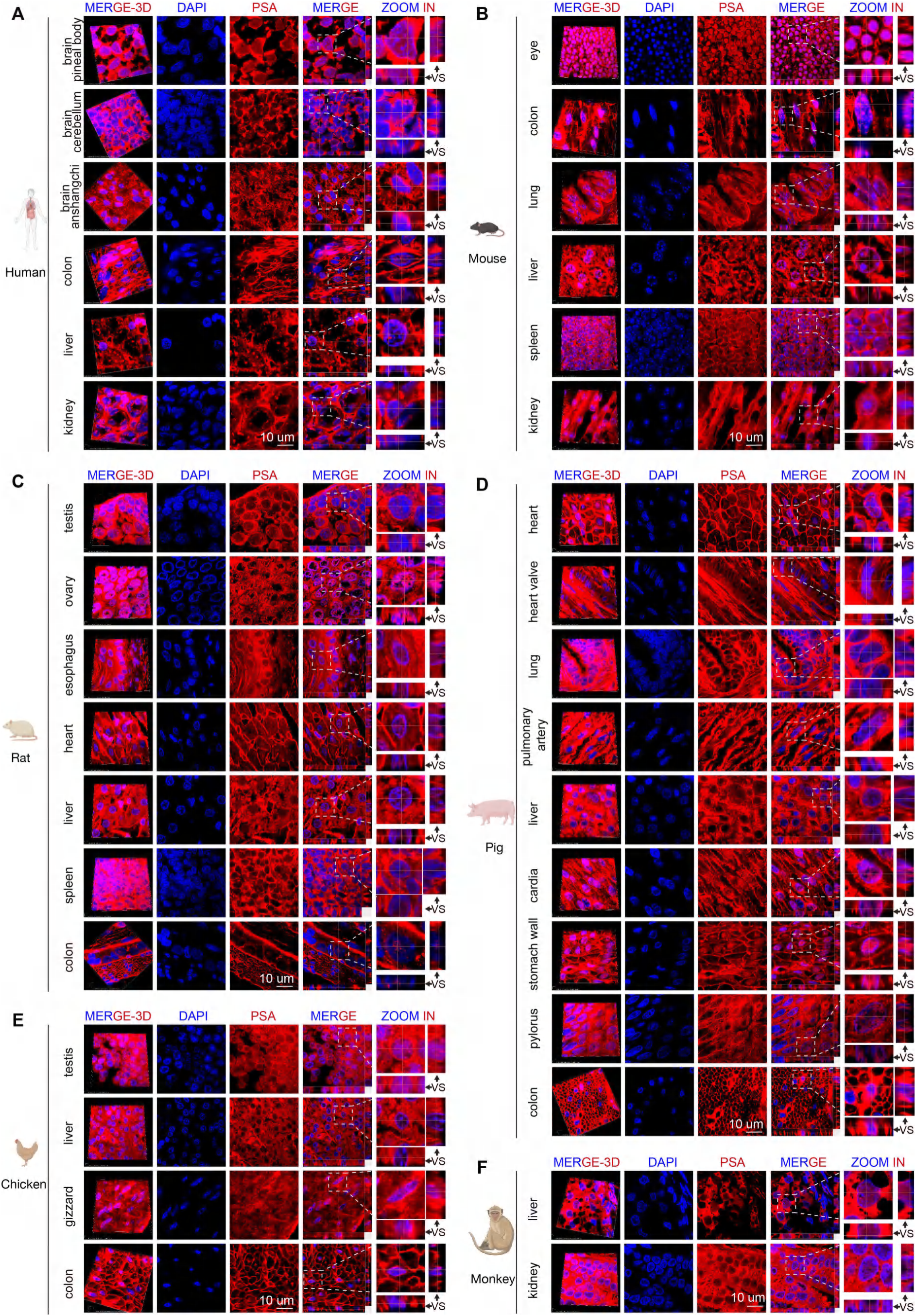
Broad distribution of nuclear glycans across tissues and species. **(A-F)** Fluorescence images of tissue sections from human (**A**), mouse (**B**), rat (**C**), pig (**D**), chicken (**E**), and monkey (**F**) stained with PSA (red) show clear nuclear glycan signals across species. Nuclei are counterstained with DAPI (blue). Images are representative of three independent experiments. VS, vertical section; scale bars, 10 μm. Additional details shown for other lectins are in **fig. S11-15.**

Other lectins also detected nuclear glycosylation in a broad range of tissues and species, albeit with variable intensities and localization patterns (**Figure S11-15**). For example, ConA, AAL, UEA I, as well as MAL II and SNA, all showed nuclear signals in certain tissues, though the strength and distribution of these signals differed between tissue types. This suggests that various nuclear glycan modifications may play distinct functional roles in different biological contexts. Overall, our results demonstrate that nuclear N-glycosylation is widespread across tissues and species, with distinct abundance and localization patterns depending on the biological context.

### Distinct Nuclear Glycan Profiles Characterize Stem and Differentiated Lineages

Because our tissue survey suggested context-specific differences in nuclear glycosylation, we next asked how these patterns vary with cell lineage or differentiation state. Core fucosylation is a hallmark modification of N-glycans, and the lectin PSA exhibits high specificity for the core-fucose epitope, as reported previously^[31]^ and independently validated in our experiments as above. We found that hiPSCs and their neural derivatives (enteric neural crest cells, neural progenitor cells) exhibit bright PSA staining in nuclei, whereas mesodermal derivatives such as osteoblasts, chondrocytes, and adipocytes display only faint nuclear PSA signals (**Figure 3A-C**). Consistently, PSA lectin affinity enrichment of nuclear proteins followed by immunoblotting confirmed an abundance of core-fucosylated nuclear proteins in pluripotent stem cells and neural lineage cells (**Figure S16**).

**Figure 3.**
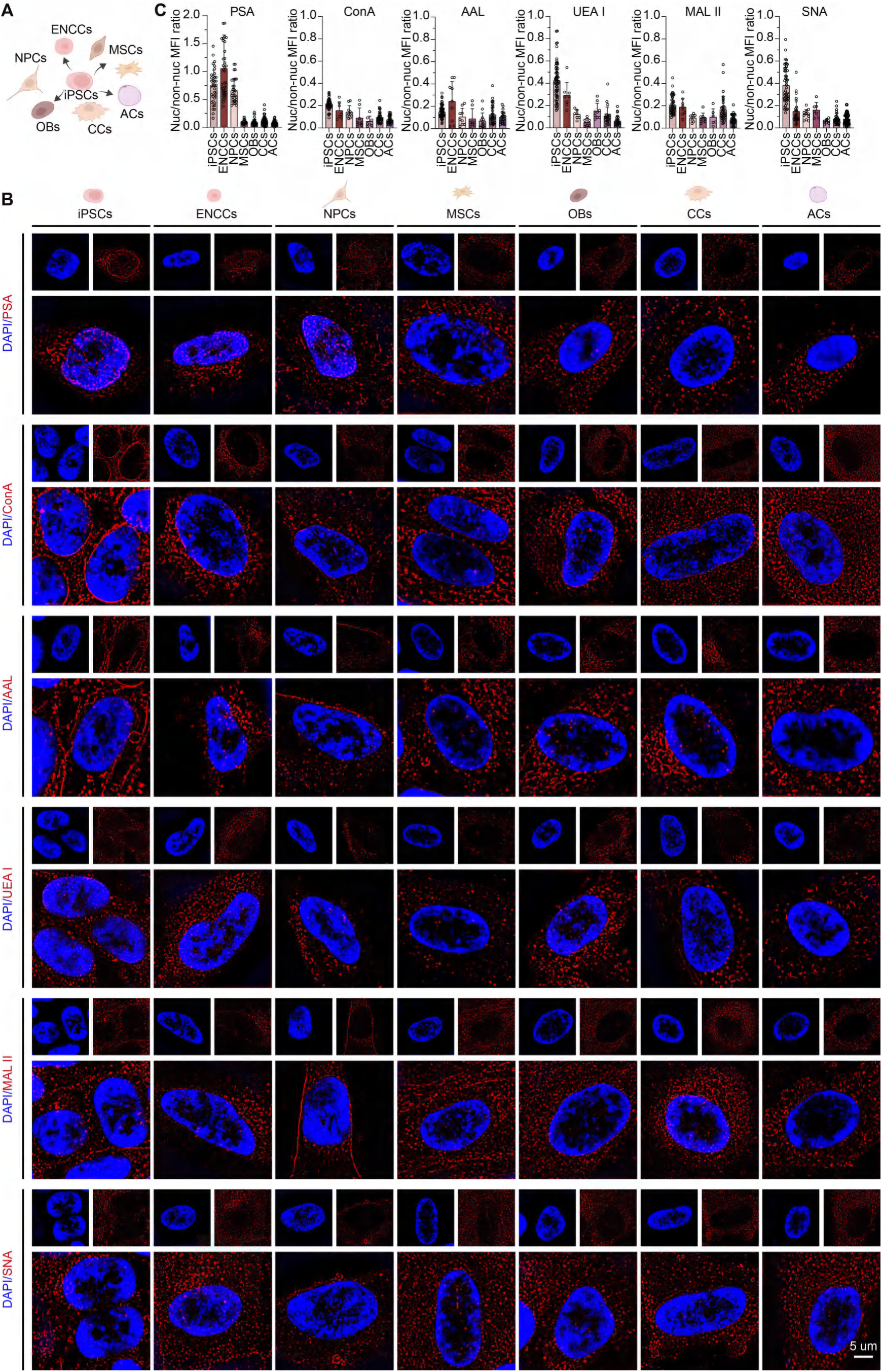
Distinct nuclear glycan profiles characterize stem and differentiated lineages. **(A)** Schematic of cell differentiation from hiPSCs into various lineages. Images made using biorender.com. **(B)** Super-resolution fluorescence imaging of nuclear glycosylation (lectin, red) during differentiation of hiPSCs into various lineages including enteric neural crest cells (ENCCs), neural progenitor cells (NPCs), mesenchymal stem cells (MSCs), osteoblasts (OBs), chondrocytes (CCs), and adipocytes (ACs). Nuclei stained with DAPI (blue). Scale bars, 5 μm. Quantification was analyzed by t-test (two tailed). (**C**) Quantification analysis of nuclear/non-nuclear intensity of lectins. MFI, mean fluoresence intensity.

Complementary staining with other lectins further revealed distinct nuclear glycosylation enrichment patterns across lineages (**Figure 3B and C**). For instance, ConA yielded uniformly low nuclear signals, with particularly weak staining in the nuclei of terminally differentiated osteoblasts, chondrocytes, and adipocytes. In contrast, the fucose-binding lectins AAL and UEA I, as well as the sialic-acid-binding MAL II, showed moderately higher nuclear staining in hiPSCs, enteric neural crest cells (ENCCs), and chondrocytes than in other cell types. These signals remained low or negligible in mesenchymal stem cells and mesodermal derivatives. Similarly, SNA binding in the nucleus was much stronger in hiPSCs relative to other lineages.

These patterns indicate that different cell types harbor unique profiles of nuclear glycosylation, with pluripotent and neural cells being especially enriched for certain modifications (notably PSA recognized core-fucosylation). Collectively, our findings demonstrate that nuclear N-glycosylation is a widespread feature across species, tissues, and developmental stages. Among the diverse nuclear glycosylation, PSA recognized core-fucosylation stands out for its pronounced enrichment in the nuclei of pluripotent stem cells and neuronal lineage cells. This prominent nuclear core-fucosylation in pluripotent states and neural contexts suggests a critical role in orchestrating cell differentiation and neurodevelopmental processes, warranting further investigation into its functional significance.

### Nuclear Import of Core-fucosylated L1CAM Facilitated by KPNB1

To identify nuclear proteins bearing core-fucosylation and explore their functional significance, we developed a novel PSA lectin-affinity enrichment combined with mass spectrometry (PSA-MS) approach. We used biotin-labeled PSA to enrich core-fucosylated glycoproteins, which demonstrated superior specificity compared to the agarose method used in previous literature (data not shown). This first-of-its-kind method allowed us to enrich and identify a series of core-fucosylated nuclear proteins from ENCCs (**Figure 4A**). We unexpectedly found that numerous proteins traditionally localized to the cell membrane or secretory pathway appeared in the nuclear fucosylated proteome. Further functional enrichment analysis showed that these nuclear fucosylated proteins are significantly enriched in neurodevelopment-related pathways (such as neuron differentiation, neuron generation, and neurogenesis) as well as in cell migration-related pathways (including cell adhesion and cellular motility). The identified proteins included many classic membrane and secreted proteins, for example, L1CAM, which has been reported to be N-glycosylated. Among these nuclear N-glycosylated proteins, we found that L1CAM is enriched in most of the neurodevelopment-related pathways and cell migration-related pathways, suggesting that nuclear N-glycosylated L1CAM may play critial roles for neurodevelopment and cell migration. Fluorescence imaging further corroborated clear nuclear co-localization of PSA lectin and L1CAM in hiPSCs, ENCCs, and enteric neurons confirming the nuclear presence of core-fucosylated L1CAM in neural lineage specification from pluripotency (**Figure 4B**).

**Figure 4.**
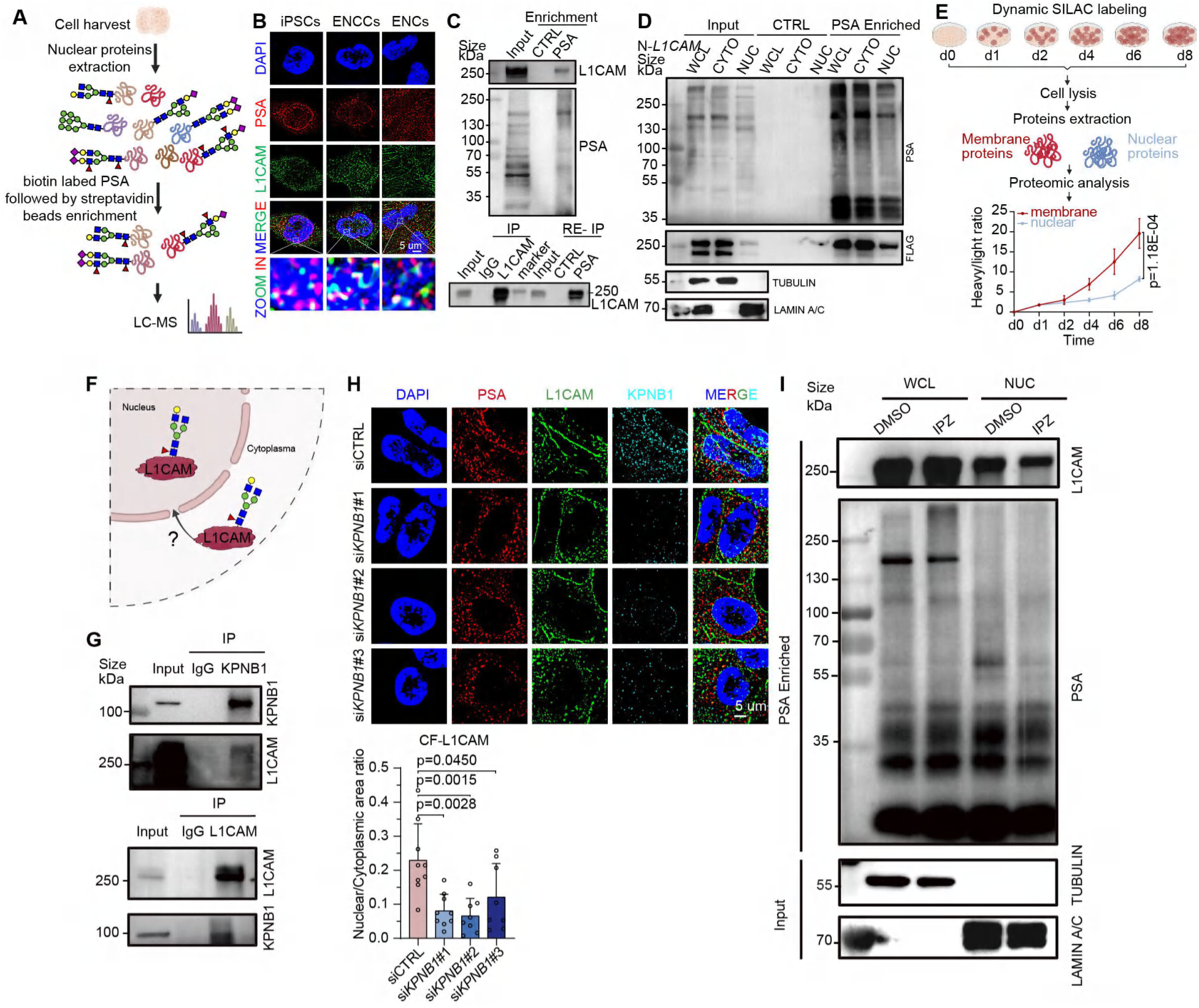
Nuclear Import of Core-Fucosylated L1CAM Facilitated by KPNB1. **(A)** Schematic of nuclear protein enrichment using biotin labeled PSA lectin affinity and subsequent mass spectrometry (MS) identification of nuclear core-fucosylated glycoproteins in enteric neural crest cells (ENCCs). Images made using biorender.com. **(B)** Fluorescence confirming nuclear colocalization of PSA (red) and L1CAM (green) in hiPSCs, ENCCs, and ENs; nuclei stained with DAPI (blue). Scale bar, 5 μm. **(C)** Tandem immunoprecipitation (IP) validating nuclear core-fucosylated L1CAM: anti-L1CAM IP followed by PSA lectin enrichment (CF-L1CAM IP). IgG and buffer served as negative controls; input represents 1% of total protein. **(D)** Immunoprecipitation of FLAG-tagged L1CAM from nuclear fractions followed by PSA lectin affinity confirms nuclear core-fucosylation of full-length L1CAM. TUBULIN and LAMIN A/C served as cytoplasmic and nuclear markers, respectively; IgG and buffer as negative controls. Input represents 1% of total protein. **(E)** SILAC proteomic analysis of proteins extracted from membrane or nucleus, respectively. Membrane L1CAM exhibited significantly faster turnover than nuclear L1CAM, supporting L1CAM in membrane is biochemically distinct from that of nuclear L1CAM. d, day. Quantification was analyzed by linear mixed model using R package “nlme”. **(F)** A model illustrating how core-fucosylated L1CAM is transported to the nucleus. Images made using biorender.com. **(G)** Co-IP assays demonstrating interaction between L1CAM and nuclear import protein KPNB1. IgG served as negative control. Input represents 1% of total protein. **(H)** Fluorescence analysis showing reduced nuclear localization of PSA (red) and L1CAM (green) upon siRNA-mediated *KPNB1*knockdown in hiPSCs; nuclei stained with DAPI (blue). Scale bar, 5 μm. Quantification was analyzed by t-test (two tailed). **(I)** Co-IP using PSA lectin affinity confirming reduced nuclear core-fucosylated L1CAM upon KPNB1 inhibition by importazole (IPZ); DMSO as control. IgG and buffer as negative controls; input represents 1% of total protein.

Earlier studies reported the nuclear localization of truncated L1CAM fragments (28- and 70-kDa intracellular domains, termed L1-28 and L1-70), but did not examine their glycosylation status^[32–39]^ (**Figure S17A**). To determine whether the nuclear core-fucosylated L1CAM detected here represents full-length or truncated forms, we performed PSA enrichment, followed by western blotting. This analysis consistently revealed a dominant band of approximately 250 kDa, indicating that the nuclear L1CAM is full-length and core-fucosylated (**Figure 4C**). Then we conducted tandem co-immunoprecipitation (co-IP), first enriching L1CAM from nuclear fractions and subsequently isolating core-fucosylated proteins using PSA lectin affinity (CF-L1CAM). This two-step approach also identified a prominent protein band of approximately 250 kDa consistent with full-length L1CAM modified by core-fucosylation in the nucleus (**Figure 4C**).

To further validate these findings, we expressed full-length *L1CAM*constructs at either the N-terminus (FLAG-tag) or the C-terminus (EGFP-APEX tag). Both N-tagged and C-tagged L1CAM (N-*L1CAM*, C-*L1CAM*) localized to the nucleus and were recognized by PSA, confirming that nuclear L1CAM includes the full-length protein bearing core-fucosylation (**Figure 4D and Figure S17B-I**). To exclude membrane contamination as an explanation for nuclear signals, we quantified protein turnover separately for membrane-resident and nuclear L1CAM pools using dynamic stable isotope labeling by amino acids in cell culture (SILAC). Membrane L1CAM exhibited significantly faster turnover than nuclear L1CAM, arguing against potential carry-over from the plasma membrane and supporting the existence of a biochemically distinct nuclear pool (**Figure 4E and Table S2**). Consistent with full-length identity, our MS detected non-contiguous peptides from amino acid 45 to 963, well beyond the known cleavage site at residue 814, supporting the nuclear presence of full-length L1CAM forms (**Figure S17J and Table S3**). Moreover, using the PSA enrichment approach, we observed nuclear localization of full-length, core-fucosylated L1CAM in additional cell lines (293T, MDA-MB-231 breast cancer cells, NCM460 colon cells, and SW480 colon cancer cells), indicating that nuclear localization of N-glycosylation and core-fucosylated L1CAM is ubiquitous across diverse cell types (**Figure S17K**).

We next explore the mechanism underlying the nuclear entry of full-length, core-fucosylated L1CAM (**Figure 4F**). Given previous evidence of interactions between truncated L1CAM variants and the nuclear transport protein KPNB1^[38]^, we hypothesized that KPNB1 facilitates nuclear import of full-length, core-fucosylated L1CAM. Indeed, co-IP confirmed the interaction between full-length L1CAM and KPNB1 (**Figure 4G and Figure S18, A and B**). Critically, siRNA-mediated depletion or pharmacological inhibition of KPNB1 with importazole (IPZ) substantially reduced nuclear levels of core-fucosylated L1CAM without altering total cellular *L1CAM*expression (**Figure 4H and J and Figure S18, C and D**). Concordantly, PSA lectin enrichment of core-fucosylated species were diminished upon IPZ treatment **(Figure 4I).** Thus, KPNB1 mediates active nuclear import of core-fucosylated L1CAM and, more generally, other core-fucosylated proteins.

Together, the PSA-MS method we developed enabled the first systematic identification of nuclear fucosylated proteins. These findings further reveal the nuclear functional role of core-fucosylated full-length L1CAM, implicating it as a novel potential regulator of chromatin remodeling, transcriptional control, and developmental processes.

### L1CAM Core-fucosylation at N979 Orchestrates Enteric Neuron Differentiation through PDE11A-cAMP Signaling

To define the functional implications of site-specific L1CAM core-fucosylation, we conducted comprehensive glycoproteomic profiling in nuclear proteins of ENCCs and identified specific core-fucosylation modifications at residues N671, N931 and N979 (**Figure 5A and Figure S19 and Table S5**). Prior work has implicated N979-core fucosylation in the metastasis of hepatocellular carcinoma^[40]^, motivating a focused analysis of this site. We generated *L1CAM* mutants lacking N979 glycosylation (substituting Asn-979 with Ala or Gln) in both N-terminal FLAG-tagged and C-terminal EGFP-APEX-tagged *L1CAM* constructs (designated N-*979A/Q*; C-*979A/Q*). Overexpression of wild-type *L1CAM*(N-*L1CAM*or C-*L1CAM*) markedly impaired ENCCs migration and neuronal differentiation, characterized by reduced neurite outgrowth and decreased expression of neuronal markers (CHAT, TUBB3, PRPH, UCHL1, GDNF). In contrast, the N979 glycosylation-deficient mutants (N-*979A/Q*, C-*979A/Q*) failed to recapitulate these defects; their overexpression failed to impede migration or differentiation, essentially restoring these processes relative to cells overexpressing wild-type L1CAM (**Figure 5B and Figure S20-23**). These results pinpoint core-fucosylation at N979 as a specific and necessary determinant of the L1CAM-dependent differentiation phenotype.

**Figure 5.**
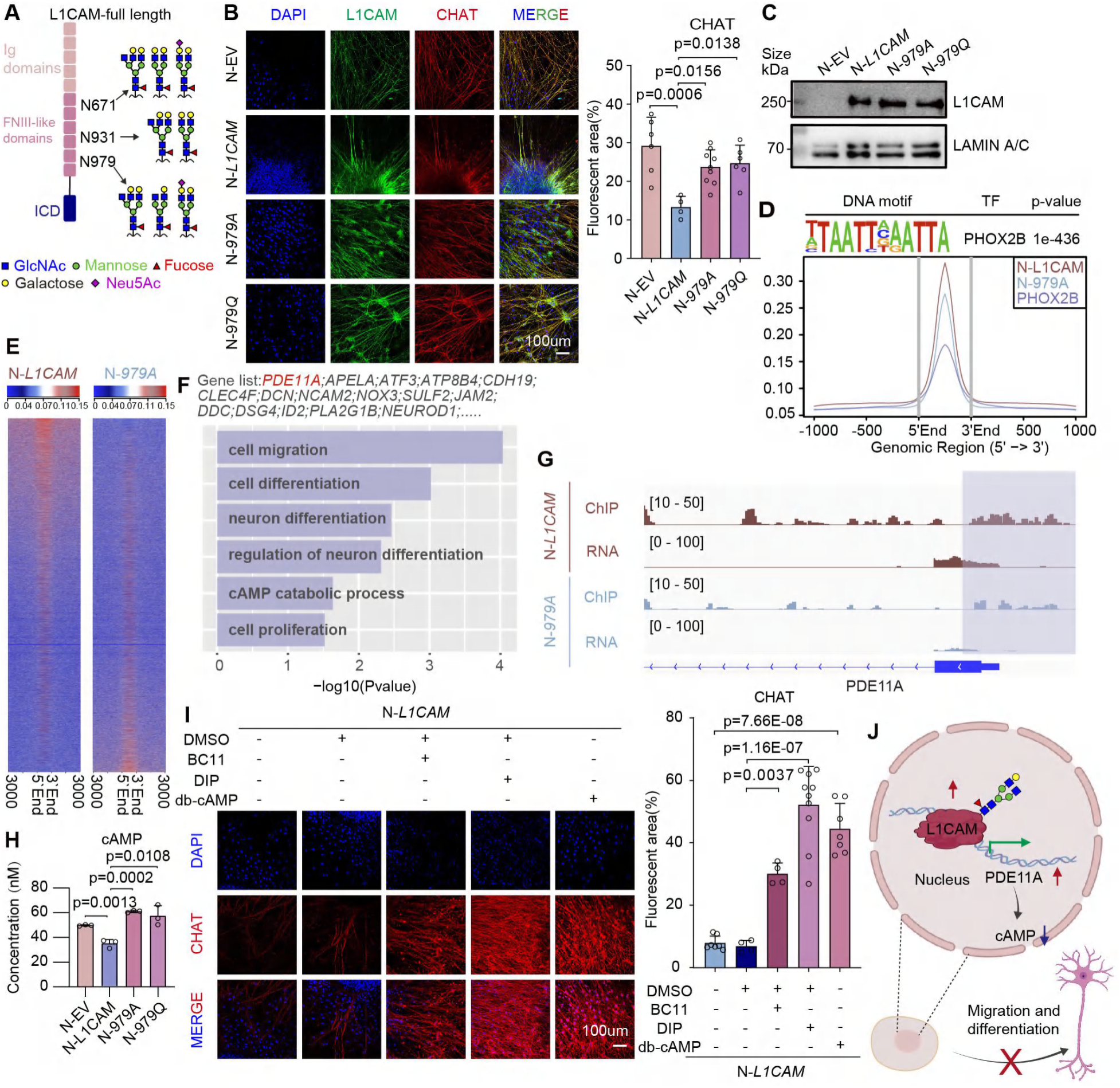
L1CAM core-fucosylation at N979 orchestrates enteric neuron differentiation through PDE11A-cAMP signaling. **(A)** Mass spectrometry (MS) chromatograms identifying N-linked core-fucosylation at residue N671, N931, N979 of L1CAM. Additional sites (N294, N671) detailed in **fig. S20**. (**B**) Immunofluorescence assessing effects of wild-type (N-*L1CAM*) and glycosylation-deficient (N-*979A*/N-*979Q*) L1CAM mutants on differentiation markers (CHAT, red) in enteric neurons (ENs). Nuclei stained with DAPI (blue). Scale bar, 100 μm. Quantification was analyzed by t-test (two tailed). (**C**) Western blot analysis of nuclear protein extracts from ENCCs expressing wild-type or mutant L1CAM variants. (**D**) DNA motif enrichment at genomic loci co-bound by core-fucosylated L1CAM and PHOX2B transcription factor. (**E**) Heatmap depicting ChIP-seq signal intensity at overlapping and distinct binding sites for core-fucosylated L1CAM and PHOX2B. (**F**) Gene ontology (GO) enrichment analysis of genes significantly differentially regulated by wild-type L1CAM (N-*L1CAM*) compared to glycosylation-deficient mutants (N-*979A*) in ENCCs. (**G**) Representative ChIP-seq tracks of FLAG-tagged L1CAM occupancy at the *PDE11A* genomic locus in wild-type (N-*L1CAM*) versus mutant (N-*979A*) expressing ENCCs. (**H**) Intracellular cAMP concentrations in ENCCs expressing empty vector (EV), wild-type (N-*L1CAM*), and glycosylation-deficient mutants (N-*979A/Q*). Statistical analysis via t-test (two tailed). (**I**) Immunofluorescence demonstrating rescue of neuronal differentiation (CHAT, red) deficits induced by *L1CAM*overexpression following treatment with PDE11A inhibitors (DIP, BC11) or cAMP analog (db-cAMP). Nuclei stained with DAPI (blue). Scale bar, 100 μm. Quantification was analyzed by t-test (two tailed). (**J**) Model summarizing N979 core-fucosylation on L1CAM as a critical regulatory mechanism regulating PDE11A-cAMP signaling and enteric neuronal differentiation. Images made using biorender.com.

To understand the mechanistic basis of N979 core-fucosylation-dependent phenotype, we compared nuclear localization and chromatin-binding profiles between wild-type and N979 glycosylation-deficient L1CAM. Fractionation and chromatin-binding assays showed comparable nuclear accumulation and global chromatin association for wild-type and mutant L1CAM (**Figure 5C and Figure S24A and B**), indicating that N979 core-fucosylation does not broadly affect nuclear entry or overall chromatin tethering. Instead, genome-wide ChIP-seq analyses revealed distinct occupancy profiles at regulatory genomic loci (**Figure S24C and Table S6**). Specifically, wild-type core-fucosylated L1CAM uniquely co-occupied genomic sites with the autonomic neurodevelopmental regulator PHOX2B, prominently at the *PDE11A* promoter (**Figure 5D-G and Figure S24D and Table S6-8**). Consistent with locus-selective recruitment, *PDE11A* expression increased in the presence of core-fucosylated L1CAM, accompanied by reduced intracellular cAMP (**Figure 5H and Figure S24E and F**), given the role of PDE11A in cAMP catabolism^[41,42]^. Importantly, pharmacological PDE11A inhibition (BC11-38, BC11; dipyridamole, DIP) or direct cAMP restoration (db-cAMP treatment) rescued the migration and differentiation defects elicited by wild-type *L1CAM*overexpression (**Figure 5I and Figure S25 and S26)**.

Together, these results define N979 core-fucosylation of L1CAM as a crucial regulatory determinant of enteric neuronal differentiation through selective regulation of PDE11A-cAMP signaling (**Figure 5J**). This represents a previously unappreciated N-glycosylation-dependent transcriptional control mechanism governing cell fate.

### FUT8-L1CAM Core-fucosylation Drives Hirschsprung’s Disease and is A Therapeutic Target

L1CAM is an established susceptibility gene for Hirschsprung’s disease (HSCR)^[43–45]^, an intestine motility disorder characterized by disrupted migration and differentiation of ENCCs, resulting in aganglionic intestinal segments^[46–48]^. Building upon our earlier observations of elevated nuclear core-fucosylation in ENCCs and the identification of L1CAM as a prominent nuclear core-fucosylated protein, we investigated whether aberrant nuclear accumulation of core-fucosylated L1CAM contributes to HSCR pathology.

Indeed, we observed significantly elevated core-fucosylated L1CAM in aganglionic (HA) segments compared to ganglionic (HG) segments from HSCR patients (**Figure 6A**). Correspondingly, FUT8, the sole enzyme catalyzing core-fucosylation, was markedly upregulated in HA segments, coinciding with increased nuclear localization and enhanced core-fucosylation of L1CAM (**Figure 6A-C**). Direct interaction between FUT8 and L1CAM was further validated using co-IP assays (**Figure S27**). These findings strongly suggest a critical role for FUT8-mediated nuclear core-fucosylation of L1CAM in HSCR pathogenesis.

**Figure 6.**
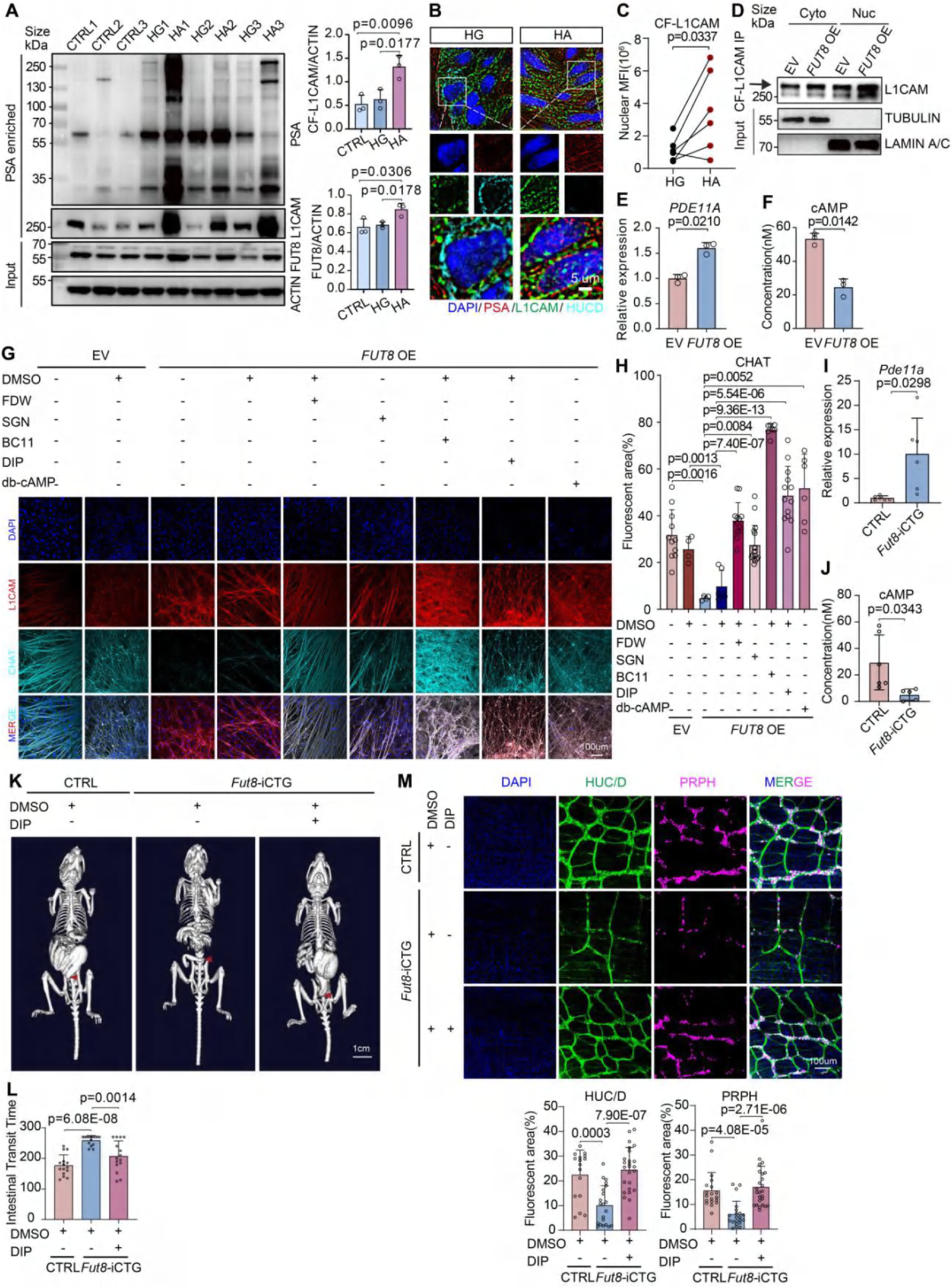
Targeting the FUT8-L1CAM core-fucosylation axis ameliorates neuronal differentiation defects in Hirschsprung’s disease. (**A**) Co-immunoprecipitation (co-IP) enrichment of core-fucosylated proteins by PSA lectin affinity in ganglionic (HG) versus aganglionic (HA) colon segments from HSCR patients as well as control samples (CTRL). L1CAM detected by immunoblot; buffer as negative control, input represents 1%. CF-L1CAM, core-fucosylated L1CAM. (**B** and **C**) Fluorescence illustrating increased nuclear core-fucosylation (PSA, red) and L1CAM (green) in HA segments compared to HG segments from HSCR patients. Enteric neurons identified by HuC/D (light blue); nuclei stained with DAPI (blue). Scale bar, 5 μm. Quantification was analyzed by t-test (two tailed) (**C**). MFI, mean fluorescence intensity. (**D**) Tandem co-IP (L1CAM IP followed by PSA enrichment, CF-L1CAM) validating enhanced nuclear core-fucosylation of L1CAM upon *FUT8*overexpression (*FUT8-*OE). IgG and buffer as negative controls, input represents 1%. (**E**) RT-qPCR analysis of *PDE11A*expression in ENCCs with *FUT8*overexpression (*FUT8*-OE) versus control (EV). Statistical significance assessed by t-test (two tailed). (**F**) Intracellular cAMP measurements by ELISA in EV and *FUT8*-OE ENCCs. Statistical significance assessed by t-test (two tailed). (**G** and **H**) Immunofluorescence analysis demonstrating impaired neuronal differentiation (CHAT, light blue) induced by *FUT8* overexpression, rescued by pharmacological FUT8 inhibitors (FDW, SGN), PDE11A inhibitors (DIP, BC11), or direct cAMP supplementation (db-cAMP). Nuclei stained with DAPI (blue). Scale bar, 100 μm. Quantification was analyzed by t-test (two tailed) (**H**). (**I** and **J**) Mouse model validation: RT-qPCR analysis of *Pde11a* (**I**) and intracellular cAMP measurements (**J**) in *Fut8*-overexpressing (*Fut8*-iCTG) mice compared to controls (CTRL). Statistical analysis via t-test (two tailed). (**K**) Micro-CT imaging of mouse to assess intestinal motility, showing that *Fut8*overexpression inhibits intestinal motility, which can be rescued by PDE11A inhibitors (DIP). The red triangle denotes the distal extent of the barium within the intestinal tract. Scale bar, 1 cm. (**L**) Carmine gavage and red feces excretion of mouse experiment evaluating intestinal motility defects induced by *Fut8* overexpression, rescued by PDE11A inhibitors (DIP). Statistical significance via t-test (two tailed). (**M**) Immunofluorescence (HuC/D, red; DAPI, blue) assessing enteric neuronal differentiation defects induced by *Fut8*overexpression in mouse, rescued by PDE11A inhibitors (DIP). Scale bar, 100 μm. Quantitative analysis shown in panel, statistical significance via t-test (two tailed).

To directly test the functional significance of this pathway, we experimentally perturbed FUT8 in ENCC models. Overexpression of *FUT8* in ENCCs led to elevated core fucosylation of nuclear L1CAM and recapitulated an HSCR-like phenotype. In particular, *FUT8*-overexpressing ENCCs exhibited increased *PDE11A* expression, lowered cAMP levels, and a failure to properly differentiate into neurons (**Figure 6D-H and Figure S28-29**). Neuronal marker expression and neurite development were severely reduced by *FUT8* overexpression, phenocopying the defects caused by wild-type *L1CAM* overexpression. Crucially, we found that these differentiation impairments could be rescued by targeting either the glycosylation enzyme or the downstream signaling. Treatment of *FUT8*-overexpressing ENCCs with FUT8 inhibitors (e.g., FDW or SGN) sharply decreased L1CAM core-fucosylation and restored ENCC differentiation (**Figure 6G,H and Figure S28-29**). In parallel, inhibition of PDE11A with dipyridamole (a clinically approved phosphodiesterase inhibitor) or supplementation of cAMP also fully rescued the neuronal differentiation of *FUT8*-overexpressing ENCCs (**Figure 6G, H**). These interventions validate that it is the FUT8-L1CAM-PDE11A axis that drives the observed differentiation defects.

We extended these findings *in vivo* using zebrafish and mouse models. In zebrafish, overexpressing the *fut8a*gene during embryogenesis resulted in heightened expression of *pde11a*, reduced basal cAMP levels, and impaired development of the enteric nervous system (**Figure S30**). The gut of *fut8a*-overexpressing larvae showed severe depletion of enteric neurons, mirroring an HSCR-like phenotype (aganglionosis). In agreement with our cell-based results, treating the *fut8a*-overexpressing fish with either FUT8 inhibitors (SGN, FDW) or PDE11A inhibitors (BC11, DIP), or supplementing cAMP, fully rescued the formation of enteric neurons in the larval gut (**Figure S30**). Similarly, we generated an inducible transgenic model overexpressing *Fut8* in enteric neural crest cells (i*Sox10*:*Fut8*-OE mice, *Fut8-*iCTG). Upon inducing *Fut8*overexpression in embryos, these mice showed heightened expression of *Pde11a*, reduced cAMP levels, severely impaired intestinal motility and a reduction of enteric neurons (**Figure 6I-M and Figure S31**), recapitulating an HSCR-like aganglionosis phenotype. In agreement with our cell-based and zebrafish results, treating the *Fut8-*iCTG mice with PDE11A inhibitor DIP fully rescued the formation of enteric neurons (**Figure 6I-M and Figure S32**).

Thus, across diverse models, we show that a hyperactive FUT8-L1CAM axis impairs neural differentiation by inducing excessive PDE11A and depleting cAMP. Importantly, this pathogenic cascade is therapeutically actionable at multiple points, offering avenues to restore normal development. Directly targeting L1CAM itself is challenging given its essential roles and broad expression^[37,49]^. By contrast, targeting FUT8 or the downstream cAMP pathway offers more feasible strategies. FUT8 is an attractive drug target because it is the single enzyme responsible for core fucosylation, and inhibitors of FUT8 are already being explored in cancer therapy^[50]^. Meanwhile, modulating cAMP levels via phosphodiesterase inhibition is a well-established clinical approach, and dipyridamole (used here) is a pediatric-safe drug that increases cAMP by blocking its breakdown^[51]^. Our findings demonstrate that both strategies, inhibiting the aberrant glycosylation at its source or counteracting its signaling consequences, can effectively rescue enteric neuron development in experimental models. This suggests a promising translational avenue for treating HSCR by targeting the FUT8-L1CAM-PDE11A-cAMP pathway.

## DISCUSSION

Our study provides compelling evidence that addresses a longstanding controversy surrounding nuclear N-glycosylation by providing rigorous biochemical and structural validation of its existence. Traditional views confined N-glycosylation strictly to the secretory pathway, fostering skepticism regarding previous observations of nuclear-localized glycoproteins due to methodological uncertainties. By leveraging advanced glycoproteomics, metabolic labeling, stringent biochemical validations, and high-resolution imaging, we definitively establish nuclear N-glycosylation as a widespread, authentic biological phenomenon across diverse tissues, species, and cellular lineages.

The discovery of nuclear N-glycosylation aligns with a growing body of evidence that is reshaping classical paradigms of molecular trafficking and localization. For example, glycoRNAs on the cell surface and glycosylated RNA-binding proteins defy traditional compartmentalization of glycosylation^[18–23]^. Likewise, certain ER-associated proteins can influence nuclear function via unconventional routes, such as the cytosolic deglycosylation and relocalization of NFE2L1 by NGLY1^[52]^. Our work adds a new dimension to nuclear trafficking of glycoproteins. We show that a fully N-glycosylated, membrane-bound protein (L1CAM) enters the nucleus via the classical KPNB1-mediated import pathway. This expands the recognized cargo repertoire of classical nuclear import to intact N-glycoproteins. This mechanism is distinct from the NGLY1/NFE2L1 route, as L1CAM retains its N-glycans en route to the nucleus. These examples highlight that cells possess underappreciated flexibility in trafficking glycosylated macromolecules between compartments.

Functionally, our data reveal that nuclear N-glycosylation, particularly core fucosylation, can serve as a regulatory mechanism for gene expression and cell fate. We found that nuclear core-fucosylation is especially enriched in pluripotent stem cells and neural lineage cells, suggesting a link to developmental potency and neurodevelopment. Indeed, among the nuclear core-fucosylated proteins, L1CAM plays a direct role in controlling transcriptional programs for neuron differentiation. Nuclear, core-fucosylated L1CAM associates with key transcriptional and chromatin-modulating proteins and can tether to specific gene promoters. Through a single glycosylation site (N979), L1CAM is able to modulate the expression of *PDE11A*, altering cAMP signaling and thereby influencing whether neural crest cells differentiate properly. This exemplifies how a post-translational glycan modification can act as a molecular switch to redirect a cell’s developmental trajectory. More broadly, it unveils a previously unrecognized crosstalk between the glycosylation machinery and the gene regulatory network in the nucleus.

Clinically, our findings have important translational implications, exemplified by our identification of the FUT8-L1CAM core-fucosylation axis as critical in the pathogenesis of Hirschsprung’s disease. We demonstrated effective pharmacological rescue of impaired neuronal differentiation by targeting PDE11A-cAMP signaling using PDE11A inhibitors, including dipyridamole, a clinically used PDE inhibitor ^[51]^. Given the widespread clinical implications of PDE-cAMP signaling modulation across diverse physiological processes such as metabolism, immunity, neuronal regulation, and reproduction, targeting this pathway represents a highly feasible therapeutic strategy^[51,53]^. Furthermore, modulating nuclear core-fucosylated L1CAM levels significantly impacted PDE11A transcription and downstream signaling, supporting the concept that multiple neural functions regulated by L1CAM are at least partially controlled by its nuclear core-fucosylation status. Clinical translation studies are currently underway to leverage these insights into novel, effective therapeutic approaches for HSCR.

Beyond HSCR, our work hints at broader implications of nuclear glycosylation in disease. We showed that N-glycosylation are present in the nucleus of cancer cells, including MDA-MB-231 and SW480, suggesting that nuclear N-glycosylation may catalyze progress in cancer research (**Figure S17K**). Preliminary analyses revealing elevated FUT8 expression in multiple cancers further highlight potential applications in cancer therapeutics, warranting deeper investigation (**Figure S32**).

Overall, this study fundamentally advances our understanding of intracellular glycosylation, reshaping traditional dogmas of protein modification and trafficking. Further exploration into nuclear glycosylation pathways offers exciting prospects for novel therapeutic strategies across developmental and oncological diseases.

### Limitations of Study

Despite these advances, several limitations of our study warrant consideration. Although we profiled multiple lectins to map nuclear glycosylation, only the core-fucosylation-specific PSA lectin signal was functionally validated and mechanistically investigated. While we conclusively demonstrated that full-length, core-fucosylated L1CAM localizes to the nucleus, we did not determine whether the nuclear pool of L1CAM carries out unique functions distinct from or overlapping with its well-characterized membrane-associated roles in neurodevelopment. Moreover, we focused primarily on core-fucosylation, leaving the broader nuclear glycome (e.g. sialylation or other glycan epitopes) underexplored. Finally, key mechanistic questions remain unresolved including when glycosylation is added during protein processing and the precise trafficking route by which a membrane-associated glycoprotein like L1CAM is rerouted to the nucleus. It should also be noted that our observations are limited to the tissues and experimental conditions specifically examined in this study. These limitations should be carefully considered, and addressing them will be an important direction for future investigations.

## ACKNOWLEGDEMENTS

We thank the support of the grant of the National Natural Science Foundation of China (Grant NO.81970450 and Grant NO.82570608 to Yan Zhang, Grant NO.31900519 to Fenjie Li, Grant NO.22434001 to Haojie Lu, Grant NO.22507114 to Yixuan Xie, Grant NO.82200561 to Kaining Chen), Guangdong Basic and Applied Basic Research Foundation (Grant NO.2021A1515220146 to Yan Zhang), Guangzhou Basic Research Plan City School (Institute) Enterprise Joint Funding Project (Grant No. SL2024A03J01 to Yan Zhang), the support of the Guangdong Provincial Key Laboratory of Research in Structural Birth Defect Disease (No: 2019B030301004 to Huimin Xia), Greater Bay Area Institute of Precision Medicine I0036(A) to Yixuan Xie, Guangzhou Municipal Science and Technology Plan Project (Grant NO.202201020596 to Yi Zheng), National Institutes of Health AI118891, HD106051, and CA196539 to Benjamin A. Garcia.

We thank Dr. Ryan Flynn from Boston Children’s Hospital and Harvard University for his constructive suggestions.

We thank the glycomics and proteomics platforms of the Greater Bay Area Institute of Precision Medicine and the Core Facility of Shanghai Medical College, Fudan University, for their support in MS analysis.

We thank Dr. Zhongyong Gou from the Institute of Animal Science, Guangdong Academy of Agricultural Sciences, for pig and chicken samples. The experimental protocol was approved by the Animal Care Committee of the Institute of Animal Science, Guangdong Academy of Agriculture Science, Guangzhou, P. R. China.

We thank Dr. Lihua Huang from Guangzhou Women and Children’s Medical Center, Guangzhou Medical University, Guangzhou, China, for monkey samples. The use and care of animals were approved by the Ethics Committee of Sun Yat-sen University.

We thank Dr. Kaishou Xu from Department of Rehabilitation, Guangzhou Women and Children’s Medical Center, Guangzhou Medical University, for rat samples.

We thank Dr. Yiyue Zhang and Dr. Gaofei Li from the Innovation Centre of Ministry of Education for Development and Diseases, School of Medicine, South China University of Technology, for zebrafish experiment.

## AUTHOR CONTRIBUTIONS

F.L. and Y.Z. designed this project. F.L., Y.X., J.X., J.Y., and Y.S. conceived the experiments. Y.X., C.Z., and Y.L. performed SILAC experiments and glycoproteomics. F.L., J.X., J.Y., Y.S., J.C., X.W., X.Z., S.H., and W.L. performed all the other experiments. X.W. performed bioinformatic analyses. F.L., Y.Z., X.W., J.Y., and Y.S. prepared the original manuscript. Y.X., X.L. and B.A.G. discussed the findings. H.X., W.Z. and Y.Z. supervised the project, interpreted results and wrote the paper. All authors reviewed, edited and approved the final manuscript.

## Materials and Methods

### Ethical statement

The Medical Ethics Committee of Guangzhou Women and Children’s Medical Center approved all human subject protocols. All procedures were conducted in accordance with the Declaration of Helsinki and international ethical guidelines for human research. Written informed consent was obtained from the legal guardians of all pediatric participants.

### Human samples

Human tissue samples were collected from pediatric patients at Guangzhou Women and Children’s Medical Center under Institutional Review Board (IRB)-approved protocols. The implementations were in concordance with the International Ethical Guidelines for Research Involving Human Subjects as stated in the Helsinki Declaration. Informed written consent was obtained from the legal guardians of all participants. In Hirschsprung’s disease patients, the narrowed aganglionic segment was identified intraoperatively by absence of peristalsis and confirmed by frozen-section histology. Adjacent dilated colon (>4 cm diameter) was obtained ∼5 cm proximal to the transition zone^[58]^. Each tissue segment was immediately divided into three parts: one fixed in 4% paraformaldehyde (PFA) (Servicebio, cat. no. G1101) for histology, one snap-frozen in liquid nitrogen for protein extraction, and one preserved in RNAlater solution (Invitrogen, cat. no. AM7020) for RNA extraction. All samples used in this study are listed in **Table S9**.

### Other animal samples

All studies involving mouse and rat were reviewed by Guangzhou Women and Children’s Medical Center, Guangzhou Medical University, Guangzhou, China. All studies involving pig and chicken were reviewed by Institute of Animal Science, Guangdong Academy of Agricultural Sciences, Guangzhou, China. All studies involving monkey were reviewed by Sun Yat-sen University, Guangzhou, China. All tissue samples were approximately 0.5 to 1 cm in diameter. Each tissue segment was immediately fixed in 4% paraformaldehyde (PFA) (Servicebio, cat. no. G1101) for histology.

### Maintenance culture of human induced pluripotent stem cells (hiPSCs)

The hiPSCs were maintained on matrigel-coated dishes in PSCeasyⅡ human pluripotent stem cell medium. The culture medium was replaced daily. Cells were passaged every 4-5 days at ∼80-100% confluency. For passaging, colonies were dissociated with 0.5 mM EDTA in PBS and passaged at approximately a 1:8 ratio into culture dishes pre-coated with matrigel. The ROCK inhibitor Y-27632 (10 μM) was supplemented to the medium on the day of passaging to enhance cell survival^[59,60]^.

### Differentiation of hiPSCs into neural progenitor cells (NPCs)

Neural induction of hiPSCs was performed as previously described^[61]^. Briefly, hiPSC colonies were dissociated and seeded onto matrigel-coated dishes in PSCeasy II medium on day 0. After plating, check the cells and if the cells have reached ∼100% confluence, wash the cells once with PBS and add 1 ml of Neural Induction Medium, which consists of DMEM/F-12 GlutaMAX supplemented with 1% N2, 2% B27, 1 mM L-glutamine, 100 µM 2-mercaptoethanol, 100 µm non-essential amino acids, 10 μM SB431542, 200 ng/mL noggin. Continue to incubate cells and replace the Neural Induction Medium every day. Between days 8 and 12 after plating, a uniform neuroepithelial sheet should appear. Gently dissociate the neuroepithelial sheet into aggregates of 300-500 cells by pipetting slowly up and down three times. Then, centrifuge the aggregates at 160 g for 2 min to form a pellet and discard the supernatant. Resuspend the cells in 10 mL of pre-warmed Neural Maintenance Medium. After centrifugation at 160 g for 2 min, discard the supernatant. Seed the cells into 2 mL of Neural Induction Medium in 35 mm dishes pre-coated with laminin. Following overnight incubation, replace the medium with Neural Maintenance Medium. Confirm successful neural induction by reverse-transcription PCR for the expression of pluripotency genes and neural progenitor cells.

### Differentiation of hiPSCs into Enteric Neural Crest Cells (ENCCs)

hiPSCs were differentiated into ENCCs using a stepwise induction protocol. On day 0, hiPSCs were dissociated and seeded onto a matrigel-coated plate in PSCeasyⅡ human pluripotent stem cell medium supplemented with 10 μM Y-27632. On day 1, the medium was replaced with the induction medium consisted of DMEM/F12 basal medium supplemented with 0.5% N2, 0.1 mM β-mercaptoethanol (β-ME), 2 μM SB431542, and 1 μM CHIR99021. This induction medium was refreshed every 1-2 days depending on cell density. On day 6, the medium was replaced with induction medium containing 1 μM retinoic acid. On day 7, cells were dissociated using Accutase and transferred to low-adhesion dishes to form free-floating neurospheres. These neurospheres were maintained in complete enteric neural crest cell medium, composed of a 1:1 mixture of DMEM/F12 and Neurobasal media, supplemented with 0.5% N2, 1% B27, 1 mM L-Glutamine (L-Glu), 0.1 mM β-ME, 10 ng/mL basic fibroblast growth factor, and 10 ng/mL epidermal growth factor. Y-27632 was added on the day of neurosphere dissociation to prevent apoptosis. The ENCC induction medium was refreshed every two days^[60]^.

### Differentiation of ENCCs into enteric neurons

ENCC-derived neurospheres (maintained in suspension for >5 days) were plated onto matrigel-coated dishes and cultured in peripheral neuron induction medium, composed of a 1:1 mixture of DMEM/F12 and Neurobasal media, supplemented with 1% N2, 2% B27, 1 mM L-glutamine, 0.1 mM β-mercaptoethanol (β-ME), 10 ng/mL BDNF, 10 ng/mL GDNF, and 0.2 mM ascorbic acid. The culture medium was replaced with fresh medium every two days^[60,62–64]^. Under these conditions, ENCCs differentiated into enteric neuron-like cells over the course of 1-2 weeks.

### Differentiation of ENCCs into mesenchymal stem cells (MSCs)

Cells with mesenchymal characteristics were generated as previously described, with some modifications^[62,63,65–67]^. ENCCs neurospheres after 2 days in suspension were plated onto matrigel-coated dishes and first allowed to adhere in the standard ENCCs expansion medium for 24 hours. The following day (designated day 1 of differentiation), the medium was replaced with MSCs medium. Cultures were maintained in this medium for 3-4 weeks. Over this period, cells adopted a spindle-shaped, fibroblast-like morphology characteristic of mesenchymal cells.

### Tri-lineage differentiation of MSCs

The multipotency of derived MSC-like cells was evaluated by inducing differentiation into osteogenic, adipogenic, and chondrogenic lineages. Cells with mesenchymal characteristics were induced to differentiation into osteogenic, adipogenic, and chondrogenic lineages as previously described^[62,68,69]^. MSCs were digested with 0.25% trypsin-EDTA and plated onto culture dishes. Upon reaching 80% confluency, the culture medium was replaced with osteogenic induction medium adipogenic induction medium, or chondrogenic induction medium. Each induction was carried out for 3 weeks, with the respective differentiation media replaced every 3-4 days.

### Plasmid transfection in hiPSCs

hiPSCs were seeded in 6-well plates and cultured at 37 °C with 5% CO_2_. When cells reached ∼30-40% confluency, they were transfected with 4 μg of a target plasmid (containing a hygromycin resistance cassette) along with 8 μg of the helper plasmid pTG9.3 using Lipofectamine™ Stem reagent (Invitrogen, cat. no. STEM00008), following the manufacturer’s protocol.

### siRNA transfection

For siRNA knockdown experiments, hiPSCs and ENCCs at ∼50-60% confluency were transfected with 20 nM siRNA using Lipofectamine™ Stem reagent according to the manufacturer’s instructions. HEK 293T cells were transfected with 20 nM siRNA using Lipofectamine™ 3000 under similar conditions. In all cases, the culture medium was replaced with fresh medium 24 hours post-transfection. Knockdown efficiency was assessed 48 hours after transfection by quantitative RT-PCR.

### Migration assay

#### (i) Wound healing assay

Cells were plated in 12-well plates at a density of 1×10^6^ cells per well and grown to ∼90% confluency in cell culture medium. A linear scratch (wound) was then created in the cell using a sterile 200 μL pipette tip. Detached cells were removed by gently washing with PBS. Wound closure was monitored by capturing phase-contrast images at 0, 16, and 24 hours after scratch. Migration was quantified by measuring the width of the wound at each time point and calculating the percentage of closure relative to the initial wound width^[70]^.

#### (ii) Transwell invasion assay

Cell invasion was examined using matrigel-coated Transwell inserts with 8 μm pore filters. The upper chamber of each insert was seeded with 5×10^4^ cells in serum-free medium, while the lower chamber contained complete medium as a chemoattractant. After 24 hours of incubation, non-invaded cells on the upper surface of the filter were gently removed. Cells that had migrated to the underside of the membrane were fixed in 4% PFA and stained with 0.1% crystal violet. The membranes were then rinsed with water, and images of stained cells (invaded cells on the underside) were acquired under a microscope. Invasion was evaluated by counting cells or measuring the stained area on the membrane in multiple fields^[70]^.

### Immunofluorescence

For multiplex immunofluorescent histology on tissue sections, a tyramide signal amplification kit was used according to the manufacturer’s protocol. Briefly, paraffin-embedded tissue sections were deparaffinized and subjected to heat-induced antigen retrieval in buffers of pH appropriate for each primary antibody: citrate buffer, pH 6.0; EDTA buffer, pH 9.0; or a universal HIER buffer. After antigen retrieval, endogenous peroxidase activity was quenched with 3% H_2_O_2_, and sections were blocked with 10% normal goat serum. Primary antibody staining and subsequent tyramide signal amplification were performed as per kit instructions, followed by counterstaining with DAPI. Stained sections were imaged using a Nikon CSU-W1 SoRa super-resolution microscope under standard settings.

For immunocytochemistry of cultured cells, cells grown on glass coverslips were fixed with 4% PFA (Servicebio, cat. no. G1101) for 20 minutes at room temperature. After fixation, cells were permeabilized with PBS containing 0.1% Triton X-100 and blocked with 10% goat serum. Cells were then incubated overnight at 4 °C with the following primary antibodies. After primary incubation, cells were washed and incubated for 1 hour at room temperature with appropriate species-specific secondary antibodies conjugated to Alexa Fluor 488, 594, or 647. Nuclei were counterstained with DAPI for 10 minutes. Images were acquired on a Leica SP8 confocal microscope.

### RNA isolation and qRT-PCR

Total RNA was extracted from human tissues and cells using TRIzol^TM^ reagent. Fresh or frozen tissues were homogenized in TRIzol with a mechanical homogenizer on ice. After 5 minutes incubation, RNA extraction auxiliary reagent was added. After 10 minutes incubation, phases were separated by centrifugation. RNA was precipitated from the aqueous phase with isopropanol, washed with 75% ethanol, and dissolved in DEPC-treated water. The concentration and purity were determined with a microvolume spectrophotometer. For qRT-PCR, cDNA was synthesized from 1μg of total RNA using PrimeScript RT Master Mix. Real-time qPCR was performed using iTaq Universal SYBR^®^ Green Supermix. Each reaction was run in technical triplicate and analysed following the 2^-ΔΔCt^ method, normalized to internal controls as described previously^[71,72]^. Primer sequences are provided in **Table S10.**

### Protein extraction

#### (i) Total tissue lysates

Protein extraction from human tissue samples was performed using 1XRIPA lysis buffer supplemented with protease inhibitor cocktail at ∼500 μL buffer per 100 mg of tissue. Fresh or frozen tissues were pulverized in liquid nitrogen, then homogenized in ice-cold RIPA buffer and lysed on ice for 30 minutes with intermittent vortexing. Lysates were clarified by centrifugation (12,000×g, 20 minutes, 4 °C), and the supernatants were collected. Protein concentration was determined using a BCA Protein Assay Kit.

#### (ii) Total cellular protein lysates

Cultured cells were harvested and lysed in 1XRIPA buffer supplemented with protease inhibitor cocktail at ∼400 μL lysis buffer per 1×10^7^ cells. Lysates were vortexed thoroughly and incubated on ice for 10 minutes, then centrifuged (12,000×g, 20 minutes, 4 °C). The supernatants were collected and protein concentration measured by BCA assay (Fude Biological Technology, cat. no. FD2001).

#### (iii) Nuclear and cytoplasmic protein lysates

Fractionation of nuclear and cytoplasmic proteins was performed using the NE-PER Nuclear and Cytoplasmic Extraction kit according to the manufacturer’s instructions^[73,74]^. Briefly, cells were harvested and sequentially treated with cytoplasmic extraction buffer and nuclear extraction buffer as per the kit protocol. The resulting nuclear and cytosolic fractions were collected, and protein concentrations were determined by BCA assay.

### Chromatin extraction

Cells were harvested, and the chromatin was isolated with Chromatin Extraction Kit according to the manufacturer’s protocol. Cell pellets were lysed in the provided buffer (volume adjusted to 380 ul per 10 million cells). The chromatin was released by sonication on ice using Uibra Cell^TM^. The extracts were then clarified, and chromatin protein concentrations were quantified using a BCA protein assays kit^[75]^. Extracted chromatin samples were either used immediately for immunoprecipitation or stored at −80 °C.

### Co-immunoprecipitation (Co-IP) assay

For co-immunoprecipitation, total protein lysates were first prepared and quantified by BCA assay. Each Co-IP reaction used an equal amount of protein. Lysates were incubated with 3 μg of the target primary antibody (or 3 μg of a species-matched IgG isotype control) pre-bound to 30 μL of Protein G magnetic beads. Binding was carried out overnight at 4 °C with gentle rotation. The beads were then washed four times with cold IP lysis buffer to remove non-bound proteins. Immune complexes were eluted from the beads by boiling in 1× SDS loading buffer at 100 °C for 10 minutes. The eluted proteins were analyzed by SDS-PAGE and western blot as described below.

### Lectin enrichment of glycoproteins

Enrichment of core-fucosylated glycoproteins recognized by PSA lectin was performed using a biotin-labelled lectin enrichment approach. Briefly, total cellular proteins were extracted and quantified by BCA assay as above. Protein samples (typically ∼1 mg in 500 μL lysis buffer) were first pre-cleared with Streptavidin T1 magnetic beads for 3 hours at 4 °C to remove any proteins that might bind non-specifically to the beads. The pre-cleared supernatants were then incubated with biotinylated PSA lectin overnight at 4 °C with gentle mixing. In parallel, control samples were processed with buffer lacking lectin to assess background binding. After the overnight incubation, Streptavidin T1 beads were added to each sample and incubation was continued for an additional 6 hours at 4 °C to capture the biotin-PSA-glycoprotein complexes. The beads were then washed thoroughly (four times) with IP lysis buffer to remove unbound material. PSA-bound glycoproteins were eluted by boiling the beads in 1× SDS loading buffer at 100 °C for 10 minutes. The enriched glycoprotein samples were analyzed by PAGE and subsequently by either western blot or mass spectrometry as required.

### Western blotting

Proteins were electrophoresed on SDS-PAGE gels and transferred to PVDF membranes. And then membranes were blocked in 5% BSA at room temperature. After 1 hour, the membranes were incubated with the primary antibodies overnight at 4 °C. Membranes were then washed and incubated with peroxidase-conjugated secondary antibodies for 1 hour at room temperature. After three washes with TBST, bands were visualized using ECL chemiluminescent substrate substrate and imaged using a CCD imaging system.

### Lectin blotting

Lectin blot analysis was used to detect specific glycoconjugates on blotted proteins^[76]^. After SDS-PAGE, proteins were transferred to PVDF membranes. The membranes were blocked and incubated overnight at 4 °C with a biotin-conjugated lectin as the primary probe. The following day, the blots were washed and then incubated with HRP-conjugated streptavidin at a dilution of 1:5000 for 1 h at room temperature. After washing off unbound streptavidin, the lectin-bound glycoprotein bands were visualized by ECL and imaged in the same manner as western blots.

### Lectin blotting for NGI-1 treatment

To assess the effects of glycosylation inhibition on lectin binding, hiPSCs were treated with the small-molecule glycosylation inhibitor NGI-1. The hiPSCs cells were cultured in medium contained 5 μM NGI-1 for 3 days and then were harvested and proteins were extracted. Protein samples were quantified using a BCA assay kit. For subsequent blotting analyses, samples were processed directly as described previously.

### Lectin-based fluorescence imaging assays

#### (i) For tissues

Lectin fluorescence histochemistry on tissue sections was performed as described previously with some modifications^[77,78]^. Paraffin sections were deparaffinized and subjected to antigen retrieval using a universal HIER buffer. Endogenous peroxidase activity was blocked by a 10-minute treatment with 3% H_2_O_2_, followed by blocking in 10% goat serum containing 0.1% Triton X-100 to reduce non-specific binding. Different lectins were, respectively, applied to the sections and incubated for 1 hour at room temperature or overnight at 4 °C, followed by a fluorescein (Alexa Fluor 488)-conjugated secondary step using an anti-biotin antibody or streptavidin for 1 hour at room temperature. Nuclei were counterstained with DAPI for 10 minutes. Images were acquired on a Nikon CSU-W1 SoRa super-resolution microscope.

#### (ii) For cell lines

Cultured cells from various lineages, including hiPSCs, enteric neural crest cells (ENCCs), neural progenitor cells (NPCs), mesenchymal stem cells (MSCs), osteoblasts (OBs), chondrocytes (CCs), and adipocytes (ACs), were dissociated and plated onto glass-bottom confocal dishes and allowed to adhere overnight. Then cells were fixed with 4% PFA (Servicebio, cat. no. G1101) for 15-20 minutes, permeabilized with 0.1% Triton X-100, and blocked with 10% goat serum. Different lectins were, respectively, applied to the sections and incubated for 1 hour at room temperature or overnight at 4 °C, followed by a fluorescein (Alexa Fluor 488)-conjugated secondary step using an anti-biotin antibody or streptavidin for 1 hour at room temperature. Nuclei were counterstained with DAPI for 10 minutes. Fluorescence images were captured using a Nikon N-SIM super-resolution confocal system.

### Metabolic glycan labeling assays

Metabolic labeling of cell-surface glycans was carried out using azide-modified sugar precursors, followed by click-chemistry detection on western blots. hiPSCs were first cultured in regular PSCeasy II medium for 24 hours, then switched to medium containing one of the following azide sugars: 50 μM peracetylated N-azidoacetylmannosamine, 50 μM GDP-fucose azide, 25 μM UDP-GlcNAz, or 25 μM Ac_4_GlcNAz. Cells were incubated with these metabolic probes for 3 days to allow incorporation into glycoconjugates. After 3 days, cells were harvested and proteins were extracted (RIPA buffer as above).

### Zebrafish husbandry and embryo rearing

All studies involving zebrafish were reviewed by the Animal Research Advisory Committee of the South China University of Technology, Guangzhou, China. All zebrafish studies were performed using AB strain zebrafish (*Danio rerio*) and conducted in compliance with institutional animal use guidelines. Embryos and larvae were raised at 28.5 °C in E3 medium (5 mM NaCl, 0.17 mM KCl, 0.33 mM CaCl2, 0.33 mM MgSO4, pH 7.2) under standard light-dark cycles (28 °C, 14/10 hour light/dark cycle). Developmental stages (hpf, hours post-fertilization; or dpf, days post-fertilization) were determined according to morphological criteria^[79]^. From 24 hpf forwards, embryos were treated with 0.003% 1-phenyl-2-thiourea (PTU), which prevents melanin synthesis, to inhibit pigment formation for imaging.

### Embryo micro-injections

To overexpress *fut8a* in zebrafish, we injected one-cell-stage embryos with 2 nL (150 ng/μL) of a pcs2 plasmid containing the zebrafish *fut8a* gene. Control embryos were injected with the empty pcs2 vector at the same dose. Embryos were then raised in E3 medium at 28.5 °C and assessed for expression and phenotypes.

### Fluorescent tracer gut transit assay

To evaluate gastrointestinal motility in zebrafish larvae, we performed a feeding and transit assay with fluorescent microspheres as a tracer. This stage (5∼6 dpf) corresponds to the period shortly after yolk absorption when larvae transition to exogenous feeding. A fluorescent tracer was prepared by mixing ∼100 μL of yellow-green fluorescent 2.0 μm polystyrene microspheres with ∼100 mg of live *Paramecium* culture (wet biomass), as previously described^[80]^. The microsphere-*Paramecium*mixture was then diluted into 10 mL of embryo medium (E3 medium) with gentle vortexing. The final tracer suspension contained approximately 1×E-04 microspheres/mL and 10 mg/mL of *Paramecium*, providing both a food source to stimulate ingestion and a visual fluorescent marker.

For the transit assay, groups of zebrafish larvae (∼6 dpf) were placed in multi-well plates with the prepared tracer suspension. Approximately 10 larvae were incubated in 1 mL of tracer per well. Larvae were allowed to feed on the fluorescent tracer at 28.5 °C for 2 hours. After this feeding period, larvae were removed from the tracer and rinsed, then transferred to fresh embryo water (without tracer) and kept in the dark at 28.5 °C for an additional 4 hours to allow gut transit of the ingested material. At the end of the 6-hour experiment, larvae were anesthetized with tricaine and examined under a fluorescence stereomicroscope. The distribution of fluorescent microspheres was quantified to assess intestinal transit. Larvae showing extensive movement of the tracer to the distal intestine were considered to have normal motility, whereas retention of tracer in anterior regions indicated delayed transit.

### Whole-mount immunofluorescent staining of zebrafish

To visualize enteric neurons in zebrafish larvae, we performed whole-mount immunostaining on larval gut tissue. Prior studies have shown that enteric neurons express the neuronal proteins HuC and HuD^[81]^, so we used an anti-HuC/D antibody to label enteric neurons. Zebrafish larvae at 5 dpf were fixed overnight in 4% PFA at 4 °C. After fixation, larvae were washed three times with PBS containing 1% Triton X-100 and equilibrated in distilled water for 5 minutes. Then they were then permeabilized in cold acetone (−20 °C) for 1 hour, re-equilibrated in water, and washed again in PBST. Non-specific binding was blocked by incubating larvae in 5% normal goat serum in PBST for 1 hour at room temperature. Larvae were then incubated with rabbit anti-HuC/D primary antibody for 96 hours at 4 °C with gentle agitation. After extensive washing in PBST (3 times over ∼45 minutes), larvae were incubated with Alexa Fluor 488 goat anti-rabbit IgG secondary antibody in PBST overnight at 4 °C. DAPI was also included with the secondary antibody to label nuclei. Finally, larvae were washed in PBST and mounted in 80% glycerol or suitable mounting medium for imaging. Enteric neurons were quantified from confocal z-stacks (Zeiss LSM 800) by counting HuC/D+ cells in the distal intestine using ImageJ. Counts were standardized by focusing on a defined segment length (a 4-somite-length region proximal to the anus) to allow comparison between individuals.

### Generation of an induced transgenic mouse line with conditional overexpression of Fut8 specifically in Sox10 positive cells (*Fut8*-iCTG)

Using CRISPR/Cas9 technology, we inserted the CAG-LSL-*Fut8*-WPRE-pA expression cassette into the Rosa26 locus via homologous recombination. This resulted in the generation of Rosa26-targeted knock-in heterozygous mice, termed R26-e(CAG-LSL-*Fut8*-WPRE-pA)1, which enable conditional overexpression of the *Fut8* gene. To generate an induced transgenic mouse line with conditional overexpression of *Fut8* specifically in enteric neural crest lineage cells (*Sox10* positive cells). LSL-*Fut8*/+ mice were self-crossed to generate LSL-*Fut8*/LSL-*Fut8* mice, while they were also bred with *Sox10*-CreERT2/+ mice (Strain #:027651, The Jackson laboratory) to produce LSL-*Fut8*/+; *Sox10*-iCreERT2/+ offspring. Then LSL-*Fut8*/LSL-*Fut8* mice were crossed to LSL-*Fut8*/+; *Sox10*-iCreERT2/+ mice to yield LSL-*Fut8*/LSL-*Fut8*; *Sox10*-iCreERT2/+ progeny, as well as LSL-*Fut8*/LSL-*Fut8* mice used as Cre-negative controls. The line was established and maintained by crossing LSL-*Fut8*/LSL-*Fut8*; *Sox10*-iCreERT2/+ mice with LSL-*Fut8*/LSL-*Fut8* mice. Mice were treated with tamoxifen to induce *Fut8* overexpression in enteric neural crest lineage cells (*Sox10* positive cells) at E7.5 with 50 ug/g tamoxifen. To evaluate the therapeutic effect of the DIP, we administered it to the mice from embryonic day 7.5 (E7.5) through postnatal day 30 (P30).

### Carmine gavage and red feces excretion of mouse

Mice (experimental/control groups) were housed in IVC under controlled conditions (22 ± 1 °C, 50 ± 10% humidity). After an 8-hour fast (with water provided ad libitum), the mice received a 1% (w/v) carmine (Sigma-Aldrich, St. Louis, MO, Batch Number: SHBL4031) suspension, prepared by homogenizing 0.5 g carmine powder in PBS using vortex mixing. Mice were orally gavaged with 200 μL of the suspension, and the administration time was documented. They were then returned to clean cages with ad libitum water. Following gavage, fecal output was monitored every 10 minutes. The gastrointestinal transit time was defined as the time from gavage completion to the expulsion of the first carmine-stained fecal pellet.

### Intestinal contrast-enhanced Micro-CT imaging of mouse

Following an 8-hour fast with ad libitum water access, experimental mice were orally gavaged with 200 μL of 30% meglumine diatrizoate (Hanfeng Pharmaceutical, Shanghai, China; Lot# H20034058) to enhance intestinal contrast. After allowing 2 hours for contrast agent distribution, mice were anesthetized by an intraperitoneal injection of tribromoethanol (10 μL/g body weight; Nanjing Aibei Bio-Technology Co., Ltd., Jiangsu, China; Lot# 2039A). Following anesthesia, the animals were placed in the micro-CT scanner.

### Whole-mount immunofluorescence staining of mouse

After dissecting the distal colon longitudinally along the mesentery, residual feces were removed, and the mucosa and submucosa layers were carefully dissected under a stereomicroscope. Tissues were fixed in 4% PFA for 2 h at room temperature, followed by washing with PBS. Penetration was achieved using pre-chilled 100% acetone at −20 °C for 1 h, followed by two rinses with ddH_2_O and three washes using 1% Triton X-100 in PBS. Endogenous peroxidase activity was blocked with 3%H_2_O_2_ at room temperature for 1 h, followed by three PBST washes. Tissues were incubated with blocking buffer (1% Triton X-100, 15% goat serum in PBS) for 2 h. Primary antibodies were incubated for 4 days at 4 °Cin darkness. After PBST washes, secondary antibodies were incubated overnight at 4 °C. After washing with PBST, the samples were dehydrated through a graded glycerol series (30%, 50%, 80%) and stored overnight at −20 °C.

### SILAC labeling

For heavy isotope labeling, ENCCs were cultured in DMEM:F-12 (Cytiva, cat. no. SH30023.01) supplemented with 0.5% N2, 0.1 mM β-mercaptoethanol (β-ME), 2 μM SB431542 (APExBIO, cat. no. A8249), and 1 μM CHIR99021, 13C615N2-Lys and 13C615N4-Arg at concentrations of 100 ug/mL at 37 °C in a humidified incubator with 5% CO2 for 0.5, 1, 2, 4, 6, 8 days, respectively. For unlabeled cells, ENCCs were grown in DMEM/F12 basal medium supplemented with 0.5% N2, 0.1 mM β-mercaptoethanol (β-ME), 2 μM SB431542, and 1 μM CHIR99021. Heavy isotope-labeled cells and unlabeled control cells were harvested separately for membrane and nuclear protein extraction.

### Membrane extraction

The membrane extraction method described here was adapted from previously established protocols^[82,83]^. Following collection and lysis of cells or tissues, the cell membrane is isolated by first removing the nucleus and mitochondria through low-speed centrifugation. The membrane fraction is then pelleted via ultracentrifugation. To further purify the membrane preparation, sodium carbonate solution is introduced to dissociate and solubilize noncovalently bound membrane-associated proteins, leaving a purified membrane pellet after subsequent centrifugation. This approach requires minimal sample preparation, is compatible with mass spectrometry (MS) analysis, and yields a membrane pellet suitable for downstream processing.

### LC-MS/MS analysis for SILAC proteomics

200 ng of digested peptides were analyzed using a nanoElute ultra-high-performance liquid chromatography (UHPLC) system (Bruker) coupled to the timsTOF Pro 2 mass spectrometer (Bruker) and were separated at a flow rate of 300 nL/min using a 60-minute gradient on a 20 cm analytical column. The mobile phase B consisted of 0.1% formic acid in acetonitrile (ACN), and its gradient comprised three linear segments: from 2 to 22% in 40 min, from 22 to 37% in 10 min, and from 37 to 80% in 5 min, with an additional 5-minute 80% sustain for analytical column washing. All separation processes were performed in an integrated toaster column oven at 50 °C. DDA was performed in PASEF18 mode with 10 PASEF MS/MS scans. The capillary voltage was set to 1500 V and the spectra were acquired in the range of m/z from 100 to 1700 Th with an ion mobility range (1/K0) from 0.6 to 1.6 Vs/cm^2^. The ramp and accumulation time were set to 100 ms to achieve a duty cycle close to 100% and a total cycle time of 1.1 s. The collision energy was ramped linearly as a function of mobility from 59 eV at 1/K0 = 1.6 Vs/cm2 to 20 eV at 1/K0 = 0.6 Vs/cm^2^. Precursors with charge state from 0 to 5 (If the charge state of a peptide precursor is 0, it indicates that the isotope ion was not detected for that peptide precursor) were selected with the target value of 20,000 and intensity threshold of 5000. Any precursors that reached the target value in arbitrary units were dynamically excluded for 0.4 min. dia-PASEF data was analyzed using DIA-NN software (v 2.2.0)^[84]^ with a precursor mass tolerance of 10 ppm and a fragment mass tolerance of 20 ppm. Carbamidomethylation of cysteine was set as a fixed modification, and methionine oxidation and N-terminal acetylation as variable modifications. A maximum of 2 missed trypsin cleavages was allowed.

### LC-MS/MS analysis of PSA-enriched and L1CAM-immunoprecipitated proteins

Protein bands of interest (for example, proteins enriched by PSA lectin or co-immunoprecipitated with L1CAM) were identified by liquid chromatography-tandem mass spectrometry (LC-MS/MS). Gel bands were excised from SDS-PAGE gels and subjected to in-gel tryptic digestion as described previously with some modification^[85]^. Briefly, excised gel pieces were destained and then reduced with 5 mM dithiothreitol (DTT) for 30 minutes at 56 °C, followed by alkylation with 11 mM iodoacetamide for 1 hour at room temperature in the dark. The proteins were digested overnight at 37 °C with sequencing-grade modified trypsin in 50 mM ammonium bicarbonate. Peptides were extracted twice from the gel (each extraction with 50% acetonitrile/1% TFA for 1 hour), pooled, and dried in a SpeedVac. Peptides were reconstituted in ∼20 μL of 0.1% TFA, and any insoluble debris was removed by a 15 minute, 20,000×g centrifugation at 4 °C. The peptide samples were then analyzed by nanoLC-MS/MS.

Chromatographic separation was performed on a Thermo Dionex Ultimate 3000 HPLC system using a custom-packed C18 reversed-phase capillary column (75 μm ID × 150 mm, 5 μm particles, 300 Å pore). Peptides were eluted at ∼300 nL/min with a 40-minute linear gradient from 0% to ∼40% solvent B (0.1% formic acid in acetonitrile) in solvent A (0.1% formic acid in water). The eluted peptides were directly introduced into an LTQ-Orbitrap mass spectrometer (Thermo Fisher) using nano-ESI. The mass spectrometer was operated in data-dependent acquisition mode: full MS1 scans (m/z 300-1500) were acquired in the Orbitrap at 120,000 resolution, followed by MS/MS (HCD fragmentation at 40% NCE) of the most intense precursor ions (typically top 10-15 peaks, dynamic exclusion enabled) in the ion trap. MS/MS spectra were searched against the UniProt human protein database using Proteome Discoverer software (v1.4). Search parameters included trypsin specificity (up to 2 missed cleavages), fixed modification of cysteine (carbamidomethylation), variable oxidation of methionine, and precursor and fragment mass tolerances of ±10 ppm and ±0.8 Da, respectively. Peptide spectrum matches were filtered to a 1% false discovery rate (FDR).

### LC-MS/MS analysis for Glycoproteomics

For global N-glycoproteomics, a mass spectrometry workflow was employed involving protein extraction, enrichment of N-glycosylated peptides, LC-MS/MS acquisition, and bioinformatic analysis followed by previously established protocols^[82]^:

**(i) Total protein extraction:** Samples (cell or tissue) were flash-frozen and ground to a powder in liquid nitrogen followed by the S-trap (ProtiFi) manufacturer instructions with minor modifications. Briefly, lysis was performed in a buffer containing 100 mM NH_4_HCO_3_ (pH 8.0), 8 M urea, and 0.2% SDS. Lysates were ultrasonicated on ice (e.g., 5 min total sonication time in short pulses) to ensure thorough disruption. Debris was removed by centrifugation (12,000×g, 15 minutes, 4 °C), and the supernatant containing solubilized proteins was transferred to a clean tube. Proteins were reduced with 5 mM DTT for 1 hour at 56 °C and alkylated with 11 mM iodoacetamide for 1 hour at room temperature in the dark. After alkylation, four volumes of pre-chilled acetone were added to each sample, and proteins were precipitated at −20 °C for at least 2 hours. Precipitates were collected by centrifugation, washed twice with cold acetone, and air-dried. The protein pellets were then re-dissolved in dissolution buffer (0.1 M triethylammonium bicarbonate (TEAB), pH 8.5, 6 M urea) for digestion.
**(ii) N-glycosylated peptide enrichment:** Protein solutions were diluted to 500 μL with DB lysis buffer (8 M urea, 100 mM TEAB, pH 8.5). Trypsin (MS-grade) was added at a ratio of ∼1:50 (enzyme:substrate) along with additional TEAB to a final ∼2 mM, and samples were digested for 4 hours at 37 °C. A second aliquot of trypsin (again 1:50) and CaCl₂ (1 mM final) were then added, and digestion proceeded overnight (∼16 hours) at 37 °C. The next day, formic acid was added to acidify each digest to pH < 3. The acidified peptide mixtures were cleared by centrifugation (12,000×g, 5 minutes) and loaded onto C18 desalting columns. Columns were washed (three bed volumes of 0.1% formic acid, 3% acetonitrile), and peptides were eluted with buffer containing 70% acetonitrile, 0.1% formic acid. The eluted peptides were lyophilized to dryness. For glycopeptide enrichment, dried peptides were reconstituted and applied to a hydrophilic interaction liquid chromatography (HILIC) column (HILICON) to enrich N-glycopeptides. The column was washed with 80% acetonitrile (in water with 1% trifluoroacetic acid) to retain glycopeptides, then N-glycopeptides were eluted with 0.1% trifluoroacetic in water. The enriched N-glycopeptide fraction was collected and dried for LC-MS/MS analysis.
**(iii) LC-MS/MS acquisition:** lyophilized samples were resuspended in 20 μL of loading buffer (0.1% formic acid in water). After centrifugation (12,000×g, 10 min, room temp), ∼10 μL of the supernatant was injected for each run. Peptides were separated using an EASY-nLC 1000 UHPLC system. The chromatography setup included a trap column (Thermo Acclaim PepMap 100, 2 cm × 100 μm, 3 μm) and an analytical C18 column. Mobile phase A was 0.1% formic acid in water; mobile phase B was 0.1% formic acid in acetonitrile. Peptides were eluted with a linear gradient at a flow rate of ∼300 nL/min. Eluted peptides were electrosprayed (Nanospray Flex ion source, 2.3 kV, capillary 320 °C) into a Orbitrap Eclipse Tribrid mass spectrometer. Full MS scans (m/z 350-2000) were acquired in the Orbitrap at 60,000 resolution (at m/z 200) with an AGC target of 125% and a maximum injection time of 50 ms. The precursors were selected for stepped higher-energy collisional dissociation (HCD) MS/MS (normalized collision energy 30± 10%). MS/MS spectra were acquired in the orbitrap, using an AGC target of 800% and a max injection time of 250 ms. Raw data files were generated for each run.
**(iv) Data analysis:** The LC-MS/MS data for glycopeptides were analyzed using pGlyco software (v3.0)^[86]^. Searches were performed with a precursor mass tolerance of 10 ppm and a fragment mass tolerance of 20 ppm. Carbamidomethylation of cysteine was set as a fixed modification, and methionine oxidation and N-terminal acetylation as variable modifications. A maximum of 2 missed trypsin cleavages was allowed. pGlyco identified N-glycopeptides and corresponding glycan compositions; results were filtered to 1% FDR at the glycopeptide level. Quantitative analysis of glycopeptides across samples was done using pGlycoQuant or similar software, and only high-confidence glycopeptides were considered for downstream interpretation.

### Chromatin immunoprecipitation sequencing (ChIP-seq)

Chromatin immunoprecipitation for sequencing (ChIP-seq) was performed essentially as described by Kim *et al*^[87]^. In brief, approximately 1×10^8^ cells were crosslinked with 1% formaldehyde (10 minutes at room temperature) to fix protein-DNA interactions, then quenched with 125 mM glycine. Cells were washed with cold PBS, and nuclei were isolated and resuspended in lysis buffer. Chromatin was sheared by sonication to an average size of ∼200-500 bp. Sheared chromatin from each sample was divided and incubated overnight at 4 °C with a specific antibody of interest. The antibodies were pre-bound to Protein G magnetic Dynabeads or, in the case of FLAG-tagged protein ChIP, to Anti-FLAG M2 magnetic beads. The following day, the beads were washed extensively with ChIP lysis buffer, and bound chromatin complexes were eluted. Crosslinks were reversed by heating at 65 °C (several hours or overnight) and proteins were digested (e.g., with proteinase K), after which DNA was purified (by phenol-chloroform or column purification). Enriched DNA from ChIP was prepared for sequencing as described below.

Purified ChIP DNA (typically 5-50 ng) was used to prepare sequencing libraries with an Illumina ChIP-seq library prep kit or equivalent, according to the manufacturer’s instructions. High-throughput sequencing was performed on an Illumina platform to generate 150 bp paired-end reads. The raw sequencing reads (fastq format) were trimmed to remove adapter sequences and low-quality bases using TrimGalore (v0.6.4) and then aligned to the human reference genome (hg38) using Bowtie2 (v2.4.5) with default parameters. Alignments were filtered to remove reads with low mapping quality (MAPQ < 30) and PCR duplicates using SAMtools (v1.18)^[88]^. Reads mapping to blacklisted genomic regions (ENCODE blacklist^[89]^) were also excluded using Bedtools. Peak calling was performed with MACS2 (v2.2.7)^[90]^ using the corresponding input DNA as background (parameters: -g hs -q 0.05 -m 5 50). The resulting peaks, representing enriched DNA binding sites or histone modification regions, were annotated to the nearest genes or genomic features. For visualization, bigWig files were generated and normalized to reads per million. Aggregate profile plots and heatmaps centered on features of interest were produced using Ngsplot (v2.63) for comparative analysis of ChIP-seq signal across conditions. Differential binding analysis between experimental groups was conducted by comparing peak intensities and genomic distributions; any notable differences were further investigated in the context of gene regulation.

### RNA-seq analysis

Transcriptome sequencing (RNA-seq) was performed on selected samples to profile gene expression changes under various conditions. Total RNA was extracted and quality-checked (RIN > 8) before library preparation. Poly(A)+ mRNA libraries were prepared using an Illumina TruSeq kit or similar, and sequenced on an Illumina platform to obtain paired-end reads (∼150 bp). Raw reads were trimmed to remove adapter sequences and low-quality bases using TrimGalore (v0.6.4)^[91]^. The cleaned reads were aligned to the human genome (GRCh38/hg38) using HISAT2 (v2.1.0) with default parameters^[91]^. Alignments were sorted and indexed with SAMtools. Gene-level read counts were obtained with featureCounts (v2.0.0)^[92,93]^ using gene annotations (Ensembl or GENCODE) corresponding to hg38. Downstream differential expression analysis was conducted in R using the DESeq2 package (v1.30.1)^[91]^. Low-expressed genes (for instance, those with reads per kilobase per million mapped reads (RPKM) < 1 in all samples) were filtered out to improve statistical power. DESeq2 was used to estimate size factors and dispersions, and to perform Wald tests for differential expression between conditions. Genes with an adjusted p-value < 0.05 and an absolute log_2 fold-change ≥ 1.5 were considered significantly differentially expressed^[94]^. Lists of differentially expressed genes were further analyzed for enrichment of biological pathways and Gene Ontology terms to gain insight into the affected molecular processes.

---

**Figure S1.**
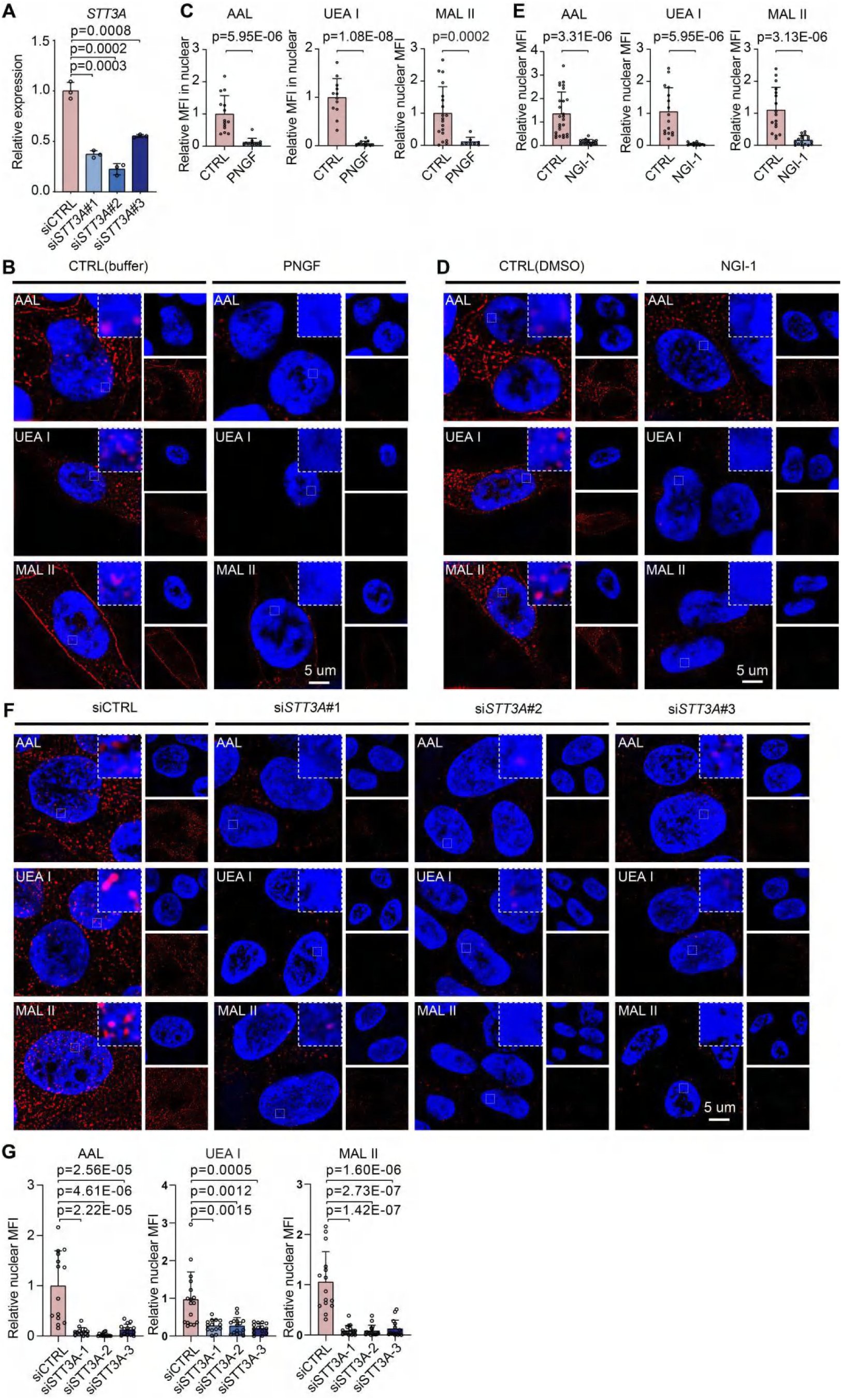
Widespread nuclear localization of complex N-glycoproteins in hiPSCs (lectin-based detection, related to Figure 1. (**A**) RT-qPCR validation of effective siRNA-mediated *STT3A*knockdown in hiPSCs compared to control (CTRL). Significance was determined by t-test (two tailed). **(B-G)** Super-resolution fluorescence imaging of hiPSCs stained with indicated lectins (AAL, UEA I, MAL II, shown in red) recognizing nuclear N-glycosylation. Nuclei stained with DAPI (blue). Cells were subjected to PNGF (**B**), NGI-1 (**D**), or si*STT3A*(**F**) treatment as indicated. CTRL, control. Scale bars, 5 μm. Statistical analysis was assessed by t-test (two tailed) (**C**) (**E**) (**G**). MFI, mean fluoresence intensity.

**Figure S2.**
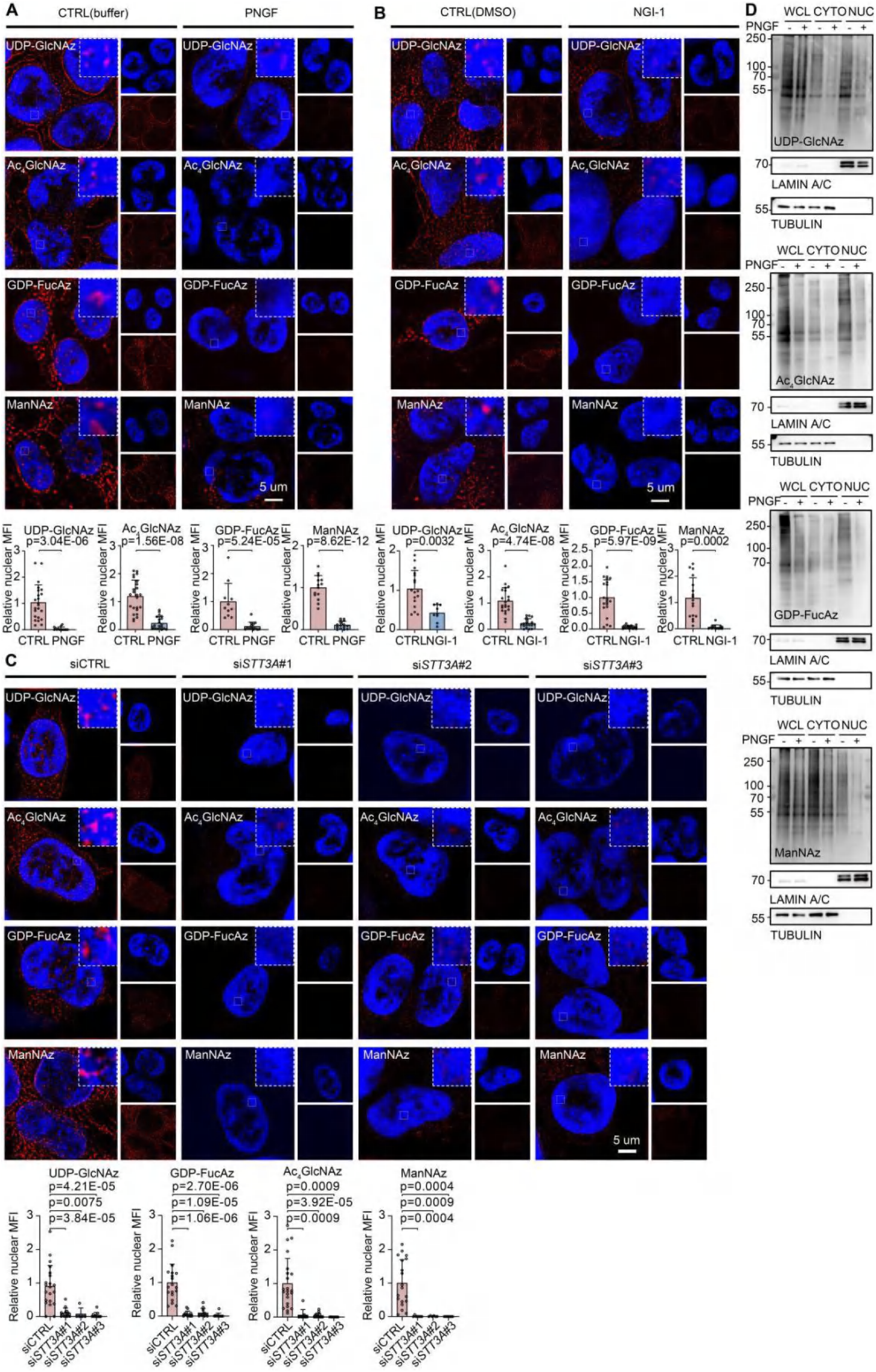
Widespread nuclear localization of complex N-glycoproteins in hiPSCs (validation by metabolic labeling, related to. **Figure 1) (A-C)** Super-resolution fluorescence imaging of metabolic labeling (Ac_4_GlcNAc, ManNAz, shown in red). Nuclei stained with DAPI (blue). Cells were subjected to PNGF, NGI-1 or si*STT3A* treatment as indicated. CTRL, control. Scale bars, 5 μm. Statistical analysis was assessed by t-test (two tailed). MFI, mean fluoresence intensity. **(D)** Western blot analysis of whole-cell lysates (WCL), cytoplasmic (CYTO), and nuclear protein (NUC) fractions from metabolic labeling hiPSCs treated with PNGF. TUBULIN and LAMIN A/C served as cytoplasmic and nuclear markers, respectively.

**Figure S3.**
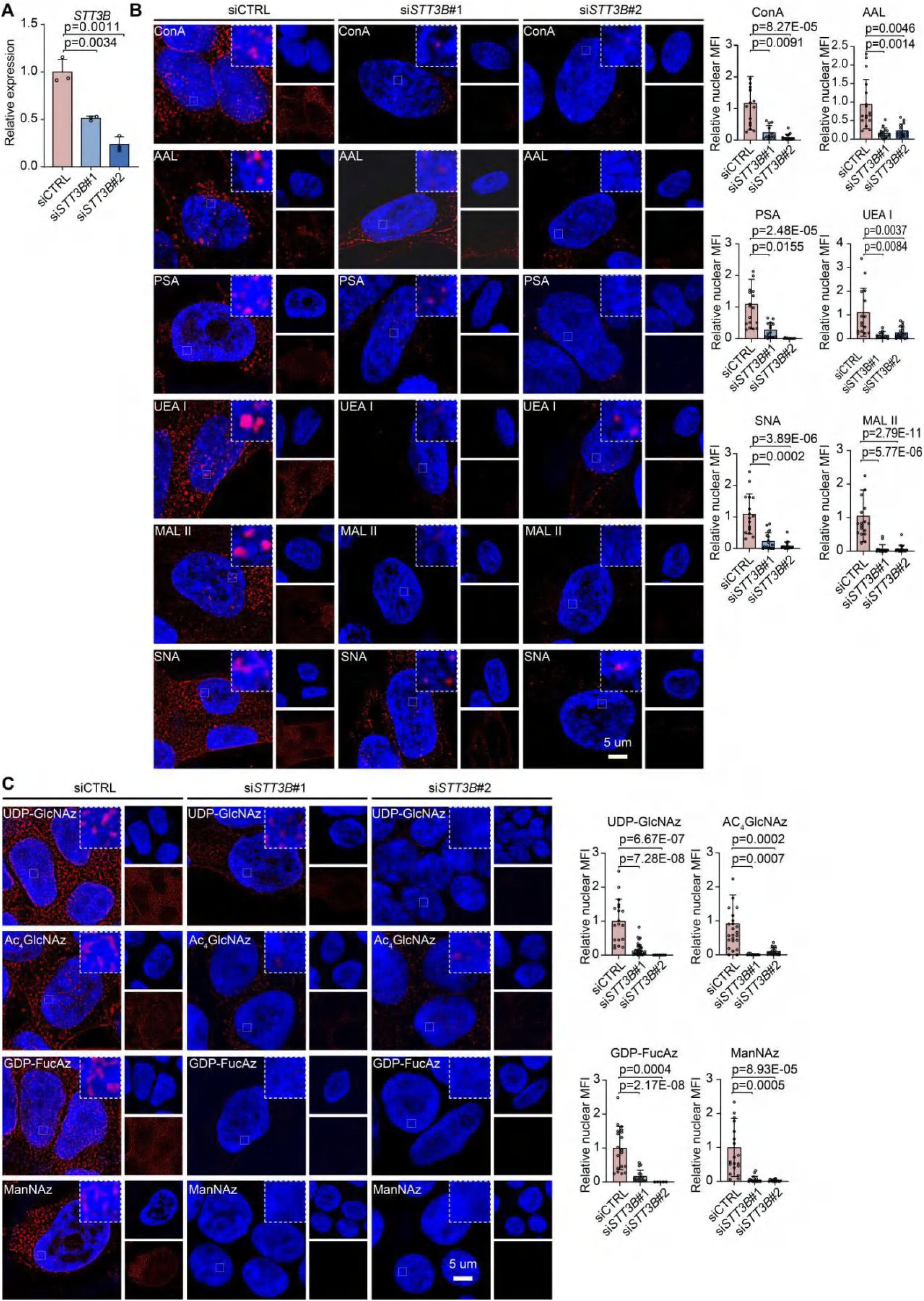
Effects of *STT3B* knockdown on nuclear N-glycoproteins in hiPSCs (related to Figure 1) **(A)** RT-qPCR analysis confirming efficient knockdown of *STT3B* via siRNA in hiPSCs compared to control. Data represent mean ± S.E.M from three independent experiments; significance assessed by t-test (two tailed). **(B)** Super-resolution fluorescence imaging of nuclear N-glycans (lectin staining shown in red) in hiPSCs after *STT3B*knockdown. Lectins used include ConA, PSA, AAL, UEA I, SNA, and MAL II. Nuclei are stained with DAPI (blue). Images are representative of three independent experiments. Images represent three independent experiments. Scale bars, 5 μm. Significance was determined by t-test (two tailed). MFI, mean fluoresence intensity. **(C)** Visualization of metabolically labeled nuclear glycans using azide-tagged sugar analog probes (UDP-GlcNAz, Ac4GlcNAz, GDP-FucAz, ManNAz) in hiPSCs subjected to *STT3B* knockdown. Significance was analysed via t-test (two tailed). Scale bars, 5 μm. MFI, mean fluoresence intensity.

**Figure S4.**
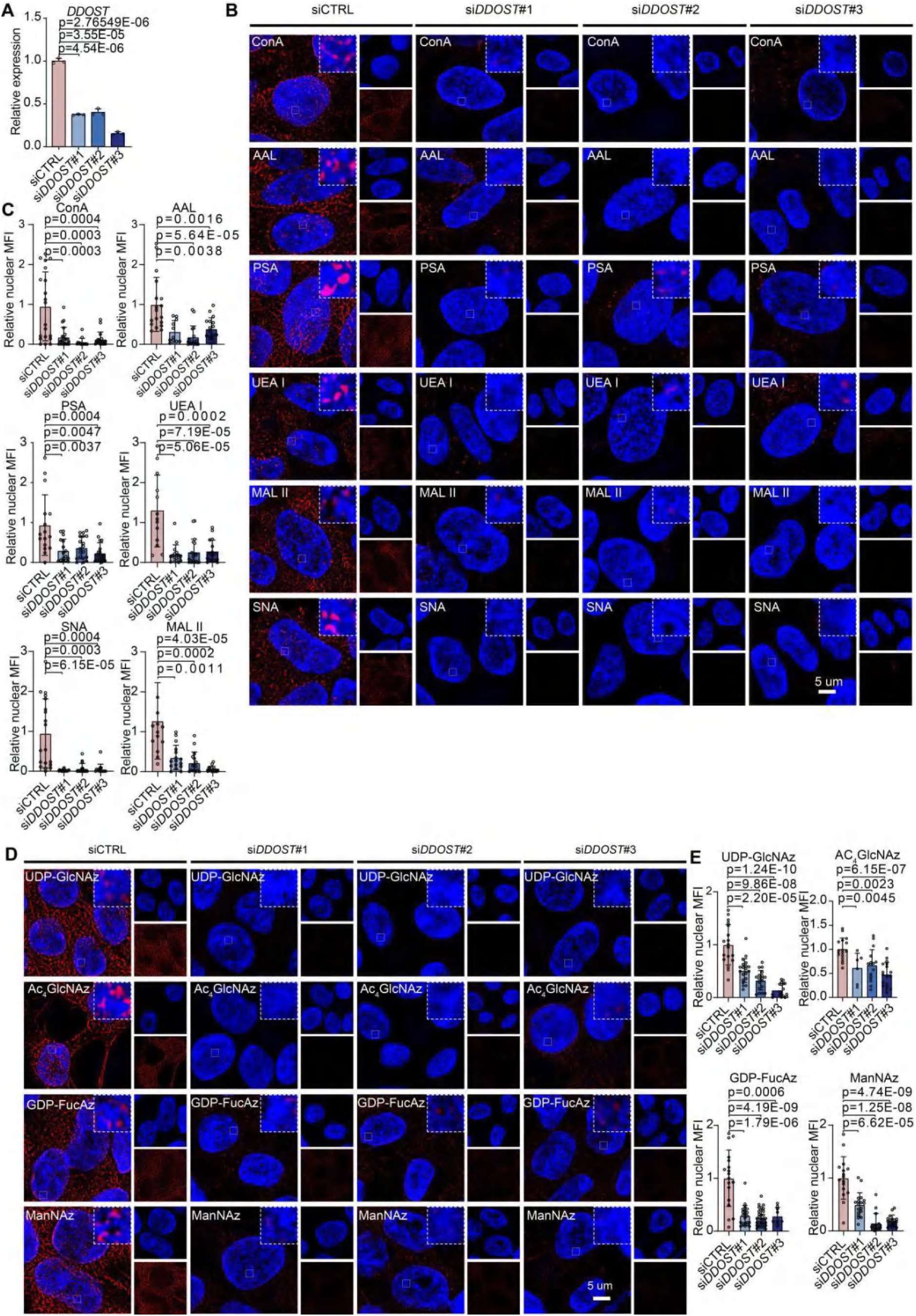
Effects of *DDOST* knockdown on nuclear N-glycoproteins in hiPSCs (related to Figure 1) **(A)** RT-qPCR analysis confirming efficient knockdown of *DDOST* via siRNA in hiPSCs compared to control. Data represent mean ± S.E.M from three independent experiments; significance assessed by t-test (two tailed). **(B** and **C)** Super-resolution fluorescence imaging of nuclear N-glycans (lectin staining shown in red) in hiPSCs after *DDOST*knockdown. Lectins used include ConA, PSA, AAL, UEA I, SNA, and MAL II. Nuclei are stained with DAPI (blue). Images are representative of three independent experiments. Images represent three independent experiments. Scale bars, 5 μm (**B**). Significance was determined by t-test (two tailed) (**C**). MFI, mean fluoresence intensity. **(D** and **E)** Visualization of metabolically labeled nuclear glycans using azide-tagged sugar analog probes (UDP-GlcNAz, Ac_4_GlcNAz, GDP-FucAz, ManNAz) in hiPSCs subjected to *DDOST*knockdown. Significance was determined via t-test (two tailed). Scale bars, 5 μm (**D**). Significance was determined by t-test (two tailed) (**E**). MFI, mean fluoresence intensity.

**Figure S5.**
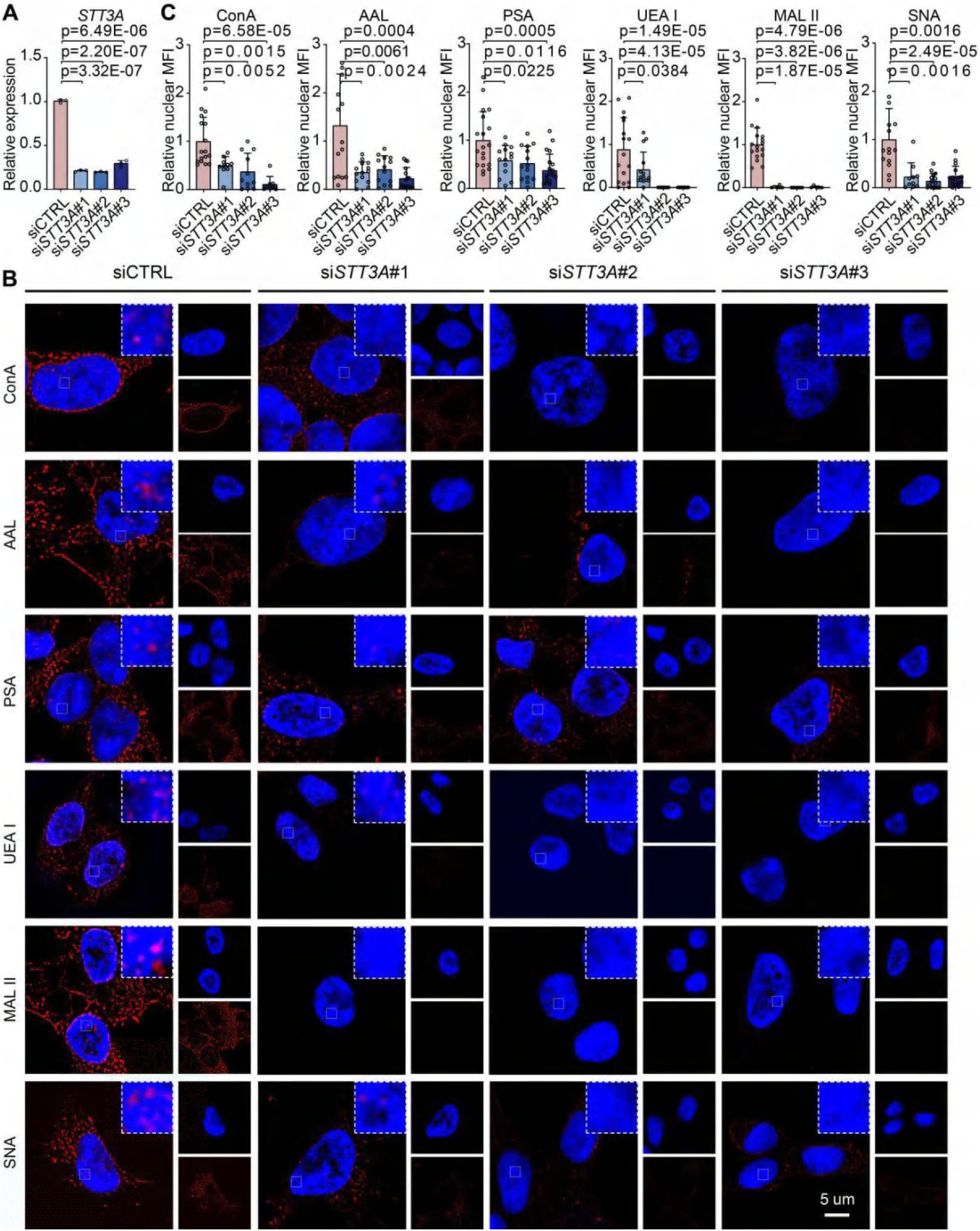
*STT3A* knockdown reduces nuclear glycosylation in 293T cells (lectin-based detection, related to Figure 1) **(A)** RT-qPCR analysis confirming efficient siRNA knockdown of STT3A in 293T cells compared to control (CTRL). **(B)** Super-resolution fluorescence imaging using indicated lectins (ConA, PSA, AAL, UEA I, SNA, MAL II, shown in red) demonstrates reduced nuclear glycosylation signals in 293T following knockdown of *STT3A*. Nuclei stained with DAPI (blue). **(C)** Quantification of nuclear glycosylation intensity corresponding to images in **(B)**. Statistical significance was determined by t-test (two tailed). Scale bars, 5 μm.

**Figure S6.**
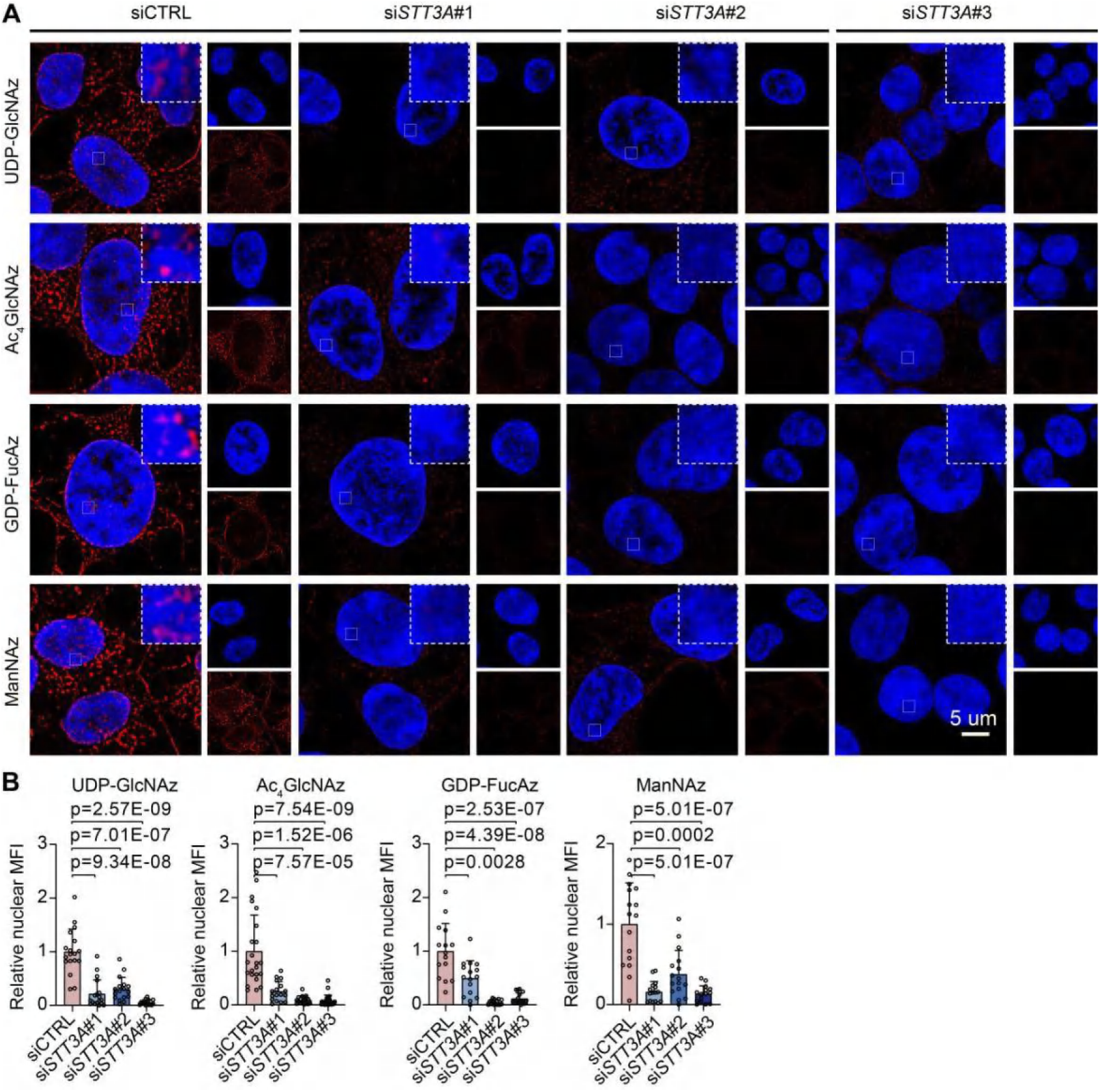
*STT3A* knockdown reduces nuclear glycosylation in 293T cells (validation by metabolic labeling, related to Figure 1) **(A)** Super-resolution imaging of metabolically labeled nuclear glycans in 293T cells with *STT3A* knockdown. Azide-functionalized glycans (UDP-GlcNAz, Ac_4_GlcNAz, GDP-FucAz, ManNAz, shown in red) were used to label nascent N-glycans, and nuclei stained with DAPI (blue). **(B)** Quantification of nuclear glycosylation intensity corresponding to images in **(A)**. Statistical significance was determined by t-test (two tailed). Scale bars, 5 μm. MFI, mean fluoresence intensity.

**Figure S7.**
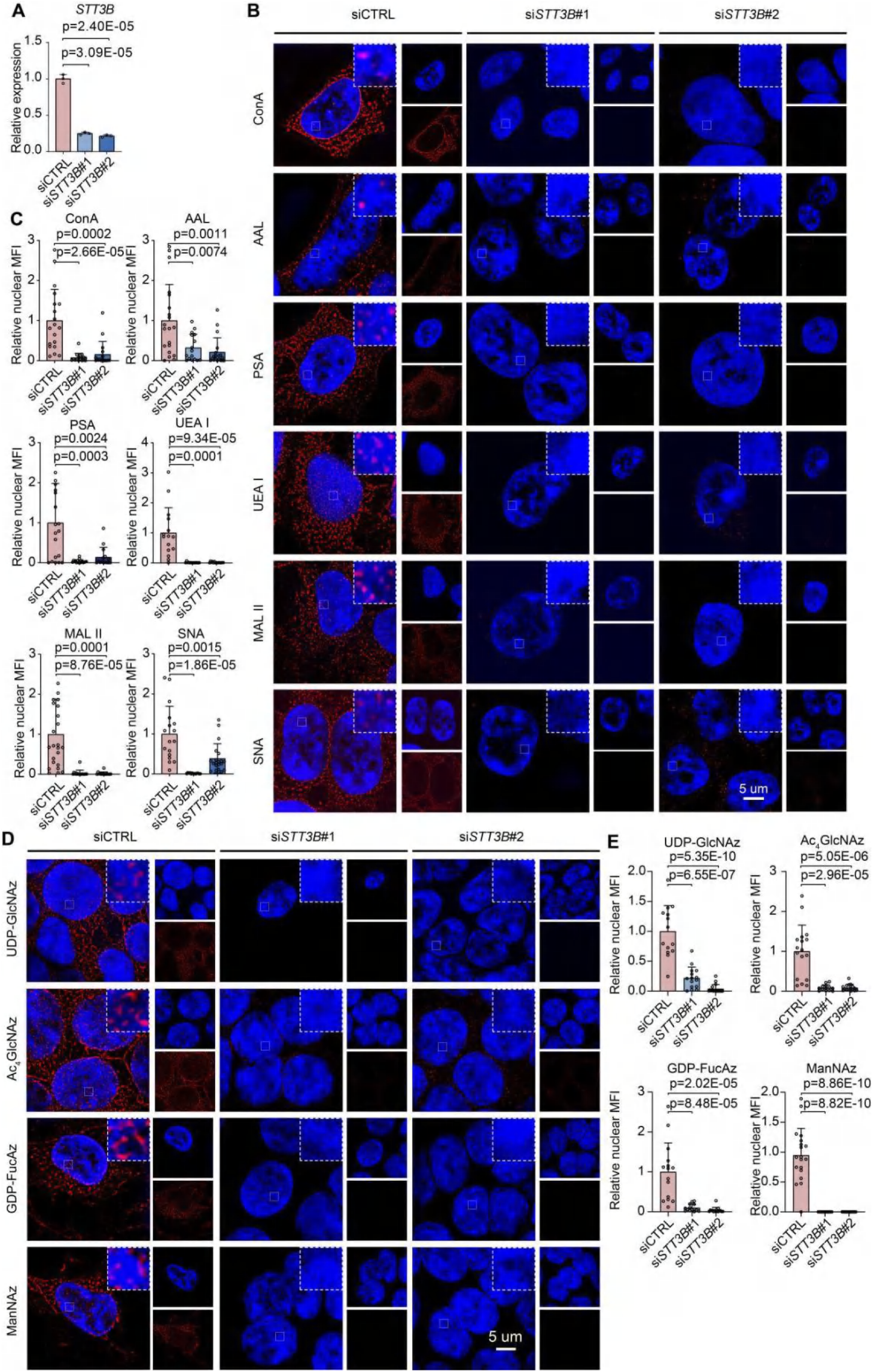
*STT3B* knockdown reduces nuclear glycosylation in 293T cells (lectin-based detection and validation by metabolic labeling, related to Figure 1) **(A)** RT-qPCR analysis confirming efficient siRNA knockdown of *STT3B* in 293T cells compared to control (CTRL). **(B)** Super-resolution fluorescence imaging using indicated lectins (ConA, PSA, AAL, UEA I, SNA, MAL II, shown in red) demonstrates reduced nuclear glycosylation signals in 293T following knockdown of *STT3B*. Nuclei stained with DAPI (blue). **(C)** Quantification of nuclear glycosylation intensity corresponding to images in **(B)**. Statistical significance was determined by t-test (two tailed). Scale bars, 5 μm. MFI, mean fluoresence intensity. **(D)** Super-resolution imaging of metabolically labeled nuclear glycans in 293T cells with *STT3B* knockdown. Azide-functionalized glycans (UDP-GlcNAz, Ac_4_GlcNAz, GDP-FucAz, ManNAz, shown in red) were used to label nascent N-glycans, and nuclei stained with DAPI (blue). **(E)** Quantification of nuclear glycosylation intensity corresponding to images in **(D)**. Statistical significance was determined by t-test (two tailed). Scale bars, 5 μm. MFI, mean fluoresence intensity.

**Figure S8.**
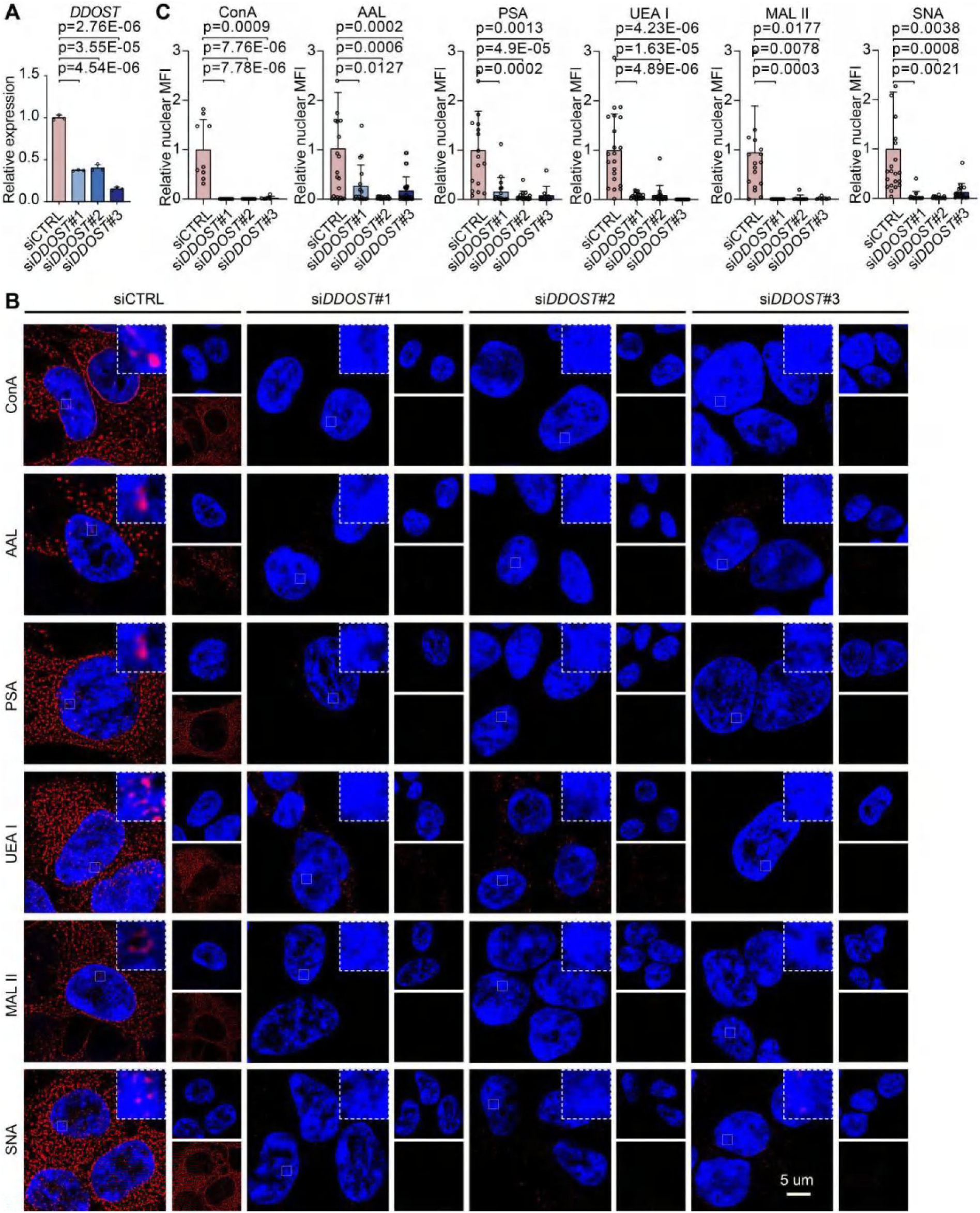
*DDOST* knockdown reduces nuclear glycosylation in 293T cells (lectin-based detection, related to Figure 1) **(A)** RT-qPCR analysis confirming efficient siRNA knockdown of *DDOST* in 293T cells compared to control (CTRL). **(B)** Super-resolution fluorescence imaging using indicated lectins (ConA, PSA, AAL, UEA I, SNA, MAL II, shown in red) demonstrates reduced nuclear glycosylation signals in 293T following knockdown of *DDOST*. Nuclei stained with DAPI (blue). **(C)** Quantification of nuclear glycosylation intensity corresponding to images in **(B)**. Statistical significance was determined by t-test (two tailed). Scale bars, 5 μm. MFI, mean fluoresence intensity.

**Figure S9.**
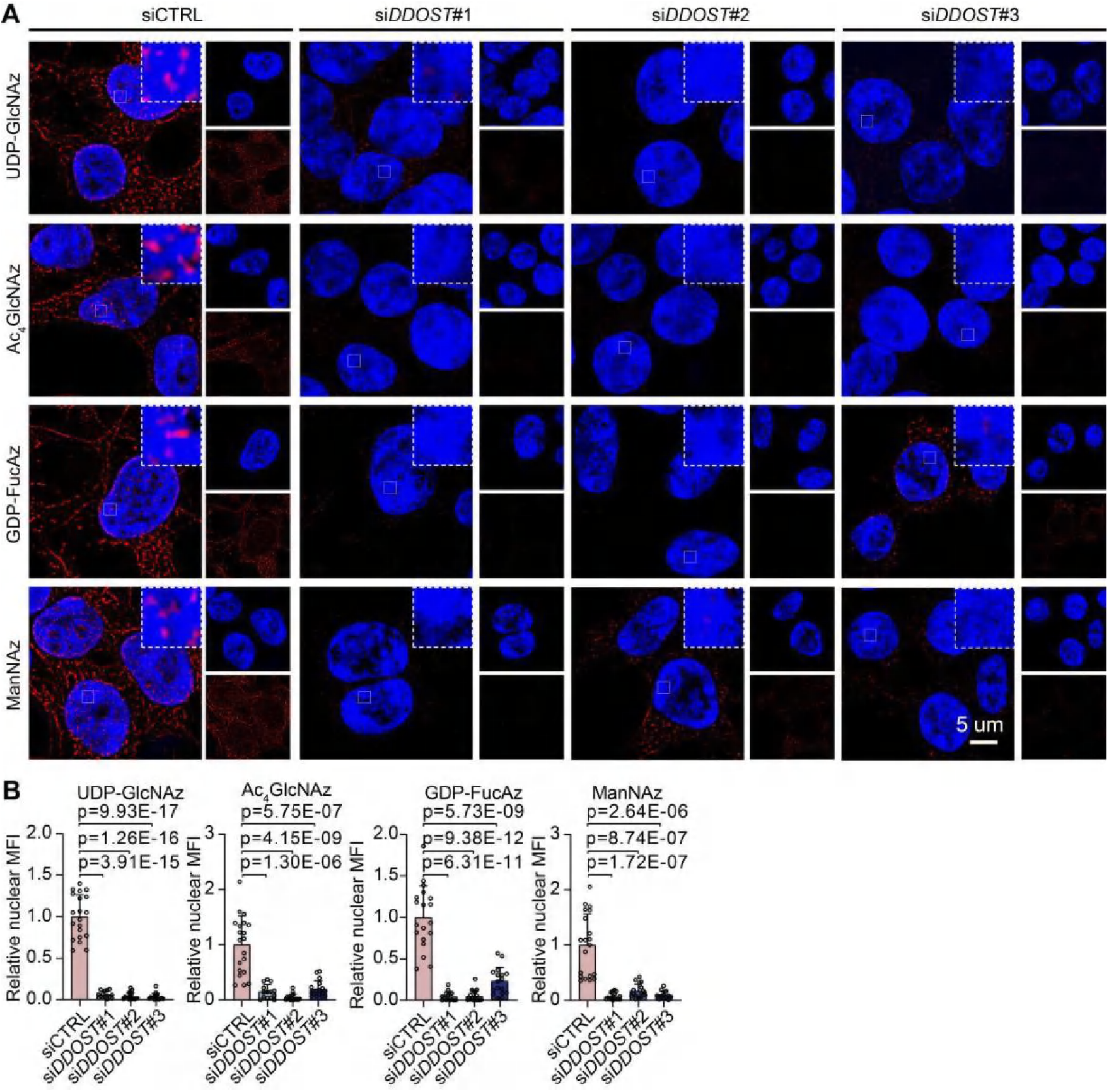
*DDOST* knockdown reduces nuclear glycosylation in 293T cells (validation by metabolic labeling, related to Figure 1) **(A)** Super-resolution imaging of metabolically labeled nuclear glycans in 293T cells with *DDOST* knockdown. Azide-functionalized glycans (UDP-GlcNAz, Ac_4_GlcNAz, GDP-FucAz, ManNAz, shown in red) were used to label nascent N-glycans, and nuclei stained with DAPI (blue). **(B)** Quantification of nuclear glycosylation intensity corresponding to images in **(A)**. Statistical significance was determined by t-test (two tailed). Scale bars, 5 μm. MFI, mean fluoresence intensity.

**Figure S10.**
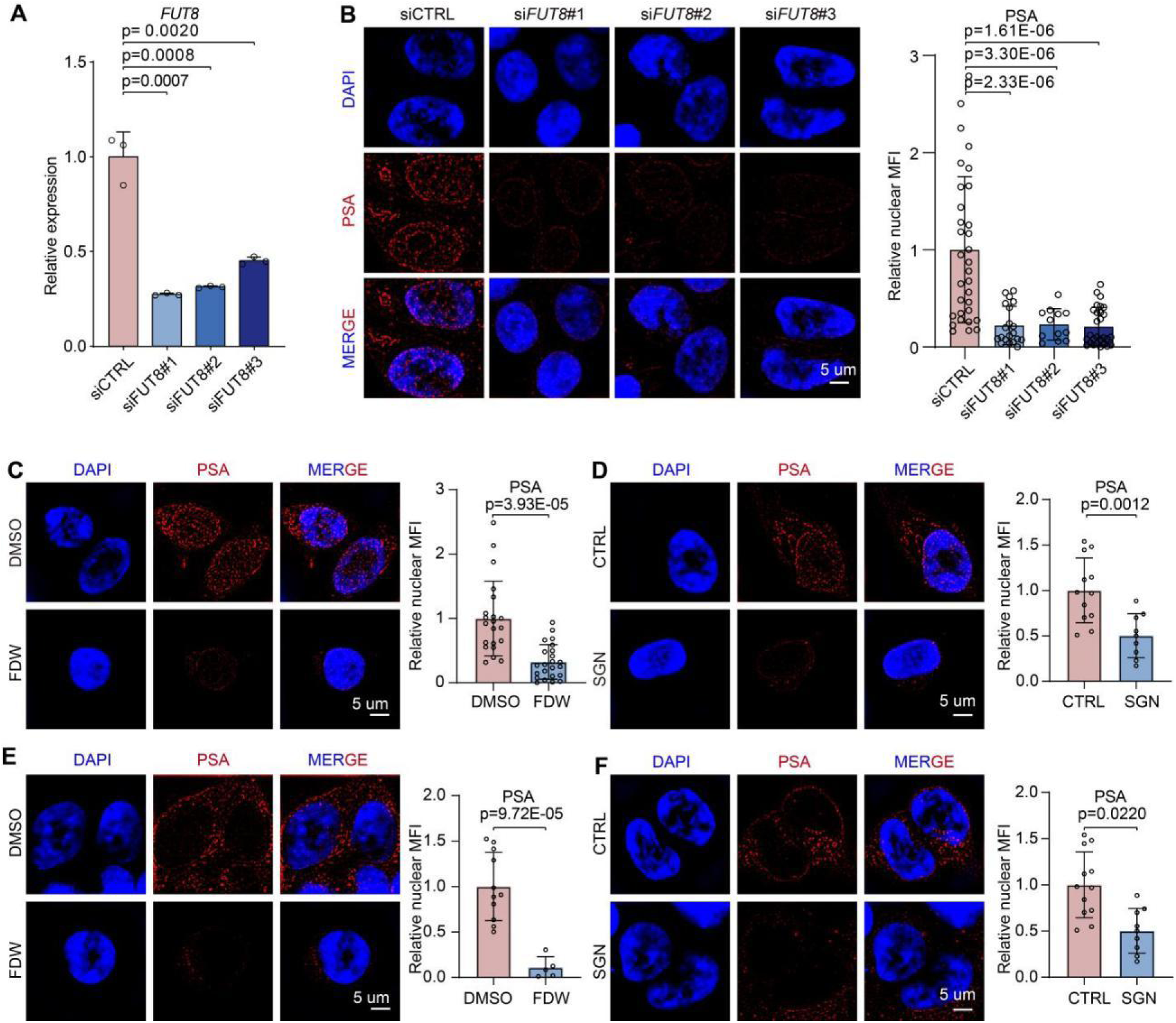
FUT8 inhibition validates PSA lectin specificity for core-fucosylation (related to Figure 1) **(A)** RT-qPCR confirming efficient *FUT8*knockdown in hiPSCs (si*FUT8*vs control). Statistical significance analyzed by t-test (two tailed). **(B)** Super-resolution fluorescence imaging of nuclear PSA staining intensity in hiPSCs following *FUT8*knockdown, showing a significant reduction in nuclear signal. **(C-F)** Super-resolution fluorescence imaging demonstrating reduced nuclear PSA staining (red) upon FUT8 loss of function. hiPSCs (**C** and **D**) and 293T (**E** and **F**) cells treated with FUT8 inhibitors FDW or SGN display markedly reduced nuclear PSA signals compared to controls. Nuclei stained with DAPI (blue). Images are representative of three independent experiments; scale bars, 5 μm. Statistical significance was determined by t-test (two tailed). MFI, mean fluoresence intensity.

**Figure S11.**
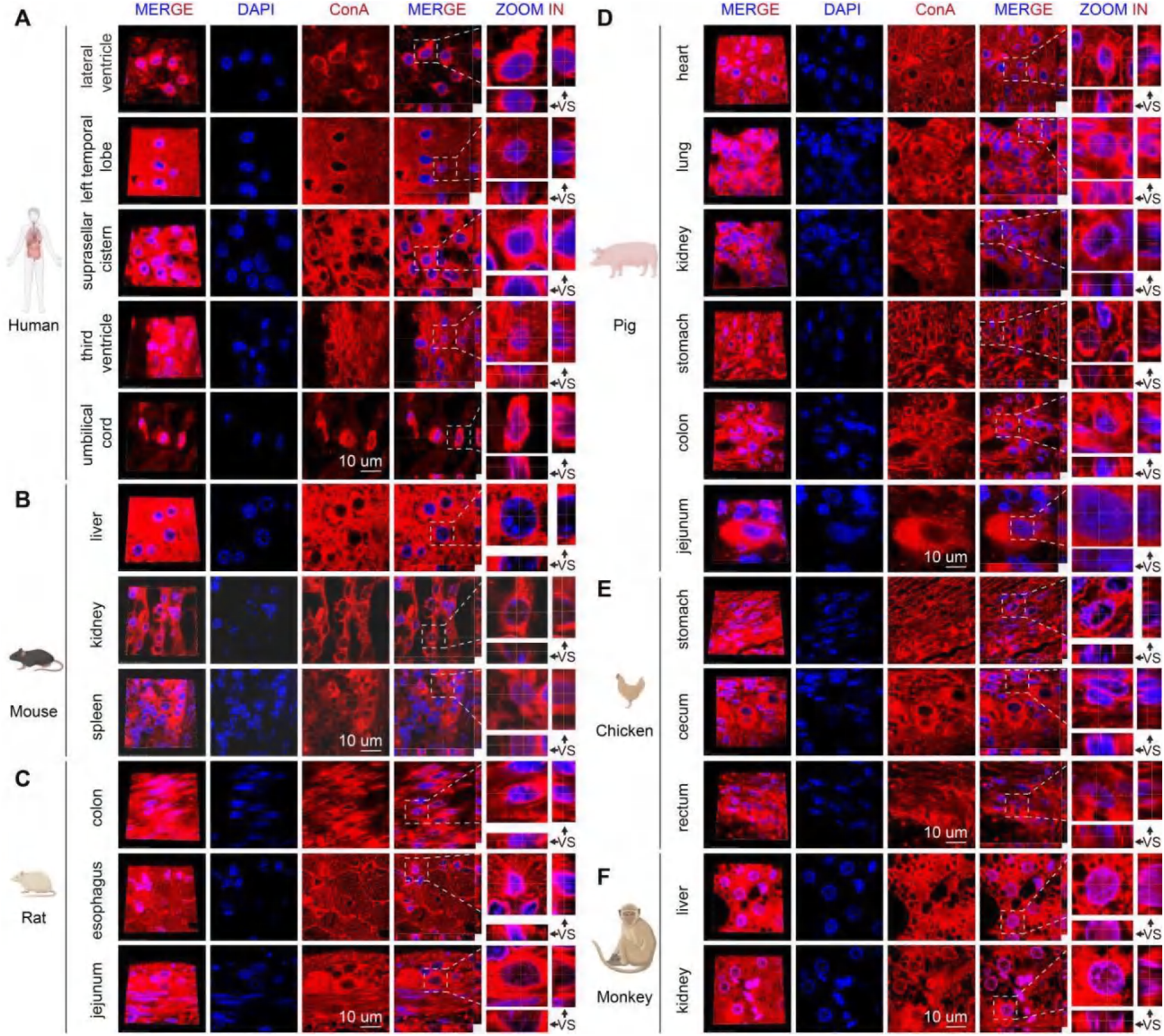
ConA lectin staining reveals nuclear glycosylation across multiple species (related to Figure 2) **(A-F)** Fluorescence images of tissue sections from human (**A**), mouse (**B**), rat (**C**), pig (**D**), chicken (**E**), and monkey (**F**) stained with ConA (red) show clear nuclear glycan signals across species. Nuclei are counterstained with DAPI (blue). Images are representative of three independent experiments. VS, vertical section; scale bars, 10 μm.

**Figure S12.**
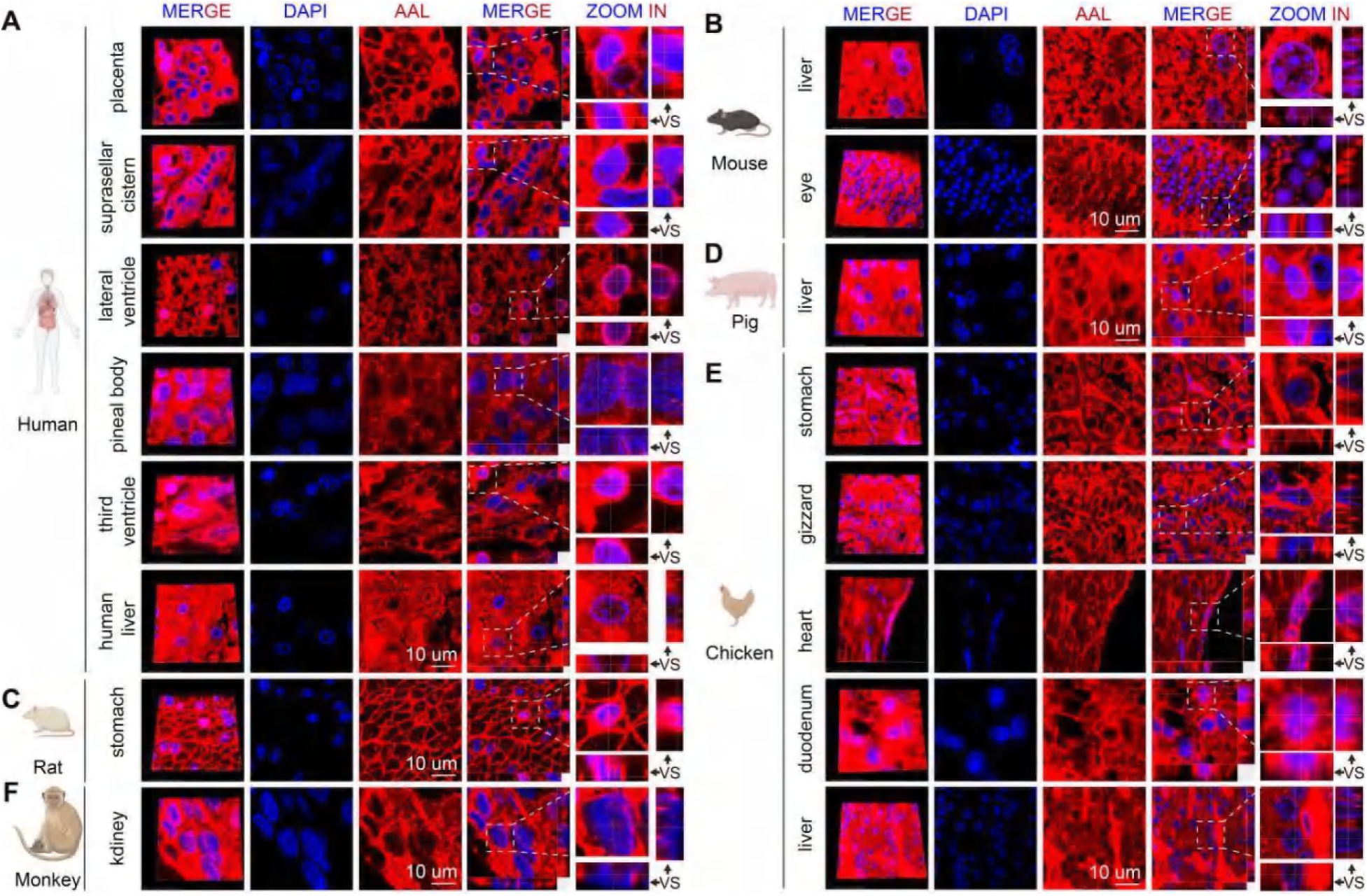
AAL lectin staining reveals nuclear glycosylation across multiple species (related to Figure 2) **(A-F)** Fluorescence images of tissue sections from human (**A**), mouse (**B**), rat (**C**), pig (**D**), chicken (**E**), and monkey (**F**) stained with AAL (red) show clear nuclear glycan signals across species. Nuclei are counterstained with DAPI (blue). Images are representative of three independent experiments. VS, vertical section; scale bars, 10 μm.

**Figure S13.**
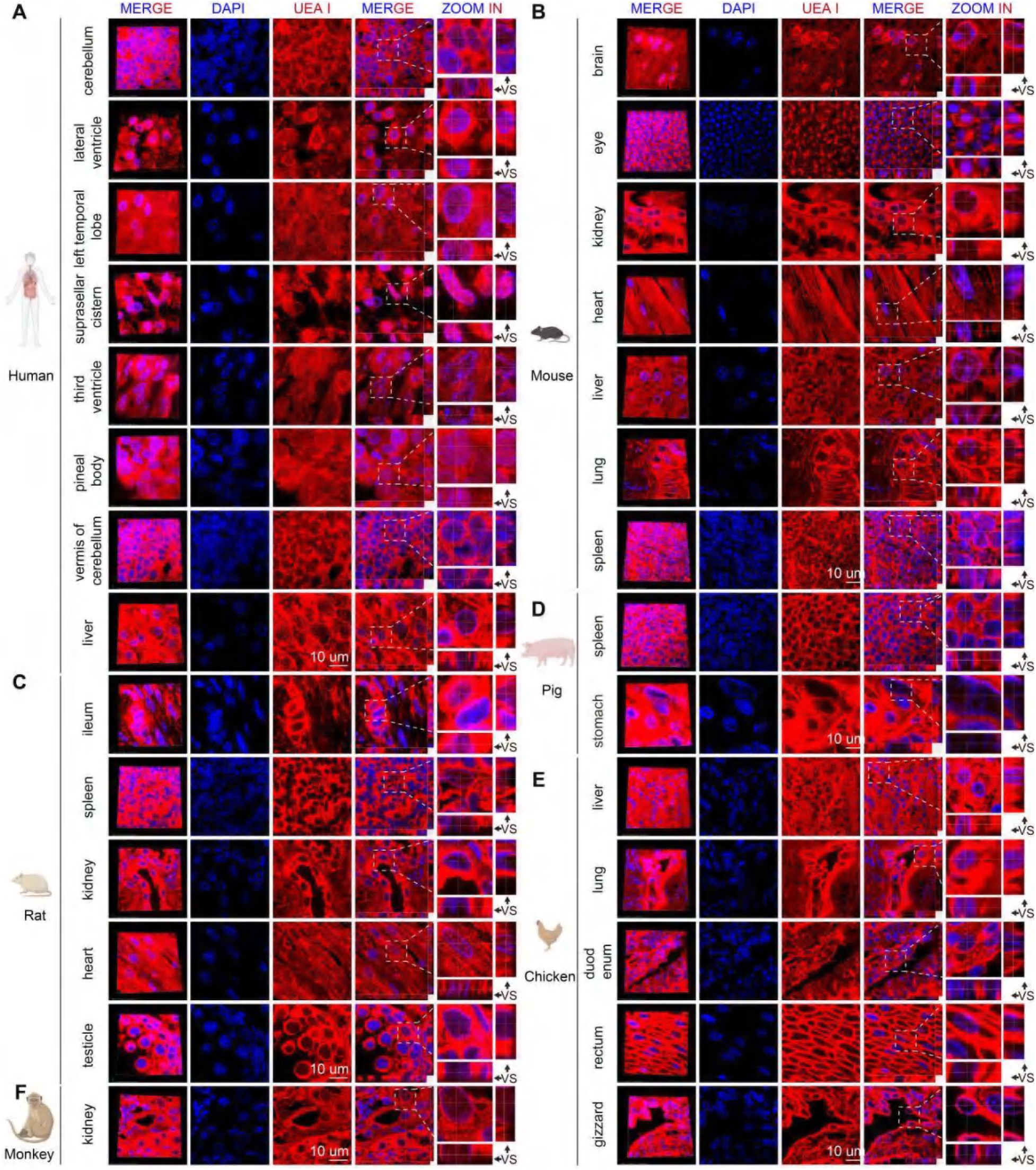
UEA I lectin staining reveals nuclear glycosylation across multiple species (related to Figure 2) **(A-F)** Fluorescence images of tissue sections from human (**A**), mouse (**B**), rat (**C**), pig (**D**), chicken (**E**), and monkey (**F**) stained with UEA I (red) show clear nuclear glycan signals across species. Nuclei are counterstained with DAPI (blue). Images are representative of three independent experiments. VS, vertical section; scale bars, 10 μm.

**Figure S14.**
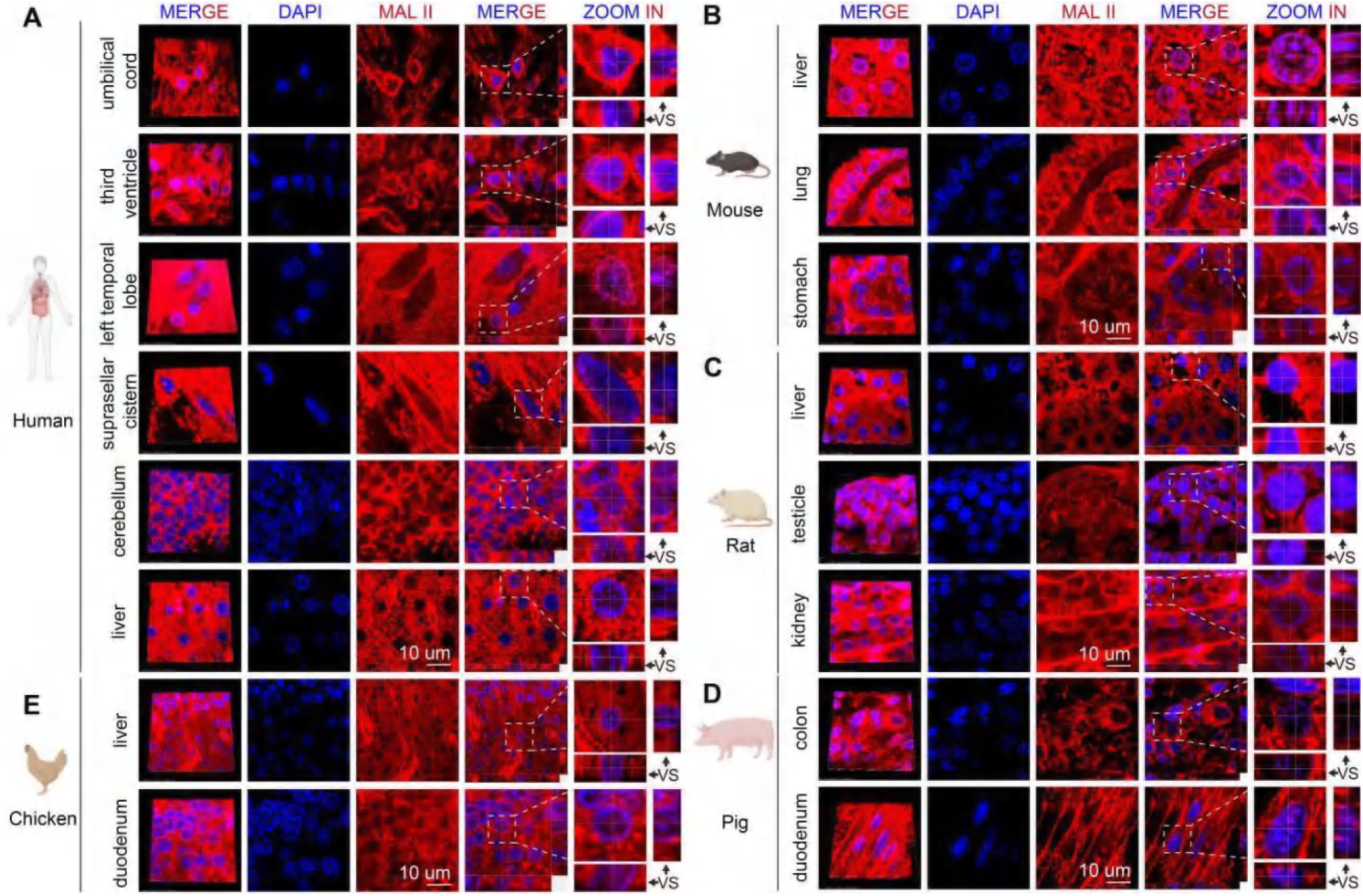
MAL II lectin staining reveals nuclear glycosylation across multiple species (related to Figure 2) **(A-E)** Fluorescence images of tissue sections from human (**A**) mouse (**B**), rat (**C**), pig (**D**), and chicken (**E**) stained with MAL II (red) show clear nuclear glycan signals across species. Nuclei are counterstained with DAPI (blue). Images are representative of three independent experiments. VS, vertical section; scale bars, 10 μm.

**Figure S15.**
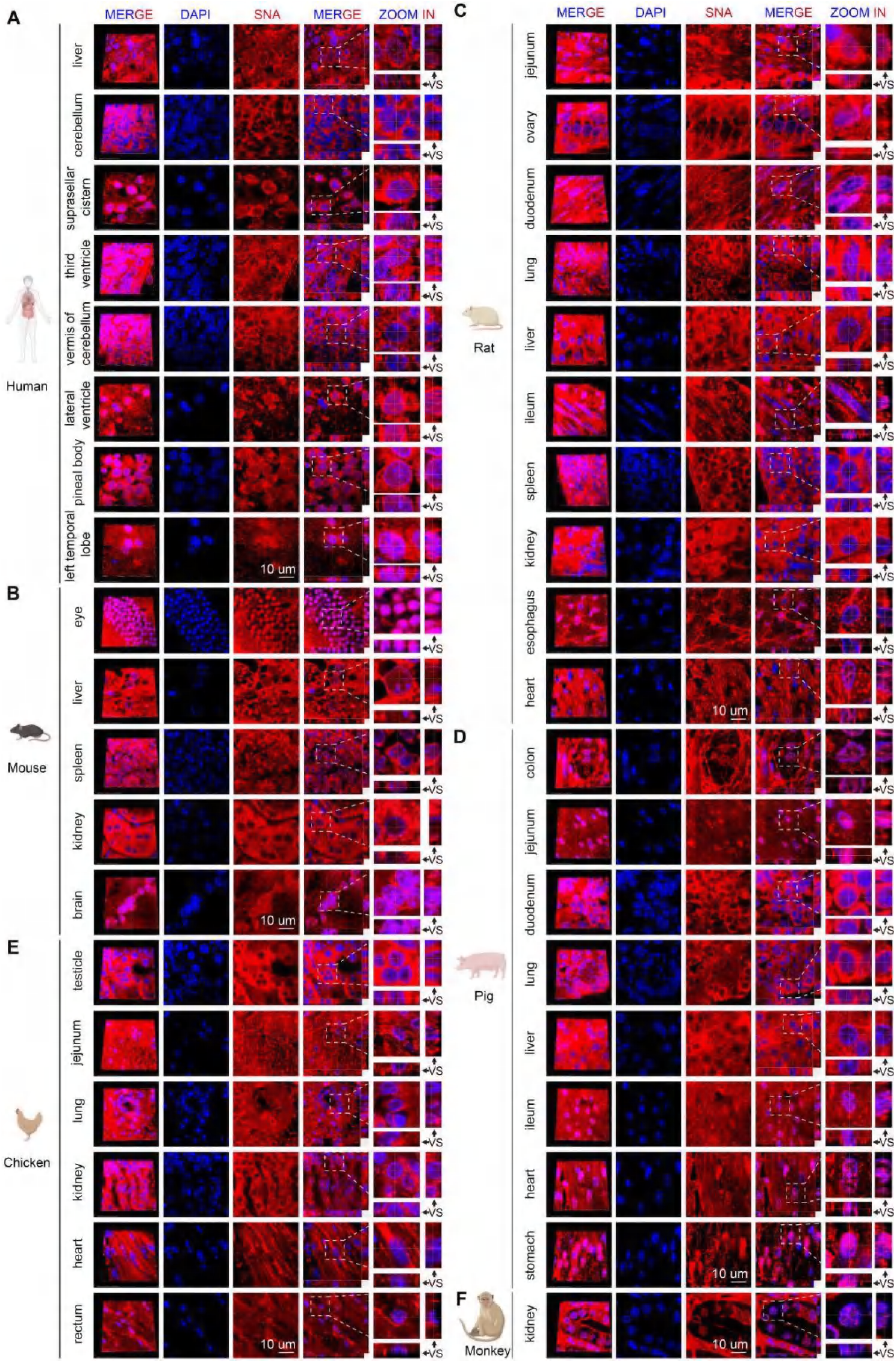
Nuclear localizations of SNA recognizing glycosylation were observed in multiple species including mouse, rat, pig, chicken, and monkey (related to Figure 2) **(A-F)** Fluorescence images of tissue sections from human (**A**), mouse (**B**), rat (**C**), pig (**D**), chicken (**E**), and monkey (**F**) stained with SNA (red) show clear nuclear glycan signals across species. Nuclei are counterstained with DAPI (blue). Images are representative of three independent experiments. VS, vertical section; scale bars, 10 μm.

**Figure S16.**
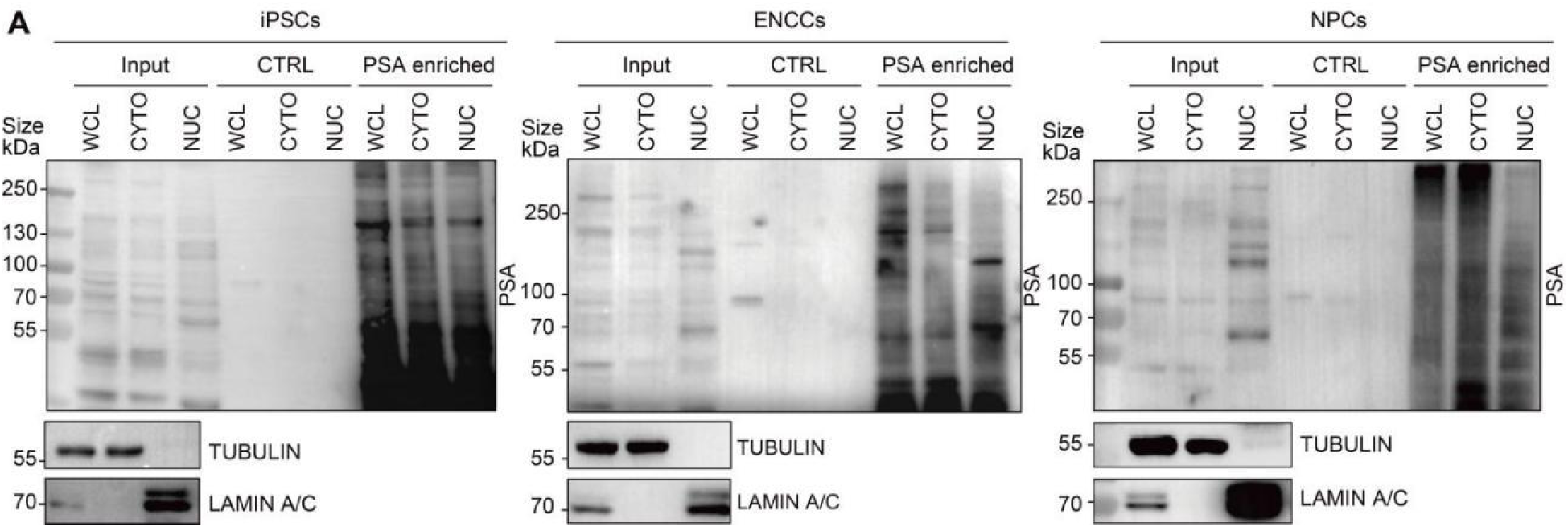
Abundance of core-fucosylated nuclear proteins in pluripotent stem cells and neural lineage cells (related to Figure 4) **(A)** PSA lectin enrichment of glycoproteins from whole-cell lysates, cytoplasmic, and nuclear fractions in various cell lines (iPSCs, ENCCs, NPCs). TUBULIN and LAMIN A/C serve as cytoplasmic and nuclear markers, respectively. Input lanes represent 1% of total protein; buffer-only controls are included.

**Figure S17.**
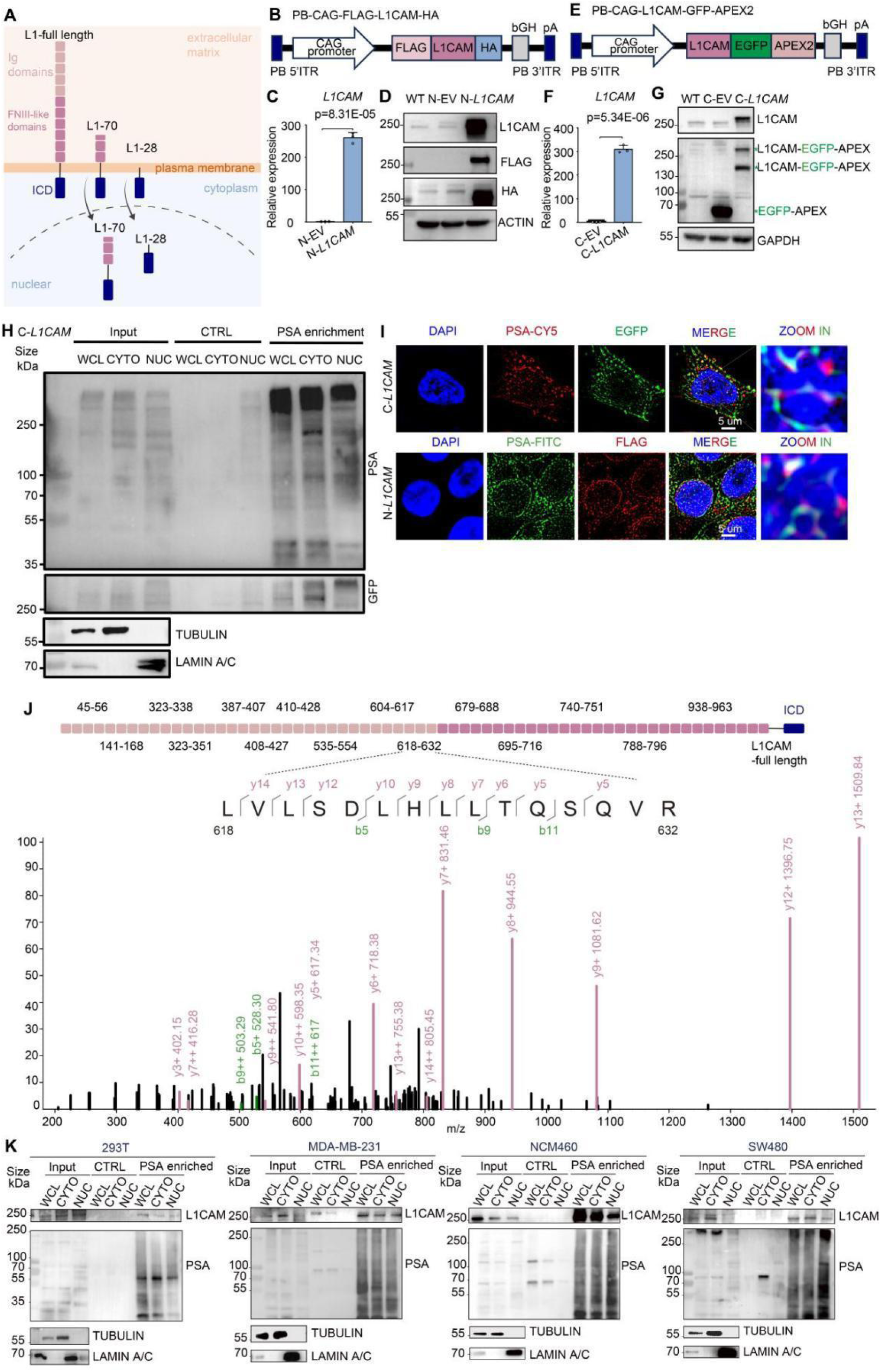
Nuclear localization of full-length core-fucosylated L1CAM (related to Figure 4. **(A)** Schematic illustrating the full-length structure of L1CAM (L1-250), comprising six Ig-like domains, five FNIII repeats, and transmembrane/intracellular domains. Previous reports documented nuclear localization of truncated L1CAM fragments (L1-28, L1-70). **(B)** Diagram of the piggyBac constructs encoding N-terminal FLAG-tagged *L1CAM*(N-*L1CAM*; PB-CAG-FLAG-*L1CAM*-HA). **(C** and **D)** RT-qPCR (**C**) and western blot (**D**) confirming overexpression of *L1CAM*in ENCCs transfected with N-*L1CAM*. Statistical significance assessed by t-test (two tailed). (**E**) Diagram of the piggyBac constructs encoding C-terminal EGFP-APEX2-tagged *L1CAM*(C-*L1CAM*; PB-CAG-*L1CAM*-EGFP-APEX2). (**F** and **G**) RT-qPCR (**F**) and western blot (**G**) analyses confirming *L1CAM* overexpression ENCCs transfected with C-*L1CAM*. Statistical significance assessed by t-test (two tailed). **(H)** Co-IP assay demonstrating that exogenous full-length L1CAM (EGFP-APEX2-tagged) localizes to the nucleus and is core-fucosylated. Nuclear fractions from C-*L1CAM*ENCCs were first immunoprecipitated for L1CAM and then enriched with PSA lectin to isolate core-fucosylated L1CAM. TUBULIN and LAMIN A/C were used as cytoplasmic and nuclear markers, respectively. IgG and buffer-only samples served as negative controls. Input corresponds to 1% of total protein used. Blot is representative of two independent experiments. **(I)** Fluorescence images showing nuclear co-localization of PSA lectin signal with overexpressed *L1CAM*in ENCCs. In N-*L1CAM*-expressing cells, PSA-FITC (green) co-localizes with FLAG-*L1CAM*(anti-FLAG immunostaining, red) in nuclei. In C-*L1CAM*-expressing cells, PSA-Cy5 (red) co-localizes with EGFP-tagged *L1CAM*(green) in nuclei. Nuclei are stained with DAPI (blue). Scale bars, 5 μm. Images are representative of three independent experiments. **(J)** Mass spectrometry (MS) chromatogram identifying non-contiguous peptides of L1CAM from aa 45 to 963, indicating the presence of L1CAM fragments longer than the known truncation site at residue 814 (consistent with full-length nuclear L1CAM). **(K)** PSA lectin enrichment of glycoproteins from whole-cell lysates, cytoplasmic, and nuclear fractions in various cell lines (293T, MDA-MB-231, NCM460, SW480). A prominent ∼250 kDa L1CAM band is detected in nuclear fractions of all tested lines. TUBULIN and LAMIN A/C serve as cytoplasmic and nuclear markers, respectively. Input lanes represent 1% of total protein; buffer-only controls are included.

**Figure S18.**
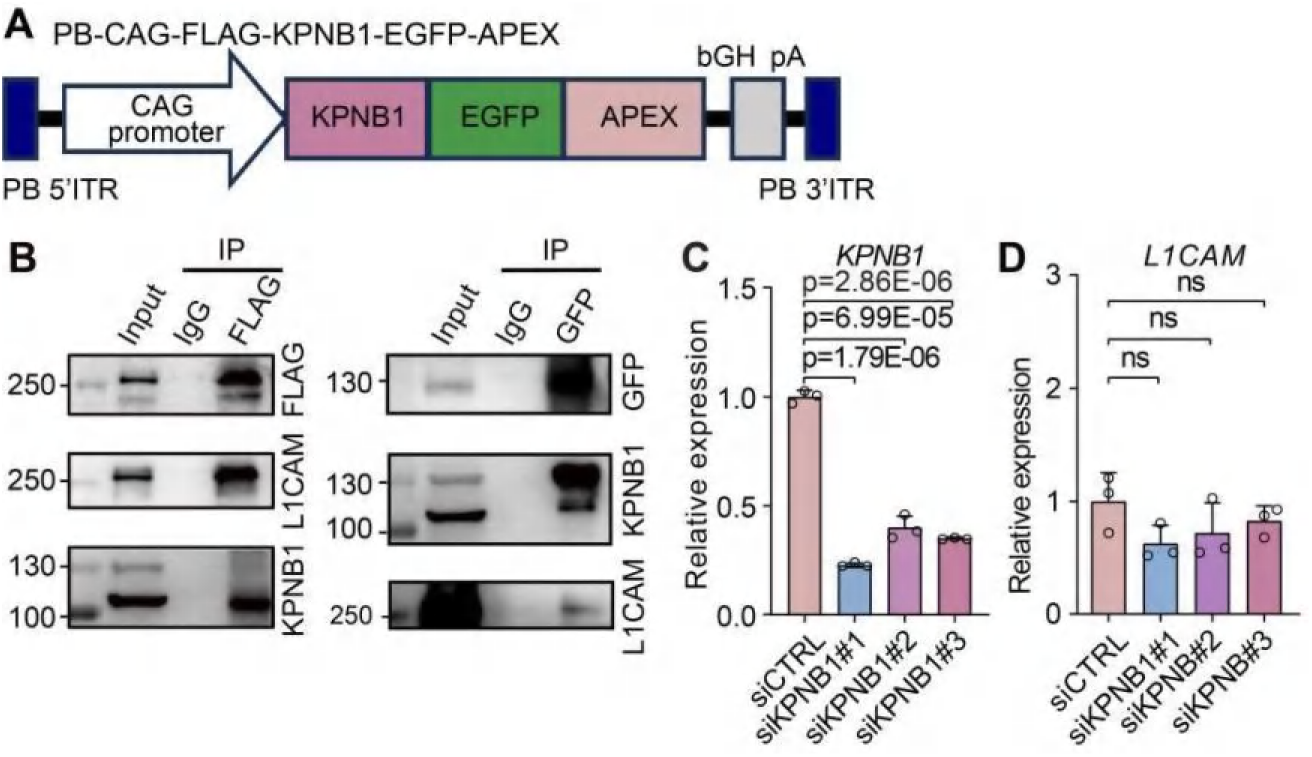
KPNB1 mediates nuclear import of full-length core-fucosylated L1CAM (related to Figure 4. **(A)** Schematic of the piggyBac construct encoding *KPNB1* (importin-β1) tagged with an N-terminal FLAG and C-terminal EGFP-APEX2 (PB-CAG-FLAG-*KPNB1*-EGFP-APEX2). **(B)** Co-IP demonstrating the interaction between full-length L1CAM and the nuclear import receptor KPNB1. ENCCs co-transfected with FLAG-*L1CAM*and EGFP-APEX2-*KPNB1* were subjected to immunoprecipitation for L1CAM, and co-precipitating KPNB1 was detected by western blot. IgG was used as a negative control. Input represents 1% of total protein. **(C** and **D)** RT-qPCR analyses showing that siRNA-mediated *KPNB1* knockdown in ENCCs significantly reduces *KPNB1*mRNA levels (**C**) and does not alter overall *L1CAM*mRNA levels (**D**). Statistical significance assessed by t-test (two tailed).

**Figure S19.**
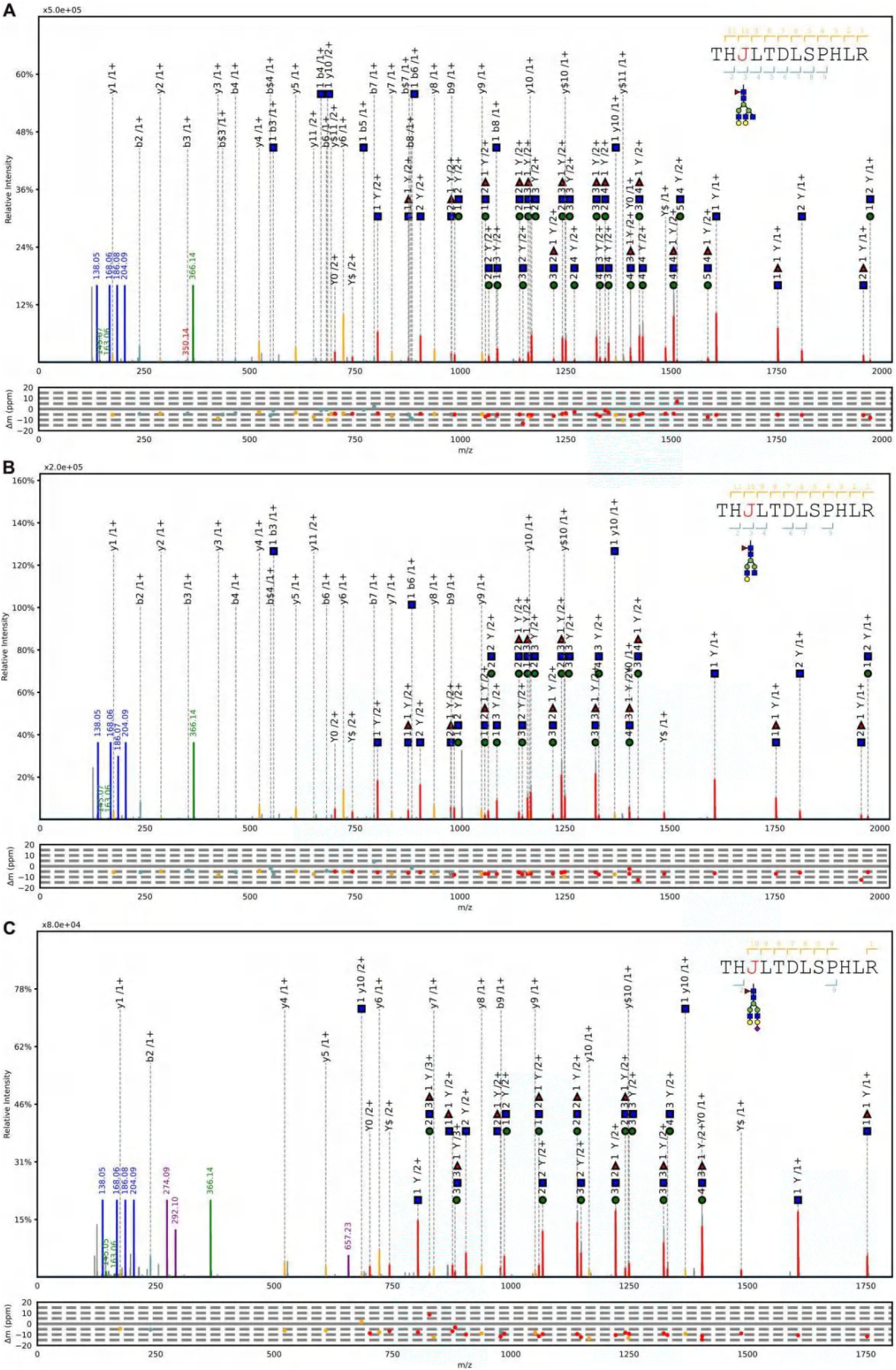
Tandem MS spectra of the L1CAM glycopeptide identified at site 979 (related to Figure 5) **(A-C)** Representative MS spectra are shown for site 979 with three different glycans compositing of H(5)N(5)F(1) (**A**), H(4)N(4)F(1) (**B**) and H(5)N(4)A(1)F(1) (**C**), respectively, confirming the presence of core-fucose modifications. The glycan symbols are green circle for Hex (H), blue square for HexNAc (N), red triangle for fucose (F) and purple diamond (A) for NeuAc. Peptide sequence with “J” indicating the N-glycosylation site. The LC-MS/MS data for glycopeptides were analyzed using pGlyco software (v3.0).

**Figure S20.**
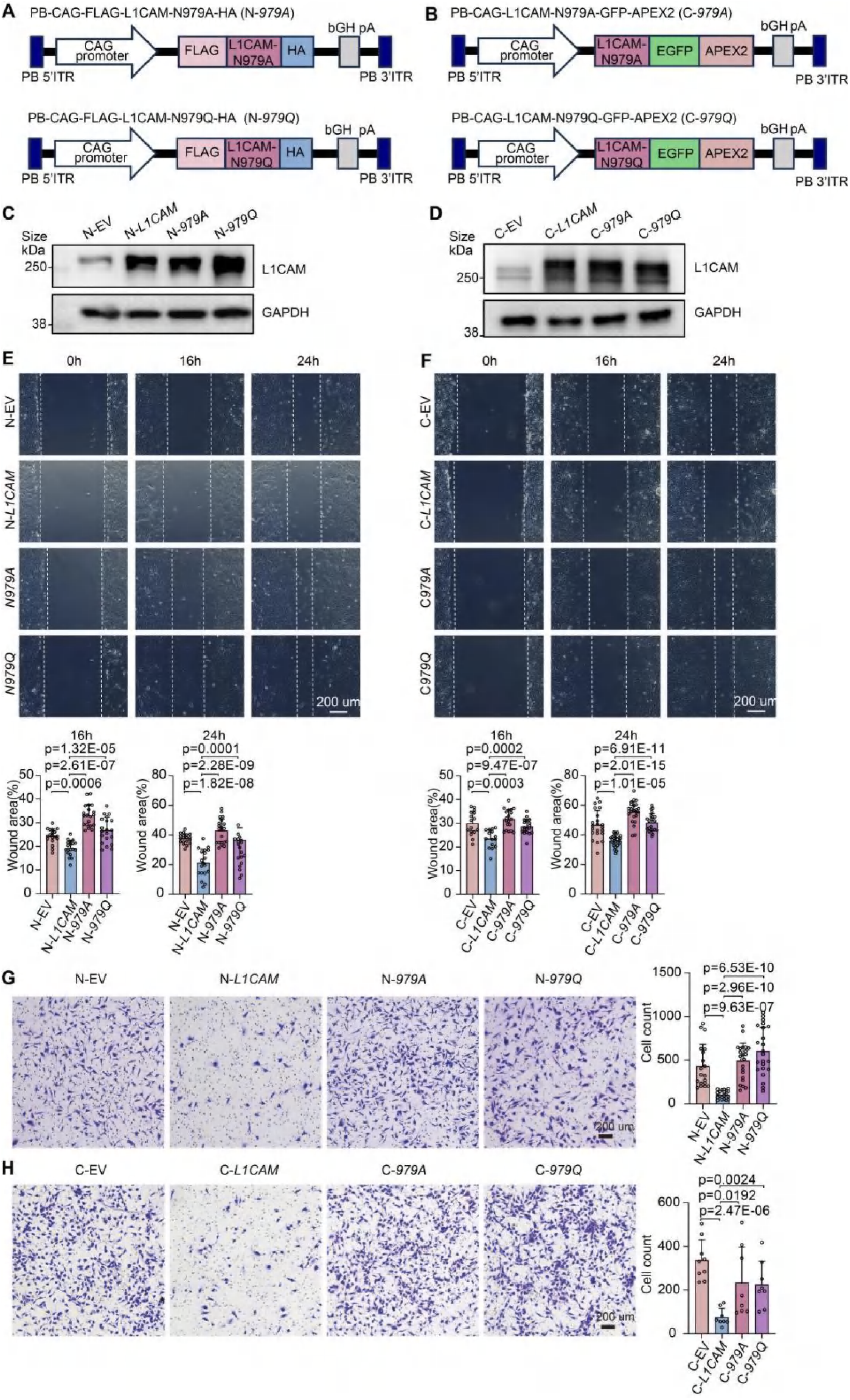
Wild-type L1CAM overexpression impairs ENCC migration compared to glycosylation-deficient mutants (related to Figure 5) **(A** and **B)** Schematic diagrams of L1CAM glycosylation-site mutants lacking core fucosylation at N979: N-terminal FLAG-tagged mutants N-*979A*and N-*979Q* **(A)**, and C-terminal EGFP-APEX2-tagged mutants C-*979A*and C-*979Q***(B)**. **(C** and **D)** Western blotting analysis confirming the expression levels of wild-type *L1CAM*(N-*L1CAM* and C-*L1CAM*), glycosylation-deficient mutants (N-*979A/Q*, C-*979A/Q*), and corresponding empty vector controls (N-EV, C-EV) in ENCCs. **(E** and **F)** Representative images of wound healing assays showing impaired migration of ENCCs overexpressing wild-type *L1CAM*(N-*L1CAM*, C-*L1CAM*) compared to glycosylation-deficient mutants (N-*979A/Q*, C-*979A/Q*) or empty vector controls (N-EV, C-EV). Statistical significance was assessed by t-test (two tailed). **(G** and **H)** Representative images of transwell migration assays confirming the reduced migration capacity of ENCCs overexpressing wild-type L1CAM (N-*L1CAM*, C-*L1CAM*) compared to glycosylation-deficient mutants (N-*979A/Q*, C-*979A/Q*) or empty vector controls (N-EV, C-EV). Statistical significance was assessed by t-test (two tailed).

**Figure S21.**
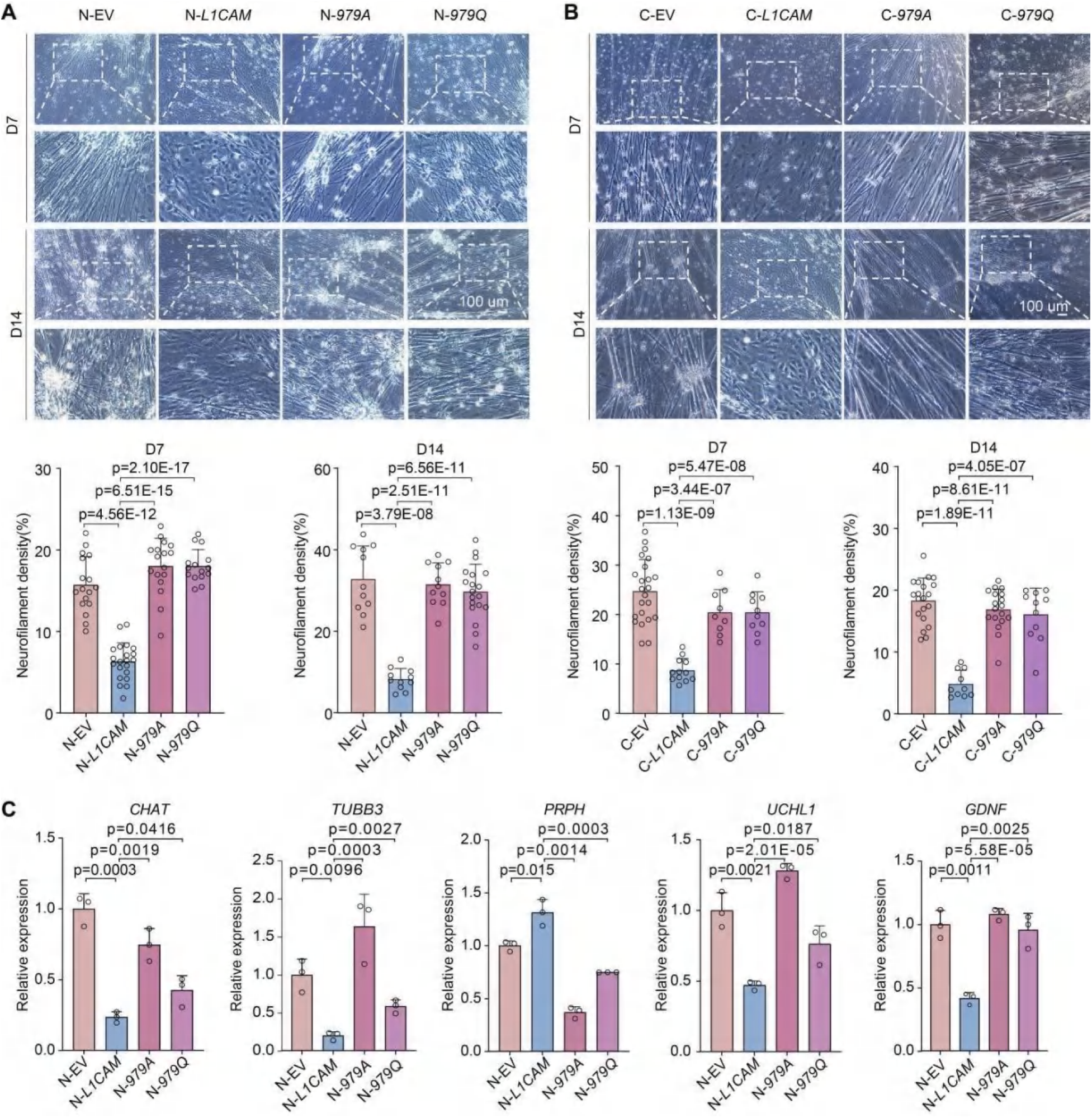
Wild-type L1CAM overexpression impairs neuronal differentiation compared to glycosylation-deficient mutants (related to Figure 5) **(A** and **B)** Brightfield microscopy images demonstrating that overexpression of wild-type *L1CAM*(N-*L1CAM*or C-*L1CAM*) markedly impairs differentiation of enteric neurons (ENCs), compared to glycosylation-deficient mutants (N-*979A/Q*, C-*979A/Q*) or empty vector controls (EV). Scale bar, 100 μm. Statistical significance was assessed by t-test (two tailed). **(C)** RT-qPCR confirming reduced expression levels of neuronal differentiation markers in enteric neurons (ENCs) overexpressing wild-type *L1CAM*compared to glycosylation-deficient mutants and controls. Statistical significance assessed by t-test (two tailed).

**Figure S22.**
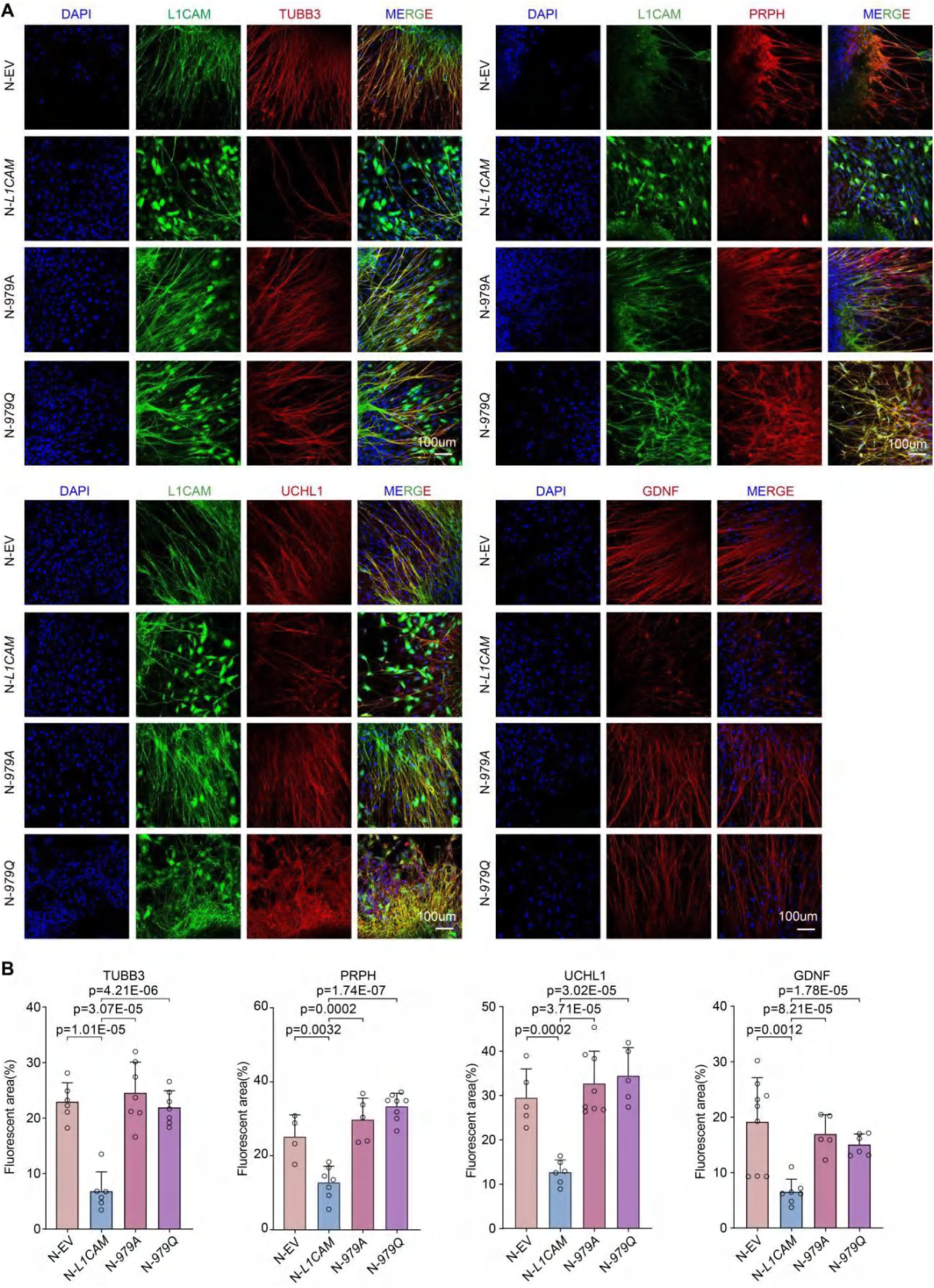
Wild-type *N-L1CAM* overexpression disrupts neuronal marker expression, unlike glycosylation-deficient mutants (related to Figure 5. **(A)** Immunofluorescence images of enteric neurons (ENCs) showing expression of neuronal differentiation markers (TUBB3, PRPH, UCHL1, GDNF; red) after overexpression of wild-type N-*L1CAM*versus glycosylation-deficient N-*979A*and N-*979Q*mutants. Nuclei are stained with DAPI (blue). Scale bar, 100 μm. Images are representative of three independent experiments. **(B)** Quantification of marker-positive cells (or fluorescence intensity) in the conditions from (**A**), demonstrating that wild-type N-*L1CAM* significantly reduces neuronal marker expression compared to N-*979A/Q*mutants (t-test (two tailed)).

**Figure S23.**
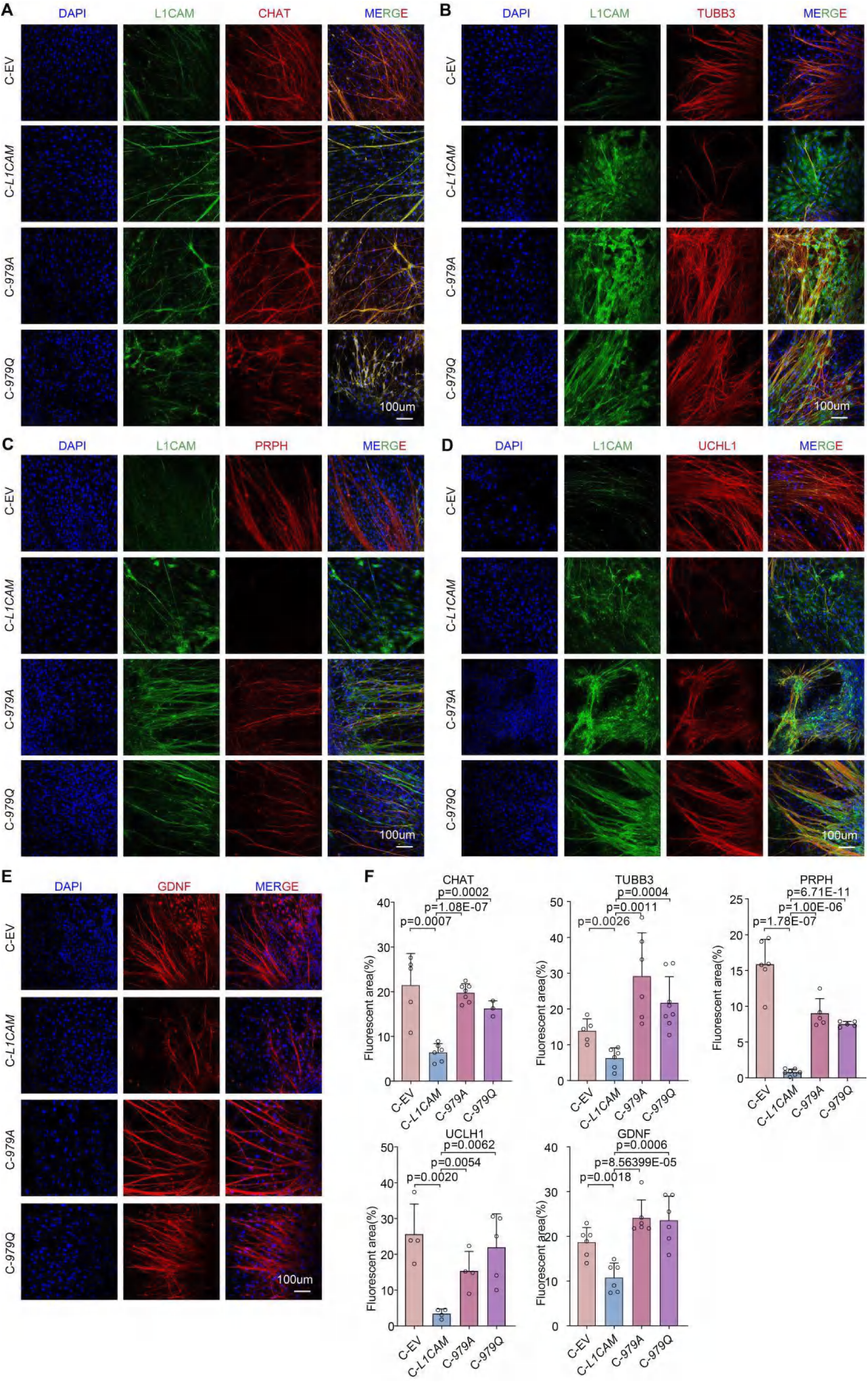
Wild-type C-*L1CAM* overexpression disrupts neuronal marker expression, unlike glycosylation-deficient mutants (related to Figure 5) **(A-E)** Immunofluorescence images of enteric neurons (ENCs) showing neuronal differentiation markers (red) after overexpression of wild-type C-*L1CAM*or glycosylation-deficient C-*979A*and C-*979Q*mutants. Evaluated markers include CHAT (**A**), TUBB3 (**B**), PRPH (**C**), UCHL1 (**D**), and GDNF (**E**), which are shown in red. Nuclei are stained with DAPI (blue). Scale bars, 5 μm. Representative images from three independent experiments. **(F)** Quantification of immunofluorescence staining shown in panels (**A**-**E**). Statistical significance was assessed by t-test (two tailed).

**Figure S24.**
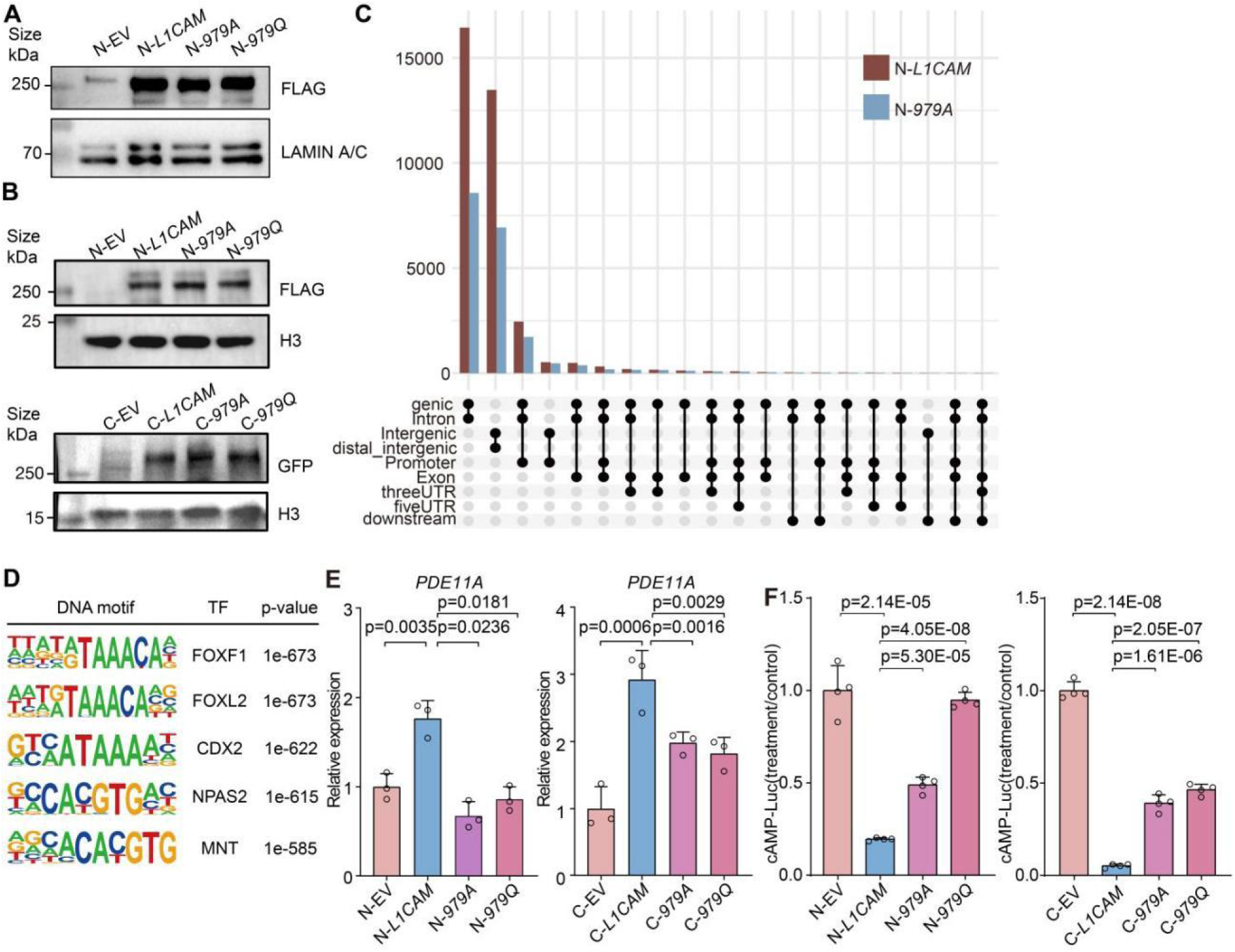
Core-fucosylation at N979 alters L1CAM chromatin binding and regulates PDE11A-cAMP signaling in differentiation (related to Figure 5. **(A)** Western blot of nuclear protein extracts from ENCCs expressing wild-type C-*L1CAM*, glycosylation-deficient C-*979A*and C-*979Q* mutants, or empty vector (C-EV). TUBULIN and LAMIN A/C serve as cytoplasmic and nuclear markers, respectively. Blot is representative of three independent experiments. **(B)** Western blot of chromatin-bound protein fractions from ENCCs expressing wild-type N-*L1CAM*and C-*L1CAM*, glycosylation-deficient mutants (N-*979A/Q*, C-*979A/Q*), or empty vector controls (N-EV, C-EV). Blot is representative of three independent experiments. **(C)** Genome-wide ChIP-seq profiles showing distinct chromatin occupancy for wild-type core-fucosylated L1CAM (N-*L1CAM*) versus the non-fucosylated mutant L1CAM (N-*979A*). Core-fucosylation at N979 enables unique binding at a subset of genomic loci. **(D)** DNA motif analysis of genomic loci co-bound by core-fucosylated L1CAM, revealing enrichment of specific DNA sequence motifs at these sites. **(E)** RT-qPCR analysis of *PDE11A*mRNA levels in ENCCs expressing empty vector, wild-type *L1CAM*constructs (N-*L1CAM*or C-*L1CAM*), or glycosylation-deficient L1CAM mutants (N-*979A/Q*, C-*979A/Q*). Statistical significance assessed by t-test (two tailed). **(F)** Quantification of intracellular cAMP levels in ENCCs expressing empty vector (N-EV), wild-type N-*L1CAM*, or glycosylation-deficient N-*979A/Q* mutants, using a cAMP-sensitive reporter (GloSensor-22F). Statistical significance determined by t-test (two tailed).

**Figure S25.**
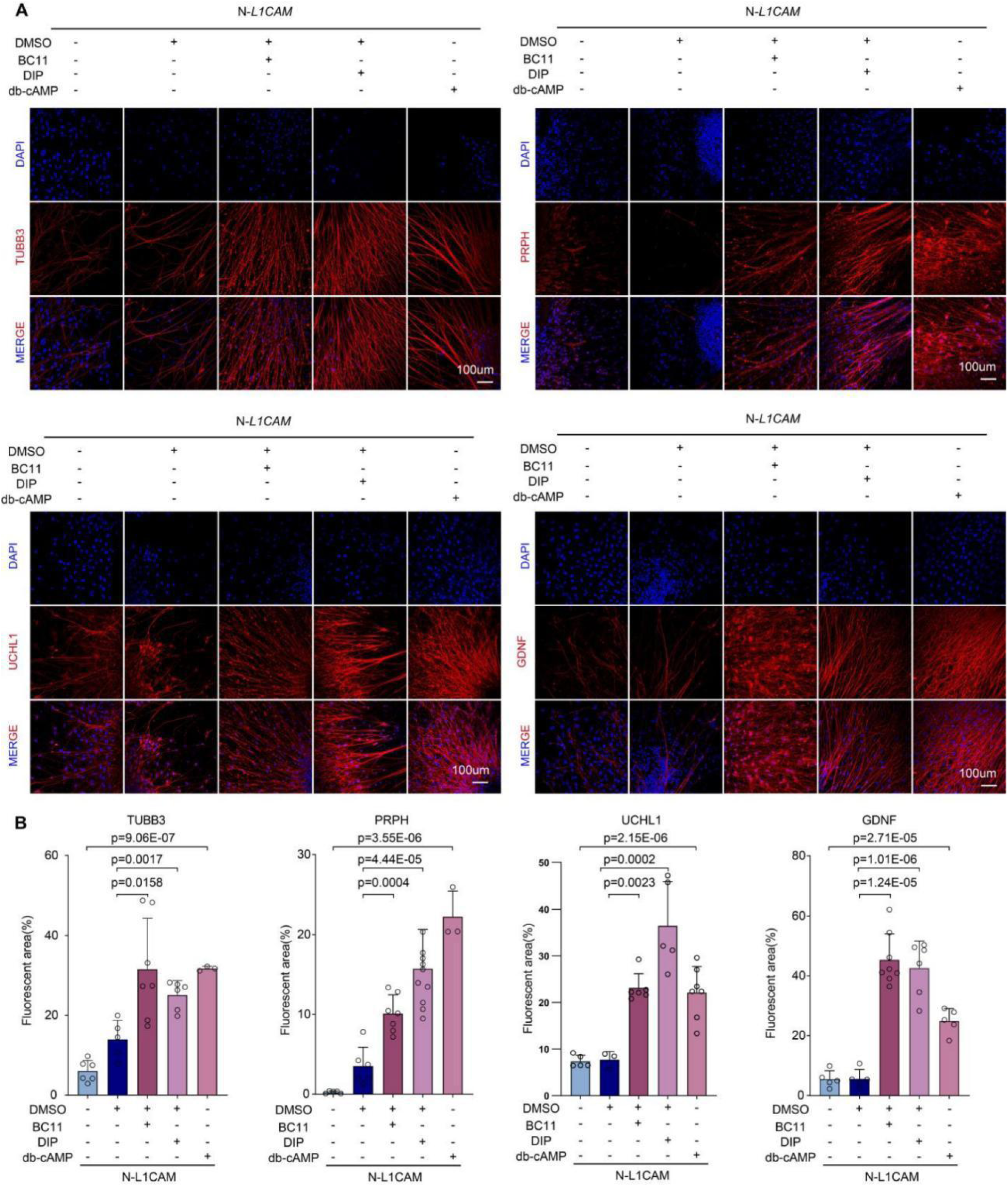
Pharmacological targeting of PDE11A-cAMP signaling rescues neuronal differentiation deficits induced by *L1CAM* overexpression in ENCs (related to Figure 5) **(A** and **B)** demonstrating restoration of neuronal differentiation marker expression (TUBB3; PRPH; UCHL1; GDNF; red) in N-*L1CAM*-overexpressing ENCCs following treatment with PDE11A inhibitors (DIP, BC11) or a cAMP analog (db-cAMP). Nuclei stained with DAPI (blue). Scale bar, 100 μm. Quantification was analyzed by t-test (two tailed) **(B)**.

**Figure S26.**
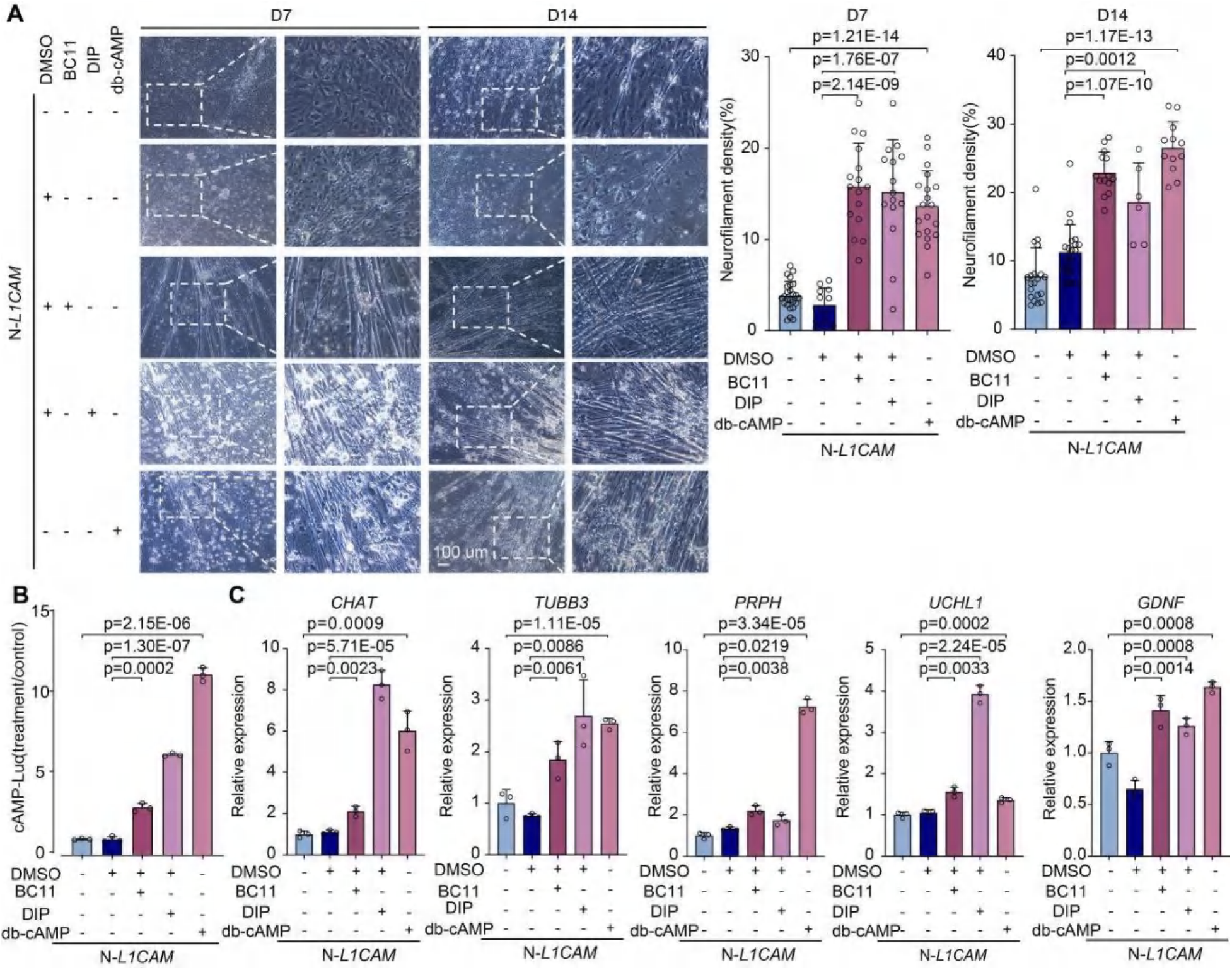
PDE11A-cAMP pathway restoration rescues neuronal differentiation in *L1CAM*-overexpressing ENCCs (related to Figure 5. **(A)** Bright-field images at day 7 (D7) and day 14 (D14) of differentiation, showing that ENCCs overexpressing N-*L1CAM*have impaired neuronal differentiation, which is reversed by treatment with PDE11A inhibitors (DIP, BC11) or a cAMP analog (db-cAMP). Scale bar, 100 μm. Quantification of differentiation outcomes (neurite outgrowth, neuron counts, or similar metrics) from at least three independent experiments shows significant improvement with each treatment (t-test, two tailed). **(B)** Intracellular cAMP levels in ENCCs stably expressing a cAMP-sensitive reporter (GloSensor-22F) with or without N-*L1CAM*overexpression, demonstrating that DIP, BC11, or db-cAMP treatment restores cAMP levels reduced by N-*L1CAM*(t-test, two tailed). **(C)** RT-qPCR analysis of neuronal differentiation marker expression in enteric neurons derived from the above conditions, showing that pharmacological activation of cAMP signaling rescues the expression of neuronal markers that were suppressed by N-*L1CAM* overexpression. Statistical significance assessed by t-test (two tailed).

**Figure S27.**
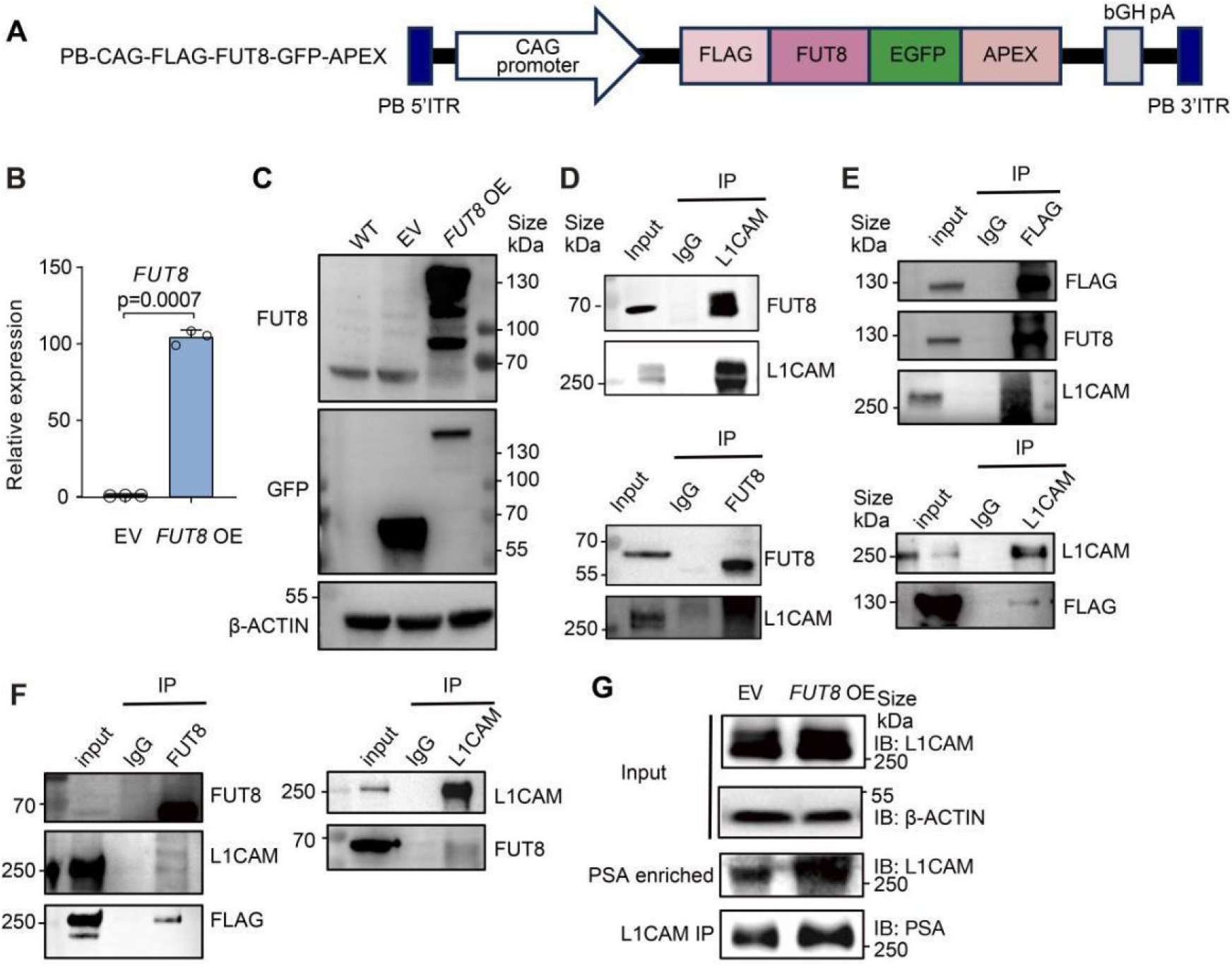
FUT8 interacts with L1CAM in ENCCs (related to Figure 6. (**A**) Schematic of the piggyBac *FUT8*overexpression construct tagged with FLAG and EGFP-APEX2 (PB-CAG-FLAG-FUT8-EGFP-APEX2). **(B** and **C)** RT-qPCR (**B**) and western blot (**C**) confirming FUT8 overexpression in transfected ENCCs. Statistical significance assessed by t-test (two tailed). **(D)** Co-IP showing the endogenous interaction between FUT8 and L1CAM in wild-type ENCCs (no overexpression). IgG was used as a negative control. Input corresponds to 1% of total protein. Result is representative of two independent experiments. **(E)** Co-IP showing the interaction between FUT8 and L1CAM in ENCCs overexpressing tagged *FUT8* (using the construct from (**A**)). IgG served as a negative control. Input is 1% of total protein. Representative of two independent experiments. **(F)** Co-IP showing the interaction between FUT8 and L1CAM in ENCCs overexpressing FLAG-tagged *L1CAM*(*L1CAM*OE; PB-CAG-FLAG-*L1CAM*-HA). IgG served as a negative control. Input is 1% of total protein. Representative of two independent experiments. **(G)** Co-IP demonstrating that core-fucosylated L1CAM levels are elevated in FUT8-overexpressing ENCCs compared to empty-vector controls. Representative of two independent experiments. Input represents 1% of total protein in each condition.

**Figure S28.**
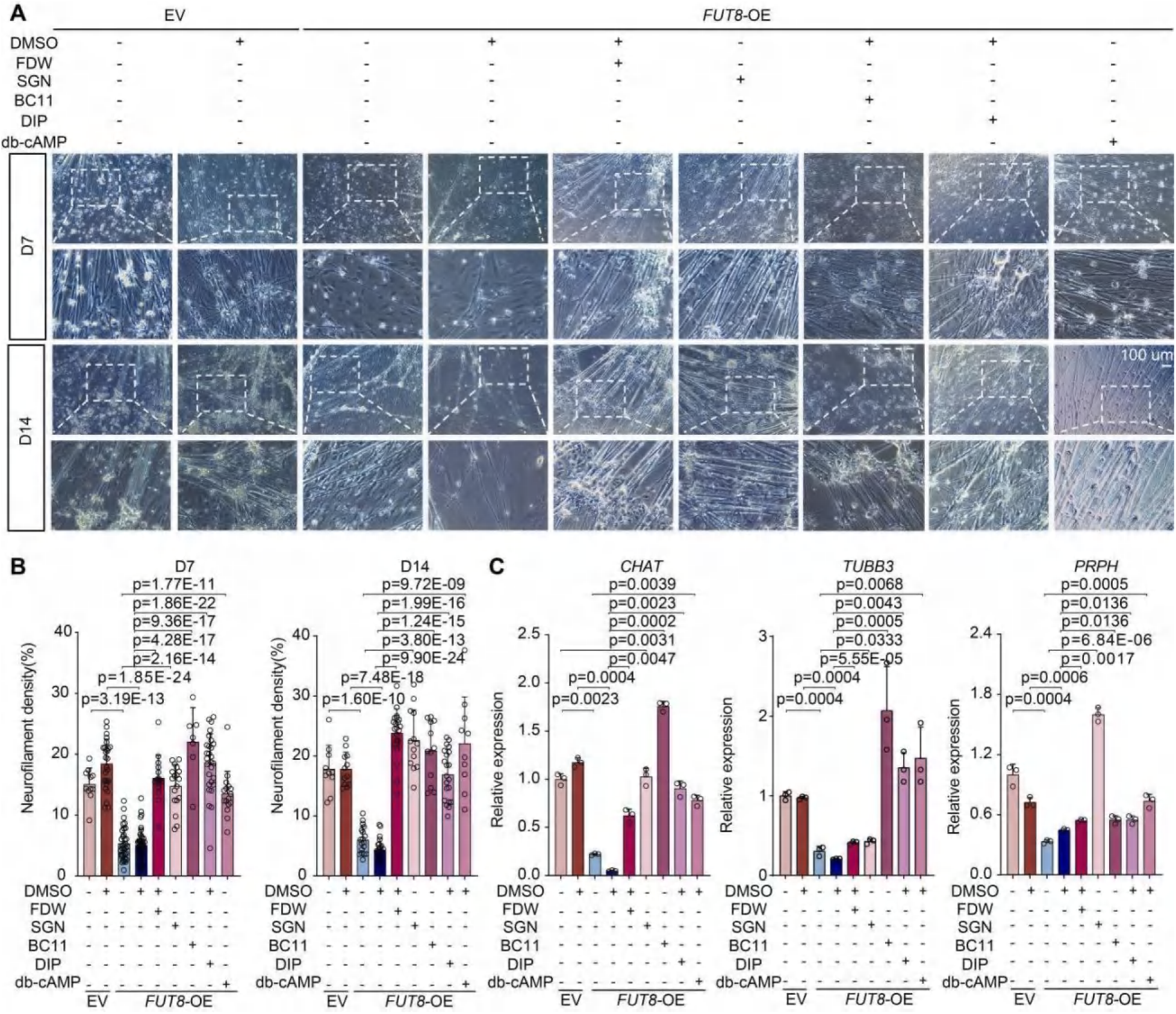
PDE11A-cAMP signaling modulation rescue of differentiation deficits caused by *FUT8* overexpression (related to Figure 6. **(A)** Bright-field images illustrating that *FUT8* overexpression (*FUT8* OE) in ENCCs impairs neuronal differentiation, and that treating these cells with PDE11A inhibitors (DIP, BC11) or a cAMP analog (db-cAMP) restores differentiation. DMSO treatment serves as a vehicle control. Scale bar, 100 μm. **(B)** Quantification of neuronal differentiation (e.g., proportion of cells expressing neuronal markers or forming neurons) corresponding to panel (**A**), showing significant rescue by DIP, BC11, or db-cAMP treatment (t-test, two tailed). **(C)** RT-qPCR analysis of neuronal differentiation marker expression (e.g., CHAT, TUBB3, peripherin) in FUT8-overexpressing ENCCs after treatment with DIP, BC11, or db-cAMP, compared to DMSO control. Data are mean ± s.e.m. from at least three independent experiments; statistical significance analyzed by t-test (two tailed).

**Figure S29.**
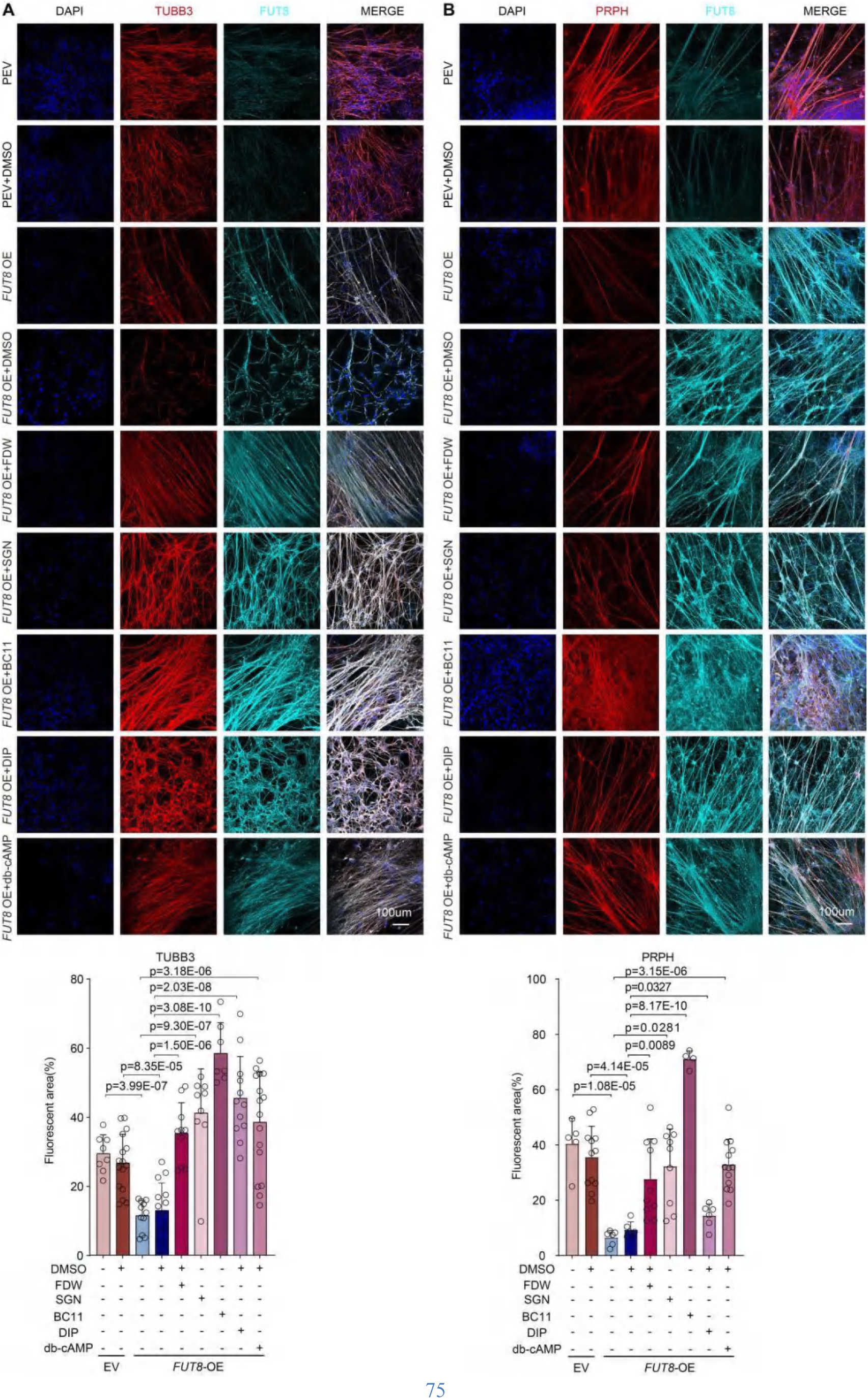
Immunofluorescence confirms rescue of FUT8-induced differentiation defects via cAMP signaling (related to Figure 6) **(A** and **B)** Immunofluorescence images showing that elevating cAMP signaling restores neuronal marker expression in *FUT8*-overexpressing ENCCs. ENCCs transduced with *FUT8*(*FUT8*OE) were treated with PDE11A inhibitors (DIP, BC11) or a cAMP analog (db-cAMP), resulting in recovery of neuronal differentiation markers TUBB3 (**A**) and PRPH (**B**) (red) that are otherwise reduced by *FUT8*OE. DMSO treatment serves as control. Nuclei are stained with DAPI (blue). Scale bar, 100 μm. Images are representative of three independent experiments. Statistical significance analyzed by t-test (two tailed).

**Figure S30.**
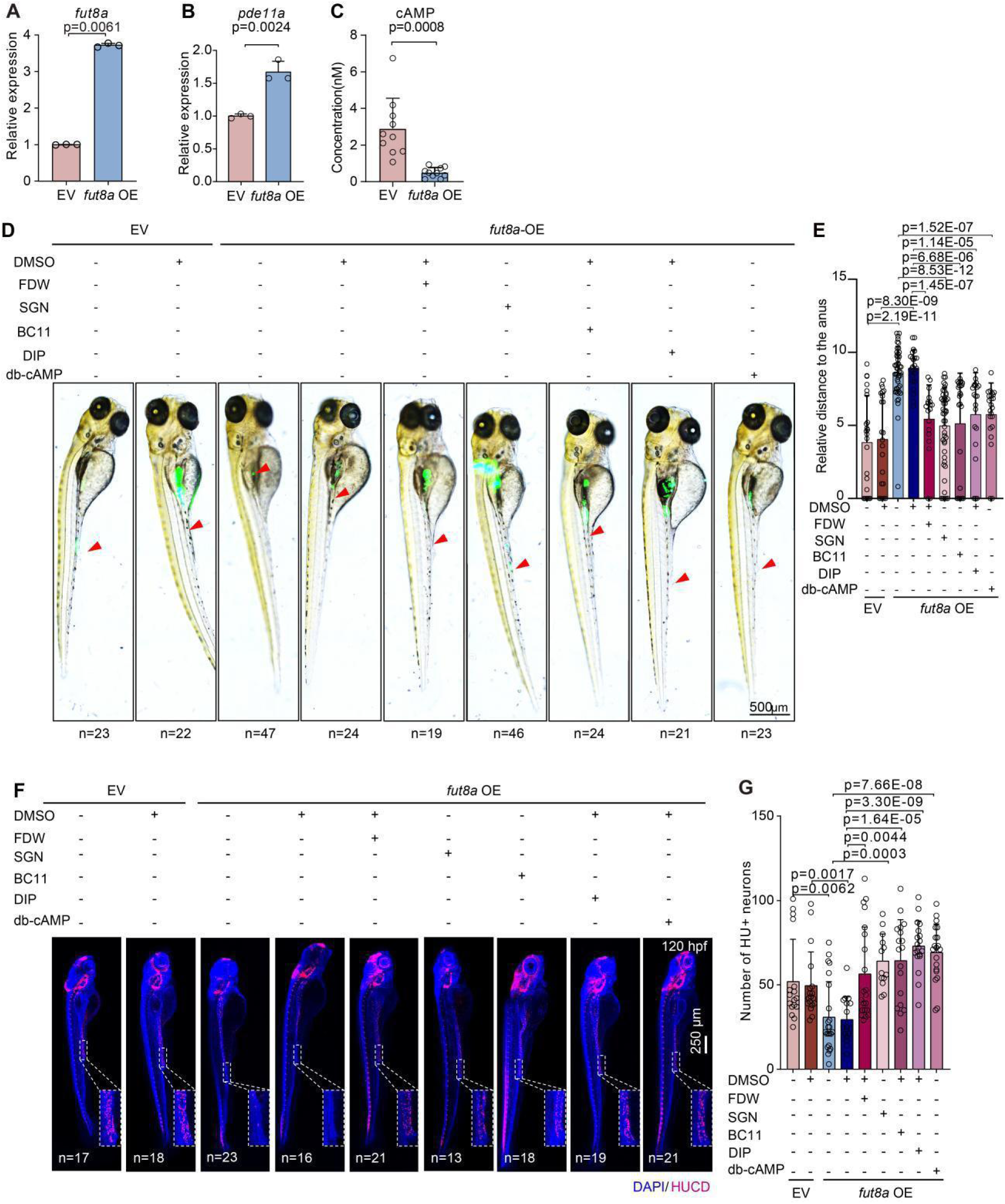
Modulating cAMP signaling rescues gastrointestinal motility defects caused by *fut8a* overexpression in zebrafish (related to Figure 6. **(A)** RT-qPCR confirming successful overexpression of *fut8a* in zebrafish embryos (relative to control). (**B and C**) Zebrafish model validation: RT-qPCR analysis of *pde11a*(**B**) and intracellular cAMP measurements (**C**) in *fut8a*-overexpressing (*fut8a*-OE) zebrafish compared to controls (EV). Statistical analysis via t-test (two tailed). **(D)** Representative bright-field images of 6-days-post-fertilization (6 dpf) zebrafish larvae showing reduced gastrointestinal motility (evidenced by gut peristalsis or transit) upon *fut8a* overexpression, and rescue of motility in larvae treated with PDE11A inhibitors (DIP, BC11) or a cAMP analog (db-cAMP). Scale bar, 500 μm. **(E)** Quantification of gastrointestinal motility in the conditions from (b), demonstrating that DIP, BC11, or db-cAMP treatment significantly improves motility in *fut8a*-overexpressing larvae. Statistical significance was determined by t-test (two tailed). (**F** and **G**) Immunofluorescence (HuC/D, red; DAPI, blue) assessing enteric neuronal differentiation defects induced by *fut8a* overexpression in zebrafish larvae, rescued by FUT8 inhibitors (FDW, SGN), PDE11A inhibitors (BC11, DIP), or cAMP supplementation (db-cAMP) (**F**). Scale bar, 250 μm. Quantitative analysis shown in panel, statistical significance via t-test (two tailed) (**G**).

**Figure S31.**
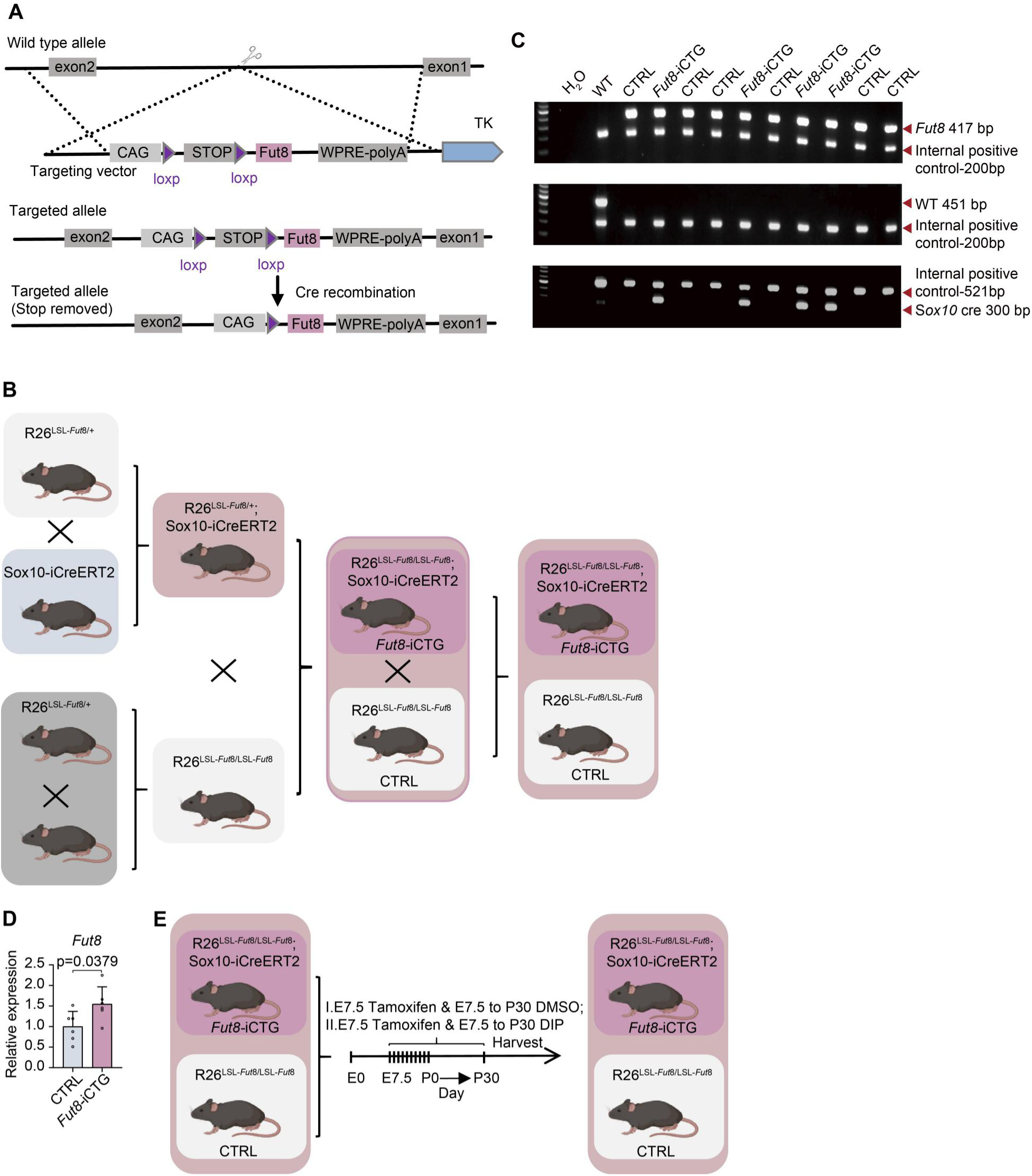
Generation of a induced transgenic mouse line with conditional overexpression of *Fut8* specifically in *Sox10* positive cells (*Fut8-*iCTG) (related to Figure 6. (**A** and **B**) Construction of transgenic mice. Induced *Sox10*-conditional *Fut8* transgenic mice (*Fut8**-***iCTG). Using CRISPR/Cas9 technology, we inserted the CAG-LSL-Fut8-WPRE-pA expression cassette into the Rosa26 locus via homologous recombination, resulting in LSL-*Fut8*/+ mice. To generate an induced transgenic mouse line with conditional overexpression of *Fut8* specifically in enteric neural crest lineage cells (*Sox10* positive cells), LSL-*Fut8*/+ mice were self-crossed to generate LSL-*Fut8*/LSL-*Fut8* mice, while they were also bred with *Sox10*-iCreERT2/+ mice to produce LSL-Fut8/+; *Sox10*-iCreERT2/+ offspring. Then LSL-*Fut8*/LSL-*Fut8* mice were crossed to LSL-Fut8/+; *Sox10*-iCreERT2/+ mice to yield LSL-*Fut8*/LSL-*Fut8*; *Sox10*-iCreERT2/+ progeny, as well as LSL-*Fut8*/LSL-*Fut8* mice. The line was established and maintained by crossing LSL-*Fut8*/LSL-*Fut8*; *Sox10*-iCreERT2/+ mice with LSL-*Fut8*/LSL-*Fut8* mice. Mice were treated with tamoxifen to induce *Fut8* overexpression in enteric neural crest lineage cells (*Sox10* positive cells) at E7.5 with 50ug/g tamoxifen. **(C)** The genotype identification of *Fut8-iTCG* mice with PCR detection. H_2_O, water; WT, wild type mouse; CTRL, control, LSL-*Fut8*/LSL-*Fut8 mouse*; *Fut8**-***iCTG, mouse with *Fut8* overexpression in *Sox10* positive cells, also labeled as *Fut8*-iCTG. **(D)** RT-qPCR confirming successful overexpression of *Fut8* in *Fut8-*iTCG mouse (relative to control, CTRL). **(E)** The schematic overview of the experiment to assess the effect of DIP treatment. mice were treated with tamoxifen (50ug/g) at E7.5 or with tamoxifen (50ug/g) at E7.5 and DIP (1ug/g) from E7.5 until after birth for 1month, or with tamoxifen (50ug/g) at E7.5 and DMSO (the same volume as DIP used) from E7.5 until after birth for 1month.

**Figure S32.**
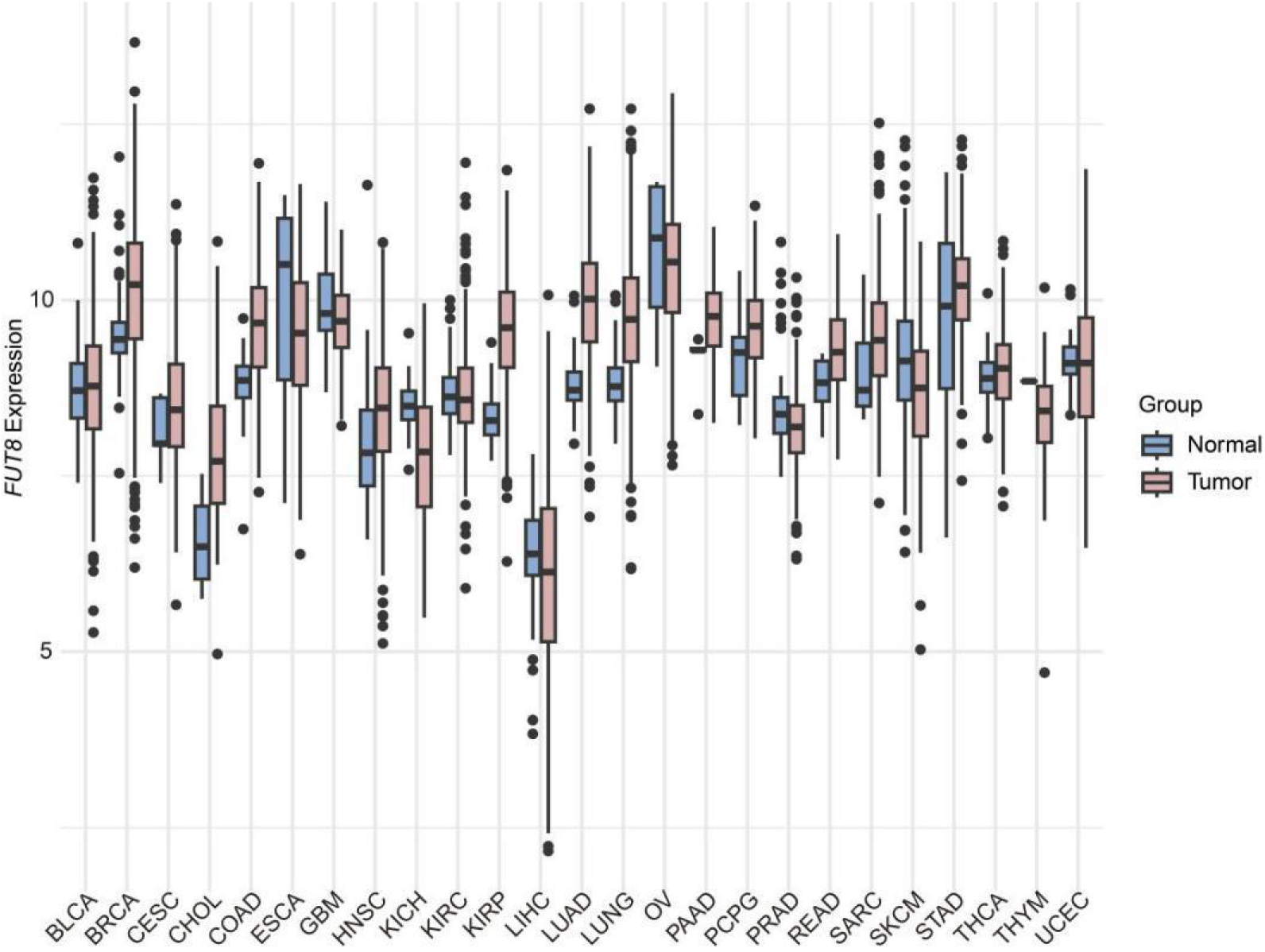
*FUT8* expression is upregulated in many human cancers (related to DISCUSSION) Gene expression analysis comparing *FUT8* levels in tumors versus matched normal tissues across 31 human cancer types (TCGA/GDC datasets, via the GEPIA portal). Each bar represents the median *FUT8*mRNA expression for a given tumor type or the corresponding normal tissue. Tumor types highlighted in red show significantly higher *FUT8*expression in tumors compared to their matched normal tissues.

## Supplementary tables

Supplementary tables are given as a separate Excel file.

**Table S1.**

Summary of PSA-interacting Proteins.

**Table S2.**

Summary of proteins identified in SILAC and quantification for L1CAM in membrane and nucleus.

**Table S3.**

Summary of L1CAM peptides identified in nuclear fraction.

**Table S4.**

Summary of L1CAM interacting proteins.

**Table S5.**

Glycoproteomics in nucleus.

**Table S6.**

Annotation of L1CAM peaks (N-L1CAM), L1CAM mutation peaks (N-979A) and of PHOX2B peaks.

**Table S7.**

Summary of the L1CAM (N-L1CAM)/L1CAM mutation(N-979A) RNA-seq.

**Table S8.**

GO analysis for L1CAM and PHOX2B targeted genes, which are also significantly differentially expressed in L1CAM (N-L1CAM)/L1CAM mutation(N-979A).

**Table S9.**

Sample information in Figure 6A and 6B.

**Table S10.**

Summary of RT-qPCR primers used in the research.

